# Light environment influences mating behaviours during the early stages of divergence in tropical butterflies

**DOI:** 10.1101/2020.07.28.224915

**Authors:** Alexander E. Hausmann, Chi-Yun Kuo, Marília Freire, Nicol Rueda-M, Mauricio Linares, Carolina Pardo-Diaz, Camilo Salazar, Richard M. Merrill

**Affiliations:** Division of Evolutionary Biology, Ludwig-Maximilians-Universität München, Grosshaderner Str. 2, 82152 Planegg-Martinsried, Germany; Smithsonian Tropical Research Institute, Gamboa, 0843-03092, Panama; Department of Evolutionary Neuroethology, Max Planck Institute for Chemical Ecology, Hans-Knöll-Straße 8, 07745 Jena, Germany; Department of Biology, Faculty of Natural Sciences, Universidad del Rosario, Carrera 24 No 63C-69, Bogotá 111221, Colombia

**Keywords:** *Heliconius*, Ecological Speciation, Sensory Environment, Magic traits, Hybrid Trait Speciation, Computer vision

## Abstract

Species divergence is facilitated when traits under divergent selection also act as mating cues. Fluctuations in sensory conditions can alter signal perception independently of adaptation to contrasting sensory environments, but how this fine scale variation affects behavioural isolation has received less attention, especially in terrestrial organisms. The warning patterns of *Heliconius* butterflies are under selection for aposematism and act as mating cues. Using computer vision, we extracted behavioural data from 1481 hours of video footage for 387 individuals. We show that the putative hybrid species *H. heurippa* and its close relative *H. timareta linaresi* differ in their response to divergent warning patterns, and that these differences are strengthened with increased local illuminance. Trials with live individuals reveal low-level assortative mating that are sufficiently explained by differences in visual attraction. Finally, results from hybrid butterflies are consistent with linkage between a major warning pattern gene and the corresponding behaviour, though the differences in behaviour we observe are unlikely to cause rapid reproductive isolation as predicted under a model of hybrid trait speciation. Overall, our results highlight that the role of ecological mating cues for behavioural isolation may depend on the immediate sensory conditions during which they are displayed to conspecifics.

## BACKGROUND

During ecological speciation, barriers to gene flow evolve as a result of ecologically-based divergent selection [1]. These barriers are generally expected to build up gradually over time [2], with premating isolation evolving early alongside ecological differences [1,3,4]. The evolution of premating isolation is facilitated when traits under ecological selection act as mating cues (sometimes known as ‘magic traits’ [5,6]), as this allows divergent natural selection acting on ecological traits to be transferred to mating behaviours. However, the strength of behavioural barriers may be influenced by the immediate, and perhaps rapidly changing, sensory conditions, but this has received relatively little empirical attention (but see [7–13]). One reason may be that robustly detecting these effects during the early stages of divergence, when they may be most relevant, presents a substantial empirical challenge.

It is well established that the sensory environment can alter signal detection and perception [14,15]. Colour perception depends not only on an object’s reflectance spectrum, but also the illumination, available light spectrum and background, all of which may change within seconds and over very short distances [16]. This can affect within-population preferences. For example, red colouration in male *Habronattus* spiders is only an efficient mating signal if presented in broad-spectrum sunlight [17]. Other sensory modalities can be also affected: Female swordtail fish are strongly attracted to the chemical signals of conspecific males in clean water, but not in polluted water [7]. Similarly, urban noise can disrupt the transmission of bird songs and interfere with acoustic-based mate choice [9].

The sensory environment may affect the evolution of reproductive isolation in two main ways. First, adaptations to meet the ecological needs of different sensory environments can influence female preferences, subsequently driving divergence in male signals, and leading to reproductive isolation through sensory drive [18]. For example, in two closely related *Pundamilia* cichlid fishes, adaptation of the visual system to the local environment is associated with divergent female mate preference for male colouration [19]. Second, prevailing environmental conditions may alter the efficacy of signals used in mate choice [14], so that individual mating preferences may act as an important reproductive barrier under some sensory conditions, but not others (‘context-sensitive preferences’ [20]). If the strength of preferences depends on the sensory environment, then this will influence their contribution to reproductive isolation. However, compared to the role of sensory adaptation, the immediate influence of local sensory conditions on mating behaviours, and how this may relate to the evolution and maintenance of new species, has been less-studied, especially in terrestrial organisms.

By affecting context-sensitive preferences, the sensory conditions during signalling may also influence the strength of genetic associations (*i.e*. linkage disequilibrium, LD) between mating and ecological traits, which are typically required for speciation to proceed when gene flow persists [21]. Specifically, when ecological traits act as mating cues, LD (between ecological and preference loci) will arise as a natural consequence of mating preferences, but this will be proportional to the strength of preference [22], which may be influenced by the sensory environment. Regardless, unless preferences are very strong, more robust coupling between mating and ecological components of reproductive isolation likely requires genetic architectures that impede recombination (or one-allele mechanisms, see [23]). These include physical linkage, which may be further strengthened by genomic rearrangements like inversions, or – in the extreme – pleiotropy, where distinct traits are controlled by the same allele [24]. To date, studies have reported physical linkage between behavioural and ecological components of reproductive isolation for a few animal taxa, including pea aphids [25], fish [26] and butterflies [27].

*Heliconius* butterflies are known for their bright warning patterns, which are often under selection for Müllerian mimicry [28]. Closely related *Heliconius* species frequently – but not always – have very different wing patterns, which additionally act as mating cues (e.g. [29–32]). This contributes to assortative mating because males almost invariably prefer wing patterns resembling their own, and warning patterns in *Heliconius* are considered one of the best examples of ‘magic traits’ in nature [6,29,33]. Variation in warning pattern is largely controlled by a few genes; the genetics of the corresponding visual mate preference are less well known, though recent work implicates a handful of genes associated with neural signalling in tight linkage to the colour pattern gene *optix* [27,34].

There is substantial evidence that colour pattern alleles have introgressed between otherwise separately evolving *Heliconius* lineages (e.g. [35]). In particular, taxa within the *heurippa-timareta* group, found in the eastern slopes of the South-American Andes, have acquired red colour pattern elements from local races of *H. melpomene* via introgression of *optix* alleles [36]. This has frequently led to near perfect mimicry between local races of *H. timareta* and *H. melpomene*; elsewhere, however, the resulting colour patterns are not shared with local *Heliconius* species. In particular, *H. heurippa* has a unique red-yellow banded forewing pattern (Fig. 1A). Its close relative *H. timareta linaresi*, which is assumed to represent the ancestral wing colour pattern of the *heurippa-timareta* group [37], only displays a yellow band (Fig. 1A) and also does not have an obvious co-mimic. These two populations are geographically adjacent (Fig. 1A) and presumably share a contact zone. Despite their nominal status as separate species, *H. t. linaresi* and *H. heurippa* likely represent an early stage of divergence; hybrids between *H. t. linaresi* and *H. heurippa* are fully viable and fertile [38] and any post-mating isolation is probably limited to selection acting against immigrant warning patterns.

**Figure 1:**
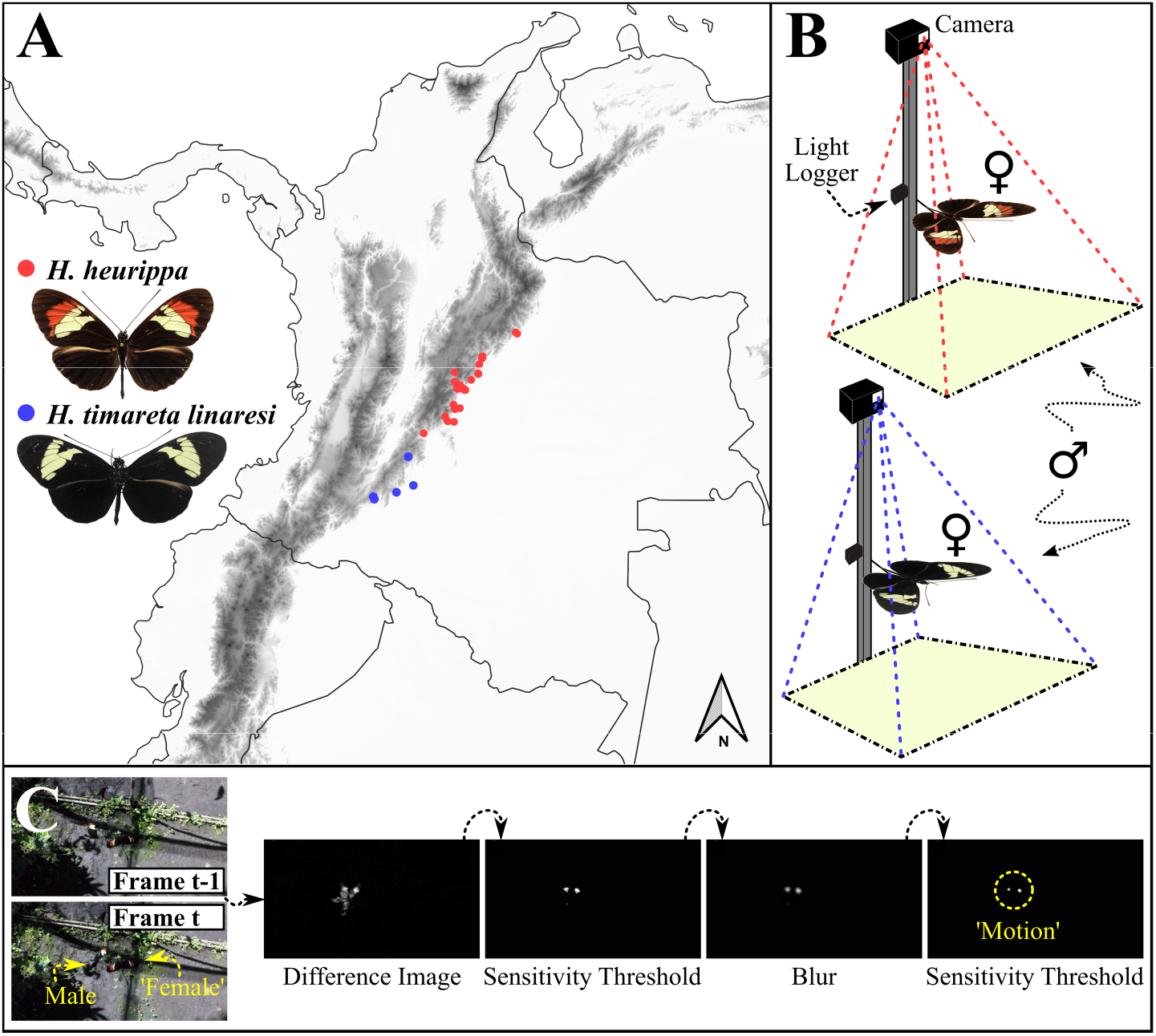
A) Locations in Colombia where *H. heurippa* (red) and *H. timareta linaresi* (blue) are known to occur. B) Video recording setup. Mounted females were presented simultaneously: Each was filmed and an associated light logger recorded illuminance every second. ‘Tripod’ was not casting shade on light logger or mounted females at any time. C) *Post-hoc* motion detection pipeline. A difference image is formed between each frame and its predecessor (identifying pixels with a change in value). Thresholding, blurring and again thresholding pick out significant local changes (‘signals’). ‘Signal’ frames (and surrounding frames) were extracted.

The red pattern element of *H. heurippa* is presumably maintained by strong frequency-dependent selection imposed by predators [39]. However, its effectiveness as intraspecific signal may depend on how it is perceived by conspecifics under natural sensory conditions. In this way, premating reproductive barriers may depend on interactions between the signal, environment and receiver. *H. melpomene* are broadly separated from *H. heurippa-timareta* (along with closely related *H. cydno*) taxa across a gradient of open to closed forest and decreasing light intensity [32,40]. These contrasting sensory environments are predicted to alter how colour patterns are perceived by butterfly visual systems [41]. While *H. heurippa* and *H. t. linaresi* are not ecologically isolated in this way, their forest habitats are highly heterogenous and *Heliconius* are known to settle with their wings open in more brightly illuminated ‘light patches’ [42]. Whether changes in illuminance affect male behaviours has not been investigated, but such an effect would suggest that the efficacy of wing patterns as premating reproductive barriers may depend on fluctuations in sensory conditions as females move through the environment.

To test whether warning patterns contribute to premating isolation between *H. heurippa* and *H. t. linaresi*, we used a novel computer vision pipeline to extract behavioural data from video footage, alongside choice trails to determine levels of assortative mating. Our data, including ~17,000 behavioural interactions for 387 individuals, allowed us to test (i) whether the two species show differences in visual attraction towards con- and hetero-specific patterns and (ii) whether these differences segregate with the colour patterns, consistent with physical linkage between the behavioural and colour patterning genes. Measuring illuminance in real-time at each female pattern also allowed us to ask (iii) whether these behavioural differences are influenced by fluctuating local light conditions.

## MATERIAL AND METHODS

### (a) Butterfly collection, maintenance and crossing design

We maintained outbred stocks of *H. timareta linaresi* and *H. heurippa*, established from wild individuals (Table S1), between January 2016 and September 2019 at the Universidad del Rosario insectaries in La Vega, Colombia (4°59’ N, 74°20’ W, elevation 1257 m). We generated F1 hybrids and backcross hybrids to each species (BL, backcross to *H. t. linaresi* and BH, backcross to *H. heurippa*) (Table S2). All butterflies were supplied with ~10% sugar solution and *Lantana, Psiguria* and/or *Gurania spp*.. Females were kept individually, with *Passiflora* for oviposition, whereas males were kept in groups. Eggs were collected every other day and the caterpillars were raised until eclosion in plastic cups, and fed on fresh *Passiflora* leaves.

### (b) Trials with mounted females

To test for differences in visual mating behaviours, we assayed males in choice trials with dead mounted *H. heurippa* and *H. t. linaresi* females presented simultaneously in an exposed 4×4×2m insectary, in which light conditions varied due to changes in cloud coverage and patterns of shade cast by vegetation at certain daytimes. Virgin females were frozen with their wings spread on the day of eclosion and kept at −20°C for >168h. They were then dried and subsequently washed in hexane to remove residual cuticular hydrocarbons and other pheromones, and mounted onto a small piece of wire. Throughout the experiment, we used 21 *H. heurippa* and 23 *H. t. linaresi* mounted females, randomly choosing a pair each day. Mounted females were individually attached to a horizontal wire (~70cm above ground) at one of six locations within the experimental cage. This location was changed every hour (hereafter ‘trial’), and the new location was chosen randomly (but female types were never in the same location twice during the same day). A *GoPro Hero 5 Black*™ (GoPro, San Matteo, CA) camera (settings and equipment in Table S3) was installed at each position, and ~50cm above the respective mounted female (Fig. 1B). At same height as the mounted female at ~30cm distance, we attached a *HOBO™ UA-002-64* logger (sensor facing up) to measure illuminance [*lux*] every second (Fig. 1B). Cameras and light loggers were synced using *GoPro Quik*™ and *HOBOware*™ software, respectively, allowing us to match video frames with light measurements.

Most of the virgin naive males matured in one separate communal cage before being introduced into the experimental cage >4 days after eclosion, where they were tested in mixed groups (median group size = 22). Males were numbered and received a unique code of dots on the dorsal side of the wings, allowing identification from videos. Each male was tested over multiple days (median = 12d). We recorded an average of 3.01h of material on each of the 246 recording days; conducting behavioural trials across different seasons and at different daytimes allowed us to capture a variety of light conditions.

### (c) Computer vision and video analyses

We used a custom motion-detection pipeline, which *post-hoc* discarded video footage with no activity. The detection of frames with motion (‘signals’) was based on difference imaging and consecutive steps of blurring and thresholding, as implemented in the *OpenCV* library available for C++ (Fig. 1C). Not all of the frames of male motion sequences made it over the threshold, so we determined the ‘signal’-frames and the surrounding frames one sec before and after a signal in R [43]. Reduced videos were created with the *OpenCV* library in C++. (Code is accessible at: github.com/SpeciationBehaviour/visual_preference_heurippa_linaresi). Video material was processed on an *HP™* Desktop computer (i7-7700 CPU, 4 cores), at runtimes ~80% of video duration. All videos were processed under the same threshold (145) and blur (30) settings. Remaining footage was curated manually at 66.6% speed using the *MPCHC™player*. We recorded three behaviours: ‘approach’ (male is changing its flight direction towards the mounted female, resulting in a curve or circling motion), ‘courtship’ (sustained hovering above mounted female) and ‘sitting’ (male sits down on mounted female). Behaviour types were combined for subsequent analyses.

### (d) Tetrad experiments with live females

We performed ‘tetrad’ trials with virgin males and females to test for assortative mating. For each trial, sexually mature *H. heurippa* and *H. t. linaresi* males (one of each) were allowed to acclimatize for 15 mins in a 2×4×2m insectary, at which point *H. heurippa* and *H. t. linaresi* virgin females (one of each) were introduced. Once the first mating occurred, the experiment was stopped.

### (e) Statistical analyses

We measured illuminance at both mounted females for 83% of recorded behaviours. Illuminance, measured in lux, is the intensity of light falling onto a surface. We log10-transformed *lux* measures (log-illuminance) and scaled (and centred) each set of log-illuminance measurements, making the choice of logarithm base irrelevant for the analyses.

Analyses were conducted in R [43] (supplemental R Markdown and https://github.com/SpeciationBehaviour/visual_preference_heurippa_linaresi). Posteriors will be described with a 95% equal-tailed credible interval and the mean as point estimate. We analysed data from trials with mounted females using logistic regression with the *brms* package [44], an interface to the Bayesian software Stan [45]. Male behaviours directed towards the *H. t. linaresi* or *H. heurippa* mounted females were fitted as binary Bernoulli-distributed response variable (*i.e*. 0 and 1, respectively); estimates from the model can be understood as a proportion of interactions with the mounted *H. heurippa* female, with higher values indicating stronger relative attraction to the *H. heurippa* ‘female’ and lower values indicating stronger relative attraction to the *H. t. linaresi* ‘female’. Models initially included all possible nested variations of the fully saturated model explaining effects of 1) male type, 2) log-illuminance at the *H. heurippa* ‘female’, 3) log-illuminance at the *H. t. linaresi* ‘female’, and their interactions. Individual ID and trial were fitted as random effects.

To test for ‘species’ differences, we initially fitted models for the ‘pure’ males (male type = ‘*H. heurippa*’ or ‘*H. t. linaresi*’). Segregation of the red bar in BL hybrids (controlled by alleles at *optix* [46]) allowed us to test for linkage between colour and preference loci (see [27,47]) (here male type = ‘red’ (*Bb* genotype) or ‘no red’ (*bb* genotype) and brood was additionally fitted as a random effect). To determine which terms to retain [48], we calculated the widely applicable information criterion (WAIC) and the leave-one-out information criterion (LOOIC) for each set of models, using the *loo* package [49], and WAIC weights using the *brms* package [44]. Weakly informative priors (centred around the value for no preference) were set for coefficients corresponding to the different male types, which gave small prior probabilities for extreme values very close to 0 or 1. For all other coefficients, default (non-informative) priors were applied. We also fitted ‘categorical illuminance’ models adopting the best fitting model structure determined for each dataset, where values =<median were ‘poorly lit’, and values >median were ‘brightly lit’ (Fig. S1). Posteriors of the estimated marginal means (EMMs) were calculated using the *emmeans* package [50]. From this, we retrieved posteriors of contrasts and calculated the posterior probability (PP) that EMMs differ.

For the tetrad data, we fitted observed counts of each mating outcome (type of male and female involved) as Poisson-distributed response variable. We included the specific male-female combination of the mating outcome as fixed factor. Transforming the resulting model estimates into proportions effectively makes this a multinomial model [51]. Non-informative default priors were applied throughout. PPs were calculated with the *brms* package [44]. We compared the observed frequency of each mating combination to the predicted frequencies based on our measurements of visual preference from the mounted female experiment. Predictions were derived by multiplying the frequency that a male type was involved in *any* mating combination by its respective EMMs from the models fitted to the mounted female data. Predictions were based either on the overall EMM for each type, or the interaction term EMMs from the categorical model. Posterior distributions for each prediction were calculated using the *binom* package under the default prior [52].

## RESULTS

### (a) Species comparisons: i) Divergent visual attraction behaviours in *H. timareta linaresi* and *H. heurippa* males are dependent on the light environment

Over 1.5 years we collected 1481 hours of footage. Our computer vision pipeline reduced this to 66 hours requiring human curation (*i.e*. 4.5% of the total footage recorded), including 16,995 behavioural ‘interactions’ from 387 males (~43.9 per male). These data allowed us to determine the effects of male type and illuminance on relative visual attraction to the *H. heurippa* mounted ‘female’ (hereafter ‘preference’). The best fitting model for the pure *H. heurippa* and *H. t. linaresi* males retained male type, log-illuminance at the *H. heurippa* ‘female’ and their interaction (Table S4: model #1). Overall, our results suggest that the local light environment influences the strength of visual attraction.

Across the entire dataset, illuminance at the mounted *H. heurippa* female increased the difference in preference between the male types, as evidenced by the interaction between male type and log-illuminance at the *H. heurippa* ‘female’ (Fig. 2A & S3, PP simple slope *H. heurippa* > *H. t. linaresi* = 0.996). This was largely driven by an increase in log-illuminance at the *H. heurippa* ‘female’ leading to a stronger conspecific preference in *H. heurippa* males (PP simple slope > 0 = 0.993); there was only limited support for an effect on *H. t. linaresi* males (PP simple slope < 0 = 0.768). Overall, *H. heurippa* males showed a higher proportion of interactions with the *H. heurippa* pattern than *H. t. linaresi* males. Although supported with high credibility (PP relative visual attraction *H. heurippa* > *H. t. linaresi* = 0.969), this difference in preference was relatively small (0.07, CrI: 0.00 - 0.14) and characterised by considerable within-population variation (Fig. 2B). Nevertheless, this effect nearly doubled when the *H. heurippa* ‘female’ was in brighter light (0.13, CrI: 0.05 - 0.22; Fig 2D) and was absent when the *H. heurippa* ‘female’ was poorly lit (0.01, CrI: −0.07 - 0.09; Fig 2C).

**Figure 2:**
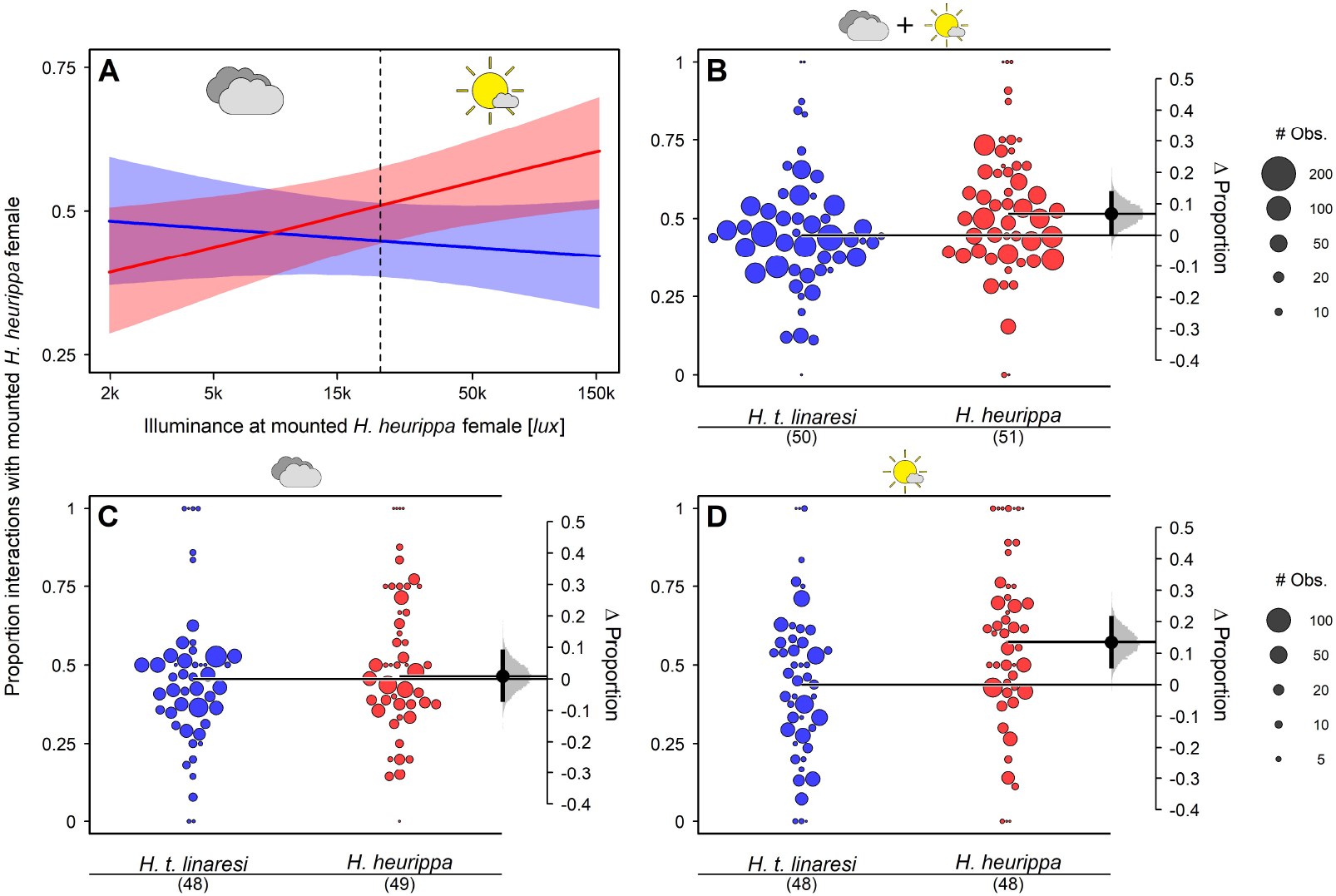
The effect of illuminance on relative attraction towards *H. heurippa* ‘females’ (*i.e*. proportion of interactions with the *H. heurippa* ‘female’ as opposed to the *H. t. linaresi* ‘female’). **A**: Relative attraction towards *H. heurippa* ‘females’ of *H. t. linaresi* (blue) and *H. heurippa* males (red) under changing illuminance at the *H. heurippa* ‘female’. Illuminance on the x-axis is log-scaled. Coloured area around each regression line represents the 95% credible interval (CrI). Dashed vertical line represents the median log-illuminance used as a cutoff to define the poorly and brightly lit conditions (below). Relative attraction towards *H. heurippa* ‘females’ for *H. t. linaresi* males (blue) and *H. heurippa* males (red): (**B**) across light environments; (**C**) for poorly lit *H. heurippa* ‘female’; (**D**) for brightly lit *H. heurippa* ‘female’. Gardner-Altman plots in B-D show the difference between the two male types: Horizontal lines project from the means of the posteriors for each male type (means and CrIs in Table S5). The mean and the 95% CrI for the posterior of the difference between the male types are shown on the right. Each point represents a single individual and its size is scaled to the number of observations. Custom swarmplot was used to distribute the dots horizontally.

### (b) Species comparisons: ii) *H. heurippa* males mate more often with conspecific females in choice trials

To test for assortative mating we also conducted ‘tetrad’ mate choice trials between *H. heurippa* and *H. t. linaresi*. During 89 tetrad trials we observed 50 con- and 39 heterospecific matings (PP for positive assortative mating = 0.88). This trend was driven by a higher proportion of conspecific matings involving *H. heurippa* males (0.405, CrI: 0.305 - 0.508) than heterospecific matings involving *H. heurippa* males (0.270, CrI: 0.183 - 0.366]) (PP = 0.94). *H. t. linaresi* males did not participate more frequently in con-rather than heterospecific matings (PP = 0.43) (Fig. 3). In general, *H. heurippa* males mated more often than *H. t. linaresi* males (60 vs. 29 times). Our results closely match predictions derived from the mounted females experiment with brightly lit *H. heurippa* ‘female’ (horizontal purple bars in Fig. 3), and, to some extent, those derived from all of the illuminance conditions combined (Table S6).

**Figure 3:**
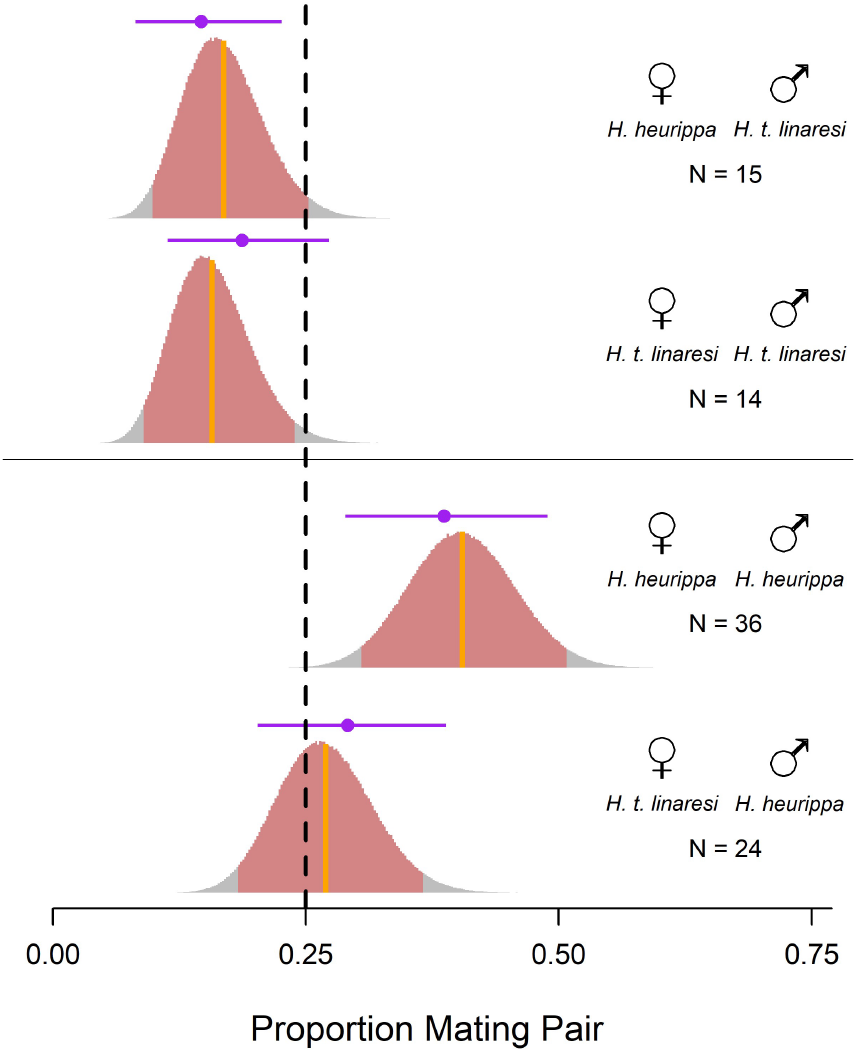
*H. heurippa* males show a preference for live, conspecific females in the tetrad experiments. Dashed vertical line indicates expected proportion under no assortative mating. Posterior distributions for each mating pair combination are displayed as histograms (red = 95% CrI, orange line = mean). Predictions based on the visual attraction data from the mounted females experiment with brightly lit *H. heurippa* ‘female’ are displayed with their 95% CrI as horizontal purple lines.

### (c) Hybrid comparisons: Patterns of behaviour in backcross hybrids are consistent with linkage between colour and preference loci

Estimates of preference suggest that first generation hybrid (F1) males behave like *H. heurippa* males (Fig. S4D), perhaps suggesting that *H. heurippa* alleles for attraction to red are dominant; however, challenging this, BH males seem to display a preference similar to *H. t. linaresi* males (Fig. S4E). For BL males, the model including all possible interactions was the best fit (Table S7, model #1). As for the ‘pure’ males, the six best fitting models included the interaction between type (*i.e*. red (Bb) and non-red (bb) wing pattern) and log-illuminance at the *H. heurippa* ‘female’ (91% of cumulative WAIC weight for an effect of this interaction term).

As for males from the parental populations, the difference between the two genotypes was higher when the *H. heurippa* ‘female’ was in bright light (Fig. 4CD). There was a small effect of the three-way interaction with illuminance measures at the *H. t. linaresi* ‘female’ - the difference was slightly higher when only the *H. heurippa* mounted female was brightly lit (Fig. 4C, 0.11, CrI: 0.02 - 0.21), as opposed to when both were (Fig. 4D, 0.07, CrI: 0.00 - 0.13). A brightly lit *H. t. linaresi* mounted female led to a slightly reversed pattern (i.e. *bb* > Bb), but only when the *H. heurippa* mounted female was poorly lit (Fig. 4B, −0.03, CrI: −0.13 - 0.07). If both mounted females where poorly lit, there was no difference in preference (Fig. 4A, 0.01, CrI: −0.05 - 0.08). Across the entire dataset, illuminance at the mounted *H. heurippa* female increased the differences in preference between the two BL genotypes (Fig. S2B, PP simple slope *Bb* > *bb* = 0.999). While the slope for *Bb* males is only slightly positive (PP = 0.859), the slope for *bb* males is strongly negative (PP = 0.999). In contrast to the pure males, we also observed an effect of illuminance at the *H. t. linaresi* ‘female’. Specifically, *bb* males showed a stronger preference for the *H. heurippa* ‘female’ when the *H. t. linaresi* ‘female’ was bright (Fig. S2D & S3, PP = 0.999). Overall, *Bb* males were more likely to interact with the *H. heurippa* ‘female’ than *bb* males (Fig. S5). This difference was supported with moderately high probability (PP relative visual attraction *Bb* > *bb* = 0.897). Although these represent small effects in absolute terms (0.03, CrI: −0.02 - 0.08]), this difference accounts for ~50% of the difference in means of the parental populations, consistent with linkage between colour and preference loci.

**Figure 4:**
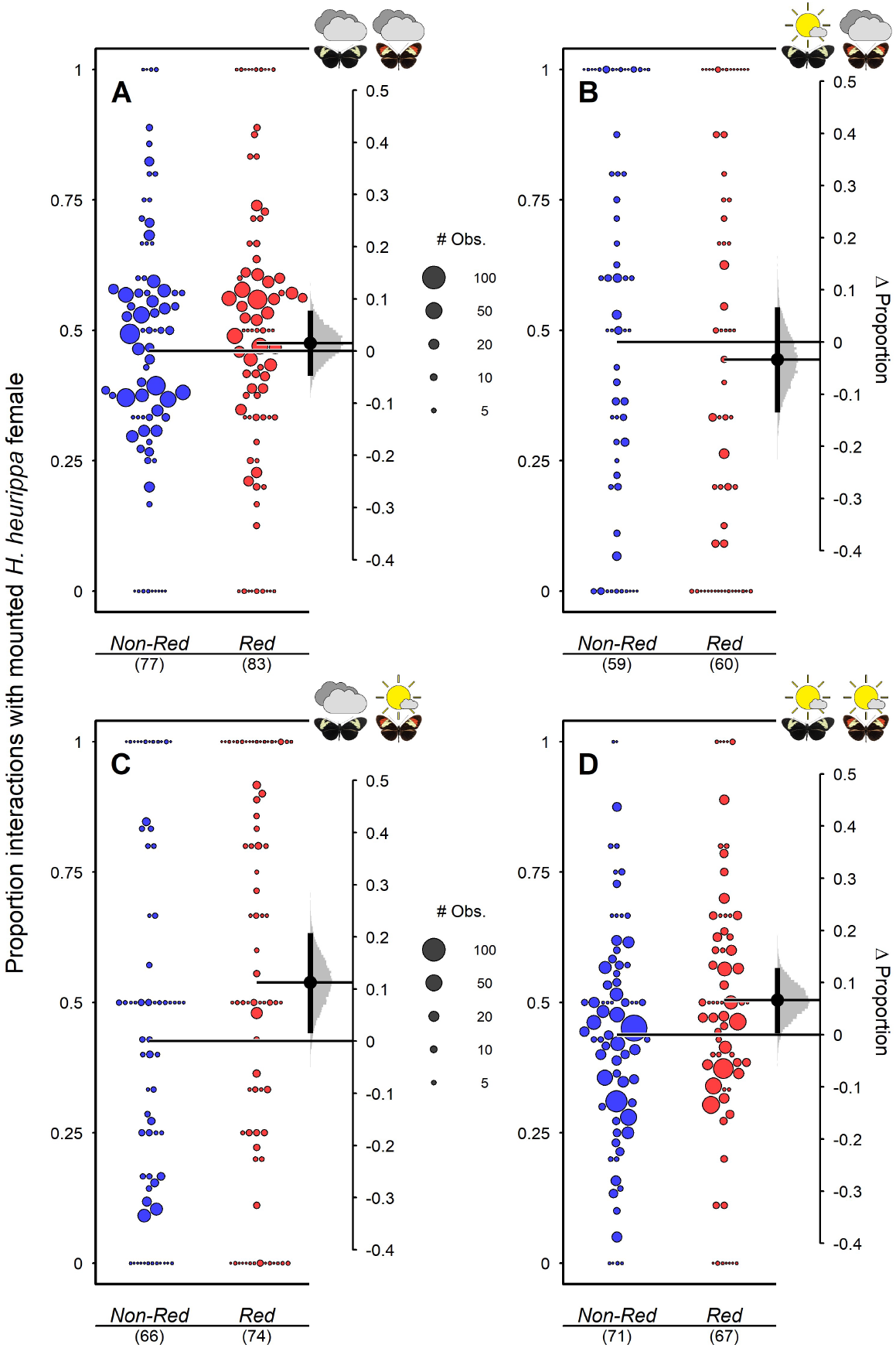
The effect of illuminance on relative attraction towards *H. heurippa* ‘females’ (i.e. the proportion of interactions with the *H. heurippa* ‘female’ as opposed to the *H. t. linaresi* ‘female’) for BL hybrid males (no red/bb = blue dots; red/*Bb* = red dots): (**A**) with poorly lit *H. t. linaresi* and *H. heurippa* ‘females’; (**B**) brightly lit *H. t. linaresi* and poorly lit *H. heurippa* ‘females’; (**C**) poorly lit *H. t. linaresi* and brightly lit *H. heurippa* ‘females’; and (**D**) brightly-lit *H. t. linaresi* and *H. heurippa* ‘females’ (means and CrIs in Table S5). Gardner-Altman plots are as in Figures 1B-D.

## DISCUSSION

By collecting ~1500h of mate choice data we have shown that the light environment can influence visual mating behaviours during the early stages of divergence in *Heliconius* butterflies. Although our data are characterised by considerable individual variation, we observed significant differences in behaviours of *H. heurippa* and *H. timareta linaresi* males, and this difference is stronger when the female patterns are more brightly lit. Experiments with live males and females revealed a degree of assortative mating, and the differences in visual attraction behaviours that we observe are sufficient to explain this. We have also shown that differences in visual attraction are associated with the presence of the red forewing band in interspecific hybrids under bright light conditions, perhaps suggesting physical linkage between an ecologically relevant colour pattern gene and those for the corresponding behaviour.

Studies of speciation often focus on already diverged groups, which are maintained by multiple reproductive barriers [53,54], making it difficult to draw conclusions about the role of the individual barriers and their ecological context. At the other extreme, barriers acting at the early stages of divergence may be of small effect and, especially in the case of behavioural phenotypes, require very large datasets to draw robust conclusions, which may not always be feasible. Behavioural researchers increasingly use computer vision [55], and software for automated tracking and individual identification is available (e.g. [56,57]). However, applying these techniques to footage with heterogenous backgrounds, variable light environments and arenas larger than the camera’s field of view remains a challenge. To overcome these limitations, we combined computer vision, allowing *post-hoc* motion detection, with subsequent human curation. This permitted a simple low-cost solution, and importantly allowed us to capture frames before and after motion is detected from many hundreds of hours of video footage.

This large volume of data revealed that increased illuminance at the red mounted *H. heurippa* female increased the frequency of interactions between *H. heurippa* males with *H. heurippa* females (relative to *H. t. linaresi* females). *H. heurippa* males were no more likely than *H. t. linaresi* males to interact with the *H. heurippa* female when she was poorly lit (lower 50% quantile of illuminance values; Fig. 2C), but were 1.3 times (i.e. 0.57/0.44, Table S5) more likely to interact with the *H. heurippa* female in brighter light (upper 50% quantile of illuminance values; Fig. 2D). Similar effects of the light environment have been observed in aquatic organisms. For example, the attractiveness of red colour during courtship is dependent on the spectrum of available light, influenced by water depth and turbidity, and recognition of con-specifics may fail entirely under certain light environments (e.g. [10–12,19]). However, far fewer studies have directly tested how changing light environments influence mating preference behaviours that contribute to population divergence in terrestrial organisms (but see [17,58,59]).

The mechanism underlying the differences in behaviour we observe under contrasting lighting conditions isn’t immediately clear. Insects, including butterflies [60], have frequently evolved colour constancy across light environments [61,62], and although this may only partially succeed [60,61], we generally expect individuals to be able to distinguish *H. heurippa* and *H. t. linaresi* patterns under different brightness conditions. Alternatively, male attraction to female colour patterns might to some degree be ‘wavelength-specific’, where triggering of behaviours depends greatly on an object’s emission of specific wavelengths and their intensity [62]. Under this scenario, a colour cue might only trigger a behaviour when presented at intense illuminance including wavelengths from the relevant part of the spectrum [62–64]. The differences in illuminance we measured may correlate with spectral differences; under shaded conditions, the available light may be reflected from the vegetation or from outside the path of direct sunlight, and consequently lack red wavelengths (but be rich in greenish or bluish light) [16,17]. Indeed, previous work modelling *Heliconius* vision suggests that red patterns are more conspicuous to *Heliconius* when presented in bright sunlight [41] (though whether this affects *Heliconius* behaviours has not previously been tested).

Regardless of the proximate mechanism, our data indicate that prevailing light conditions can influence visual mating behaviours in *Heliconius*. Unlike many other examples, often involving anthropogenic induced changes in aquatic environments [10–12], the effects we see here potentially act across very short time and spatial scales. This reflects the forest environment where the light environment can vary rapidly across both time and space. As such, male response may depend less on the broader environment (perhaps influenced by forest type), than on the movement of females across smaller spatial scales (for example between dimly and brightly lit patches). *Heliconius* butterflies are known to bask in the sun with their wings open, particularly during the morning hours [42]. This may primarily be for thermoregulation, though other species are known to display in environments where they are most conspicuous, and *Heliconius* might follow a similar strategy. *Anolis* lizards, for example, occupy micro-habitats in which their dewlap colour is most conspicuous [65]. Visual modelling indicates *Heliconius* red patterns are more conspicuous to avian predators when presented in bright sunlight, suggesting it may also enhance aposematism [41]. Whatever the ultimate reason for aggregating in sun-exposed patches, our data suggest that these behaviours could enhance the strength of divergent mating preferences.

Whether the differences in visual attraction behaviours that we observe translate to assortative mating, and therefore contribute to reproductive isolation, is an important question. Previous studies testing for assortative mating between *Heliconius* species in tetrad experiments similar to ours report low frequencies of interspecific mating [29,66,67]. This was not the case in our experiments. Nevertheless, the large number of trials in our study allowed us to detect positive assortative mating, albeit at low levels, though this was much stronger for the 60 trials in which *H. heurippa* males mated (Fig. 3). Unfortunately, we do not have illuminance data for these experiments, and this would be difficult to measure given the movement of females. Considering that the live females may have been basking in the sun, that overall activity of *Heliconius* is highest when it is sunny [68] and that mating in *Heliconius* seems to occur more frequently on sunny than on cloudy days (M. Linares, pers. obs.), it is likely that a large proportion of the trials with mating outcomes included sun-exposed females. When accounting for the more frequent involvement of *H. heurippa* males in mating, the mating rates between *H. heurippa* and *H. t. linaresi* in our tetrad experiments closely match estimates derived from our experiments with dead mounted females in bright light. Although we cannot rule out a role for cues acting across other sensory modalities, differences in visual attraction alone are sufficient to explain the levels of assortative mating we observe.

Considering our data for hybrids, it is overall difficult to ascertain patterns of dominance for visual attraction. This is perhaps not surprising given that the differences in behaviour between populations are subtle and are shaped by considerable variation. However, segregation of the *optix* alleles, which control red pattern elements in *Heliconius* [46], in the backcross to *H. t. linaresi* did allow us to test for linkage between the warning pattern cue and the corresponding behaviour. Once again, we observed illuminance-induced shifts in visual attraction, so that the two types of backcross to *H. t. linaresi* males (i.e. red/Bb vs. non-red/bb) differed in behaviour, but only under higher illuminance conditions. These results are consistent with physical linkage between behavioural loci and *optix*, as has been shown elsewhere – but with much greater effects – between the closely related species *H. cydno* and *H. melpomene* [27]. Physical linkage will help to maintain key genetic associations (i.e. LD) between loci underlying ecologically relevant traits and those for premating isolation, like mating preferences, facilitating speciation with gene flow in general [21,23], and hybrid trait speciation [69] in particular. Although the differences between backcross to *H. t. linaresi* genotypes account for ~50% of that observed in the two parental taxa, in absolute terms, the effects are unlikely to permit sufficient power for a formal QTL study and caution should be exercised when interpreting our results. Our results are consistent with a simple genetic mechanism by which behavioural alleles were acquired alongside red colour pattern alleles through introgression from *H. melpomene* into the ‘*heurippa/timareta/cydno*’ lineage [69,70]. However, it seems unlikely that the behavioural differences we observe here would rapidly lead to strong reproductive isolation, as predicted under a model of hybrid trait speciation [66,69].

In conclusion, the behavioural differences we observe for *H. heurippa* and *H. t. linaresi* are similar in strength to those reported elsewhere for other *Heliconius* taxa at the early stages of divergence (e.g. [30,53]). Alone, these may represent only weak barriers to gene flow; however, by augmenting divergent ecological selection acting on the warning pattern cue, they may facilitate the accumulation of additional barriers as speciation proceeds. In addition, our results suggest that the degree to which *Heliconius* warning pattern contribute to premating isolation may depend on local illuminance, which can change rapidly in both time and space. Without more stable differences in the sensory environment, perhaps facilitated by shifts in habitat use, these fluctuations may constrain speciation. However, *Heliconius* are known to aggregate and display their warning patterns in sun-exposed patches within the broader forest environment, and our results suggest that this context would enhance premating isolation. Traits predominantly shaped by ecology frequently act as mating cues, which by coupling divergent natural selection to premating isolation can promote speciation with gene flow [6]; our results highlight that this effect may depend on the sensory conditions during which these ecological mating cues are displayed to conspecifics.

## Supporting information

Supplemental R Markdown

## ACKNOWLEDGEMENTS

We are very grateful to the Abondano-Almeida family for helping AEH and MF settle in Colombia; Juan Sebastián Sánchez, Óscar Penagos, Isabel Leon and Lina Gabriela Melo for assistance in the insectaries; Annika Neuhaus, Morgan Oberweiser, Saoirse McMahon and Lucas Asis for help scoring videos; and Martin Küblbeck, Lucie Queste, Matteo Rossi, Daniel Shane Wright and Stephen Montgomery for comments on the manuscript. Field collections and insectary rearing were conducted under permit no. 530 issued by the Autoridad Nacional de Licencias Ambientales of Colombia (ANLA). CS and CP were funded by Universidad del Rosario (Grant Number: IV-FGD005). ML was partially funded by the Faculty of Natural Sciences at Universidad del Rosario. AEH, MF and RMM were supported by an Emmy Noether fellowship and research grant awarded to RMM by the Deutsche Forschungsgemeischaft (DFG) (Grant Number: GZ: ME 4845/1-1).

## Supplementary material

### 1) Supplementary Figures

**Figure S1:**
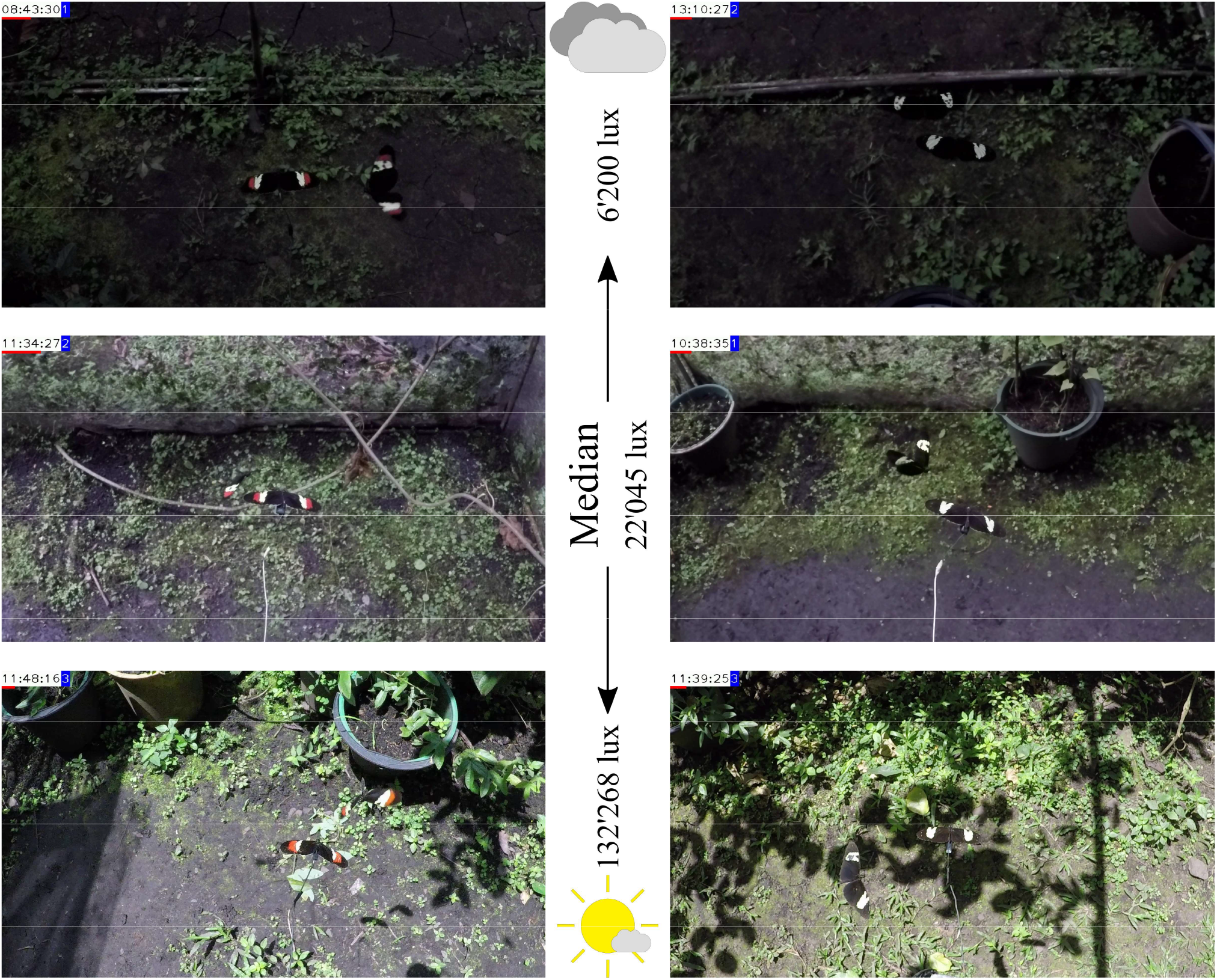
Examples from processed videos showing ‘poorly lit’, median and ‘brightly lit’ *H. heurippa* ‘female’ (left) and a *H. t. linaresi* ‘female’ as a comparison at same illuminance (right). Median illuminance at the *H. heurippa* ‘female’ was calculated from illuminance data at the *H. heurippa* ‘female’ when ‘pure’ males showed a response to either model. This median value was used as threshold to build the categorical model for ‘pure’ males. The exact same procedure was used for BL hybrid males. The specific *lux* values for ‘poorly lit’ and ‘brightly lit’ conditions (6’200 and 132’268 lux) shown here were randomly chosen from the bottom and the top quartile of the distribution. The presented frames were picked from different recording days. Additionally to the mounted female, each frame shows a conspecific male approaching/courting the ‘female’ and dots used to identify individual males are visible on their wings. A change in hue of the red patch on *H. heurippa* seems to be visible under different illuminance conditions, but it should not be forgotten that 1) human vision is vastly different from butterfly vision and 2) that this is a representation produced by the GoPro camera (GoPro, San Matteo, CA), which is post-processing colour-composition of frames.

**Figure S2:**
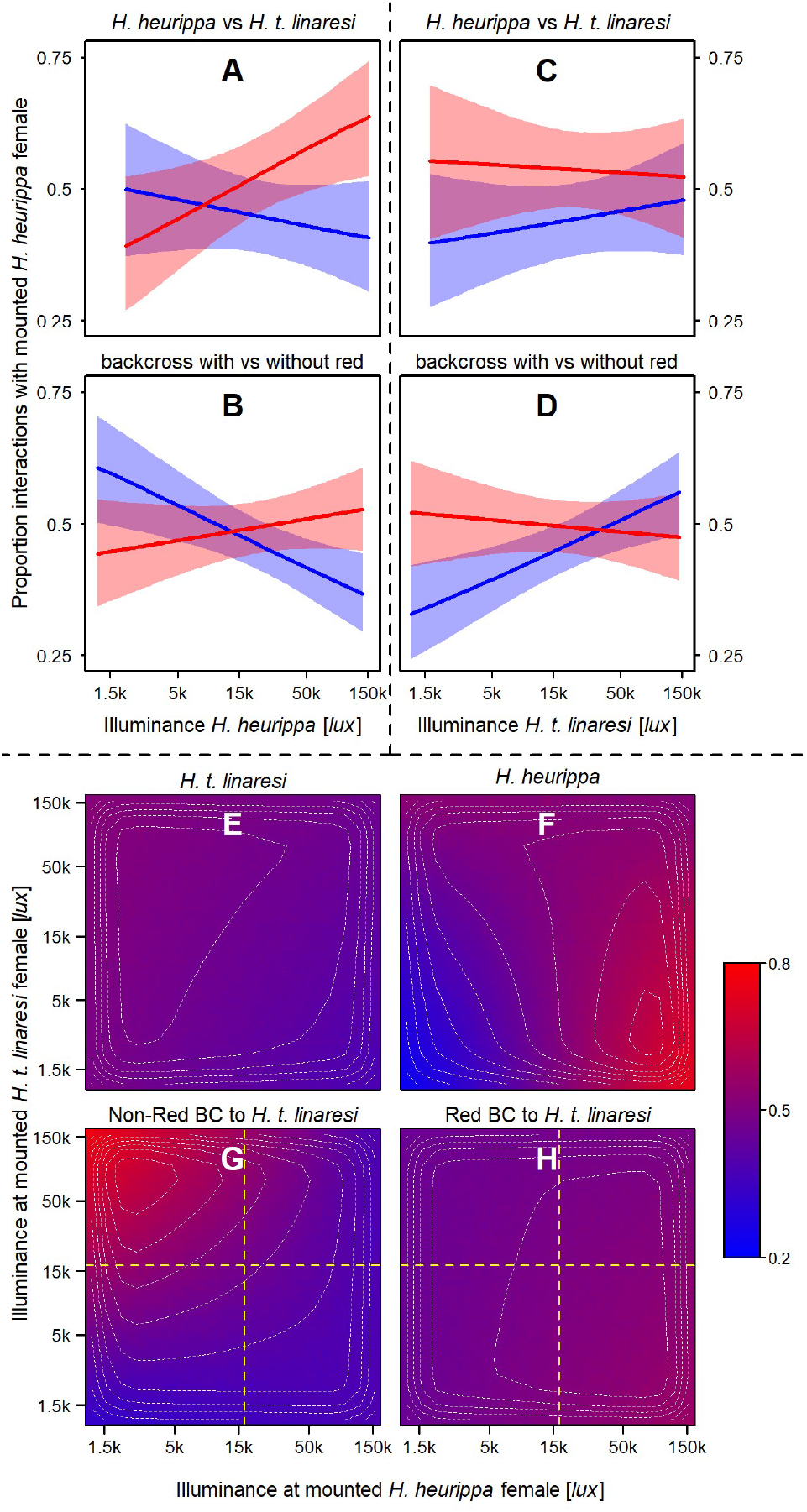
All interaction terms involving male type from fully saturated models (*i.e*. Table S2, model #3 for *H. heurippa* and *H. t. linaresi* males and Table S4, model #1 for backcross to *H. t. linaresi* males). **A+B**: Two-way interactions between illuminance at the *H. heurippa* mounted female and male type. Proportion of interactions with the *H. heurippa* female of *H. heurippa* (red) and *H. t. linaresi* (blue) males (A; resembles Fig. 2A, but comes from a different underlying model); and differently coloured backcross to *H. t. linaresi* males (B, red males (red), males without red (blue)). **C+D**: The same for the two-way interactions between illuminance at the *H. t. linaresi* mounted female and male type **E-H**: Three-way interactions between male type and the two illuminance measures. Male type is indicated on top of each graph. Colouration of each region in the graph depends on predicted preference for the *H. heurippa* mounted female at the given light conditions (scale depicted on right). White contour lines show Kernel densities weighted by preference score, which essentially gives the same information as the colour gradient (‘peaks’ are where the landscape is most red). All illuminance-axes are logarithmically scaled. Dashed lines in G and H show median illuminance measures at the respective axis, used to categorize conditions into ‘poorly’ and ‘brightly’ lit for the categorical models.

**Figure S3:**
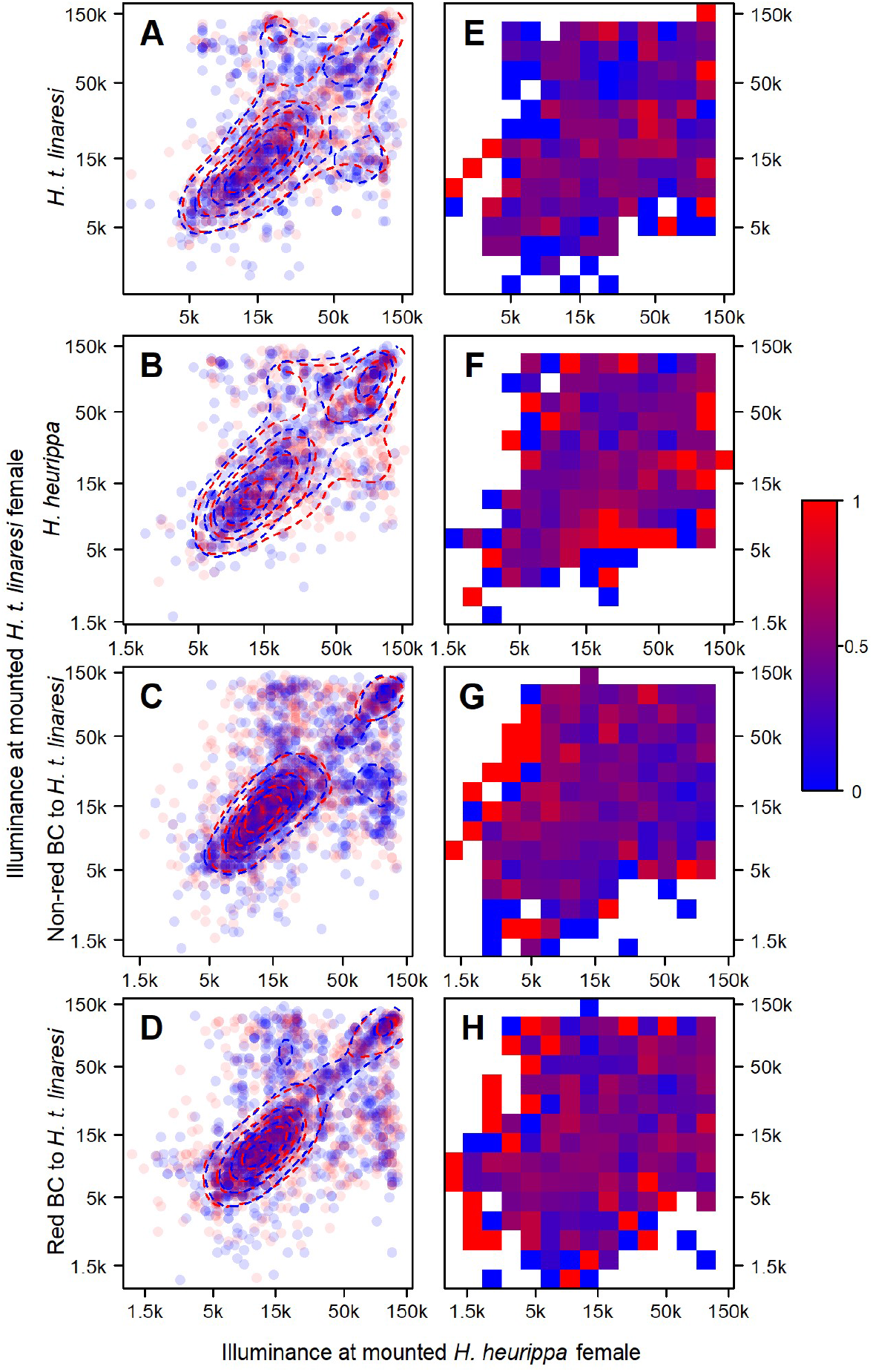
**A-D**: Raw data points of *H. t. linaresi, H. heurippa*, non-red and red backcross to *H. t. linaresi* males (from top to bottom) in a two-dimensional illuminance space. Horizontal dimension shows illuminance at the mounted *H. heurippa* female, vertical dimension shows illuminance at the mounted *H. t. linaresi* female. Blue dots indicate responses of males to the mounted *H. t. linaresi* female and red dots indicate responses to the mounted *H. heurippa* female. Kernel densities are shown for each set of data (blue lines for responses to *H. t. linaresi*, red lines for responses to *H. heurippa*). **E-H**: the two-dimensional illuminance space was then divided into 15 squares for each male type and local preferences within each square were calculated. Graphs directly relate to graphs A-D. The higher the preference for the mounted *H. heurippa* female in a square, the more red the square; the higher the preference for the *H. t. linaresi* female, the more blue (see scale on the right). Squares without response are white. These local preferences have to be interpreted in combination with the local sample sizes (as shown in left column), since they often rely on very few observations. All axes are logarithmically scaled.

**Figure S4:**
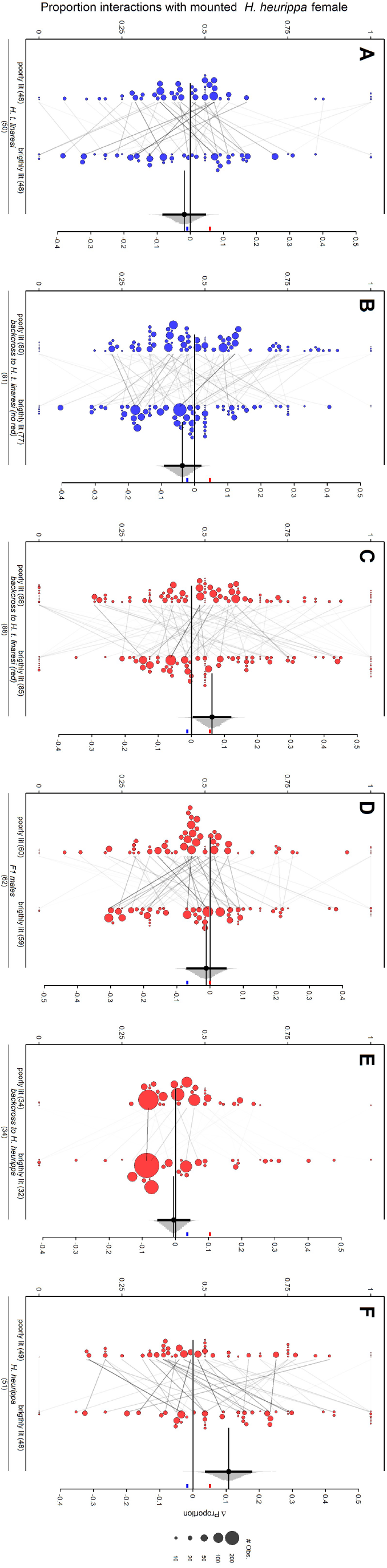
Individual trajectories from categorical models. Visual preferences of pure males and hybrid males are displayed by type. Within each type, data is split depending on illuminance around the *H. heurippa* mounted female, into poorly and brightly lit. Whenever an individual appeared under both light conditions, it is connected by a reaction norm line in this plot. Transparency of reaction norm lines within each panel is scaled by the maximum number of observations available for one individual within this panel. For *H. t. linaresi, H. heurippa* and BL males, posterior means for each EMM are calculated from the same underlying categorical models as used for Fig. 2 and Fig. 4, respectively. For F_1_ and BH males, a categorical model was fit including male type, the illuminance category around the *H. heurippa* ‘female’ (determined again by the median measurement) and their interaction. EMMs were then extracted using the same logic as for the other types. The difference between two groups (in this case between poorly and brightly lit conditions within each type) is shown by the same Gardner-Altman plot type as used for Fig. 2 and Fig. 4. Small red and blue line located at the right side of each panel show estimators for *H. t. linaresi* and *H. heurippa* males from Fig. 2B as a reference. [*Figure follows on next page*]

**Figure S5:**
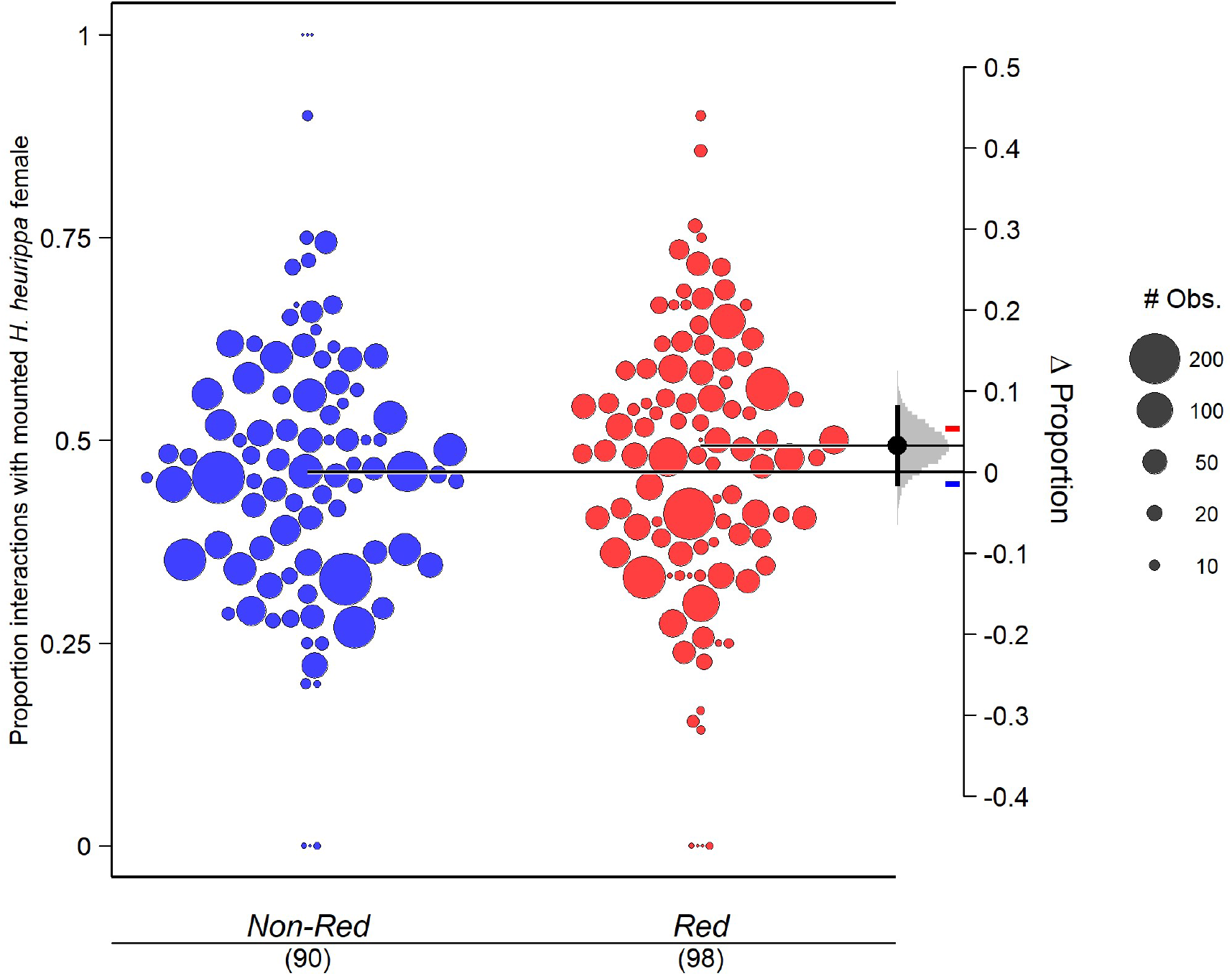
Relative attraction towards *H. heurippa* ‘females’ (i.e. the proportion of interactions with the mounted *H. heurippa* ‘female’ as opposed to the *H. t. linaresi* ‘female’) for non-red (*bb*) BL males (blue dots) and red (*Bb*) BL males (red dots). Data and estimators are across all light environments. Note that only preference data with associated illuminance data were included in the underlying statistical models (a few of the raw data points displayed here therefore were not considered by the underlying model). The difference between the two groups is shown with the same logic as used for Fig. 2 and Fig. 4. Small red and blue line located at the right side of each panel show estimators for *H. t. linaresi* and *H. heurippa* males from Fig. 2B as a reference.

### 2) Supplementary Tables

**Table S1:** Collection locations of *H. t. linaresi* and *H. heurippa*.

| Taxon | Location |
| --- | --- |
| *Heliconius timareta linaresei* | Guayabal (2°41'04"N, 74°53'17"W) |
| *Heliconius heurippa* | Lejanías (03°34'0"N, 74°04'20.4"W)<br>Buenavista (4°10'30"N, 73°40'41"W)<br>Santa María (04°53'28.2"N, 73°15'11.4"W) |

**Table S2:** Information on different experimental hybrid lines. Left column shows F_1_ broods from which lines originated. For the third line, we summarized two F_1_ crosses and their subsequent crosses into one hybrid line. These two F_1_ crosses had the same *H. heurippa* father, but a different *H. t. linaresi* mother. Individual counts in brackets show how many males of each brood were tested in the experiment. 32 other F_1_ males from 7 broods were tested in the experiment and are not mentioned here, as their brood was not involved in generating backcrosses (13 individuals/5 broods of those being from a cross between *H. t. linaresi* mother and *H. heurippa* father and 19 individuals/2 broods being from the reciprocal F_1_ cross with *H. heurippa* mother and *H. t. linaresi* father). *H. heurippa* is abbreviated as *heu* and *H. t. linaresi* as *lin* in the table.

| F<sub>1</sub> brood | Backcross to *lin* with male F<sub>1</sub> | Backcross to *lin* with female F<sub>1</sub> | Backcross to *heu* with male F<sub>1</sub> |
| --- | --- | --- | --- |
| mother: *lin*,<br>father: *heu* | mother: *lin*, father: F<sub>1</sub> | mother: F<sub>1</sub>, father: *lin* | mother: *heu*, father: F<sub>1</sub> |
| **C18_002 (12 indiv.)** | 7 broods (45 indiv.) | 2 broods (9 indiv.) | 2 broods (20 indiv.) |
| **C18_020 (6 indiv.)** | 3 broods (83 indiv.) | 1 brood (15 indiv.) | 1 brood (9 indiv.) |
| **C18_034 & C18_036 (half-sib.) (13 indiv.)** | 4 broods (18 indiv.) | 2 broods (10 indiv.) | 3 broods (6 indiv.) |
| **C19_014 (-)** | 1 brood (8 indiv.) | - | - |

**Table S3:** *GoPro Hero 5 Black™* (GoPro, San Matteo, CA) camera settings and equipment. All auto-settings (except were indicated) were disabled.

| Settings |  |
| --- | --- |
| Resolution | 1920 x 1080 pixels |
| Frame rate | 60 fps |
| Shutter speed | 1/480 |
| Field of view | *Narrow* |
| Colour | *GoPro* |
| Sharpness | *High* |
| ISO | 400 |
| White balance | *Auto* |
| Equipment | • 128 GB *SanDisk™ Extreme* SD card<br>• *Neewer™* UV filter (reduced overexposure of yellow bands)<br>• *Xlayer™* 210033 powerbank (charging while recording) |

**Table S4:** 19 possible fixed-effect structures for the pure males of *H*. heurippa and *H. t*. linaresi (fixed effect ‘type’). ‘I_H’ and ‘I_L’ stand for log-illuminance measures at the *H*. heurippa and the *H. t*. linaresi mounted females, respectively. A ‘*’ sign in a model formula indicates that all involved variables as well as all possible interactions between them are included. All models include trial and male ID as random effects. Including trial (and in many cases also individual ID) as a random effect has often been disregarded in previous analyses. However, we found strong correlation among individuals’ behaviour during trials (see supplementary R Markdown), likely due to the unique combination of males, mounted females and their position in the experimental cage during a given trial. The latter may have also affected sensitivity of our motion-detection software. We therefore considered it inevitable to correct for this. For each model we calculated a LOOIC and WAIC value, which mostly agree. Based on the WAIC value, differences between best model and other models are calculated (ΔWAIC) as well as a model-weight score.

| # | Fixed Effects Term | LOOIC | WAIC | $\Delta\text{WAIC}$ | Weight<sub>WAIC</sub> |
| --- | --- | --- | --- | --- | --- |
| 1 | ~ type * I_H | 3773.40 | 3768.52 | 0.00 | 0.29 |
| 2 | ~ type * I_H + I_L | 3774.63 | 3769.68 | 1.16 | 0.16 |
| 3 | ~ type * I_H * I_L | 3775.20 | 3770.08 | 1.56 | 0.13 |
| 4 | ~ type * I_H + I_H * I_L | 3775.99 | 3771.04 | 2.52 | 0.08 |
| 5 | ~ type * I_H + type * I_L | 3776.53 | 3771.52 | 3.01 | 0.06 |
| 6 | ~ type * I_H + type * I_L + I_H * I_L | 3776.62 | 3771.56 | 3.04 | 0.06 |
| 7 | ~ type | 3777.82 | 3773.07 | 4.55 | 0.03 |
| 8 | ~ type + I_L | 3778.10 | 3773.20 | 4.68 | 0.03 |
| 9 | ~ type + I_H | 3778.11 | 3773.33 | 4.81 | 0.03 |
| 10 | ~ I_L | 3778.50 | 3773.62 | 5.10 | 0.02 |
| 11 | ~ type * I_L | 3778.83 | 3773.95 | 5.43 | 0.02 |
| 12 | ~ 1 | 3779.24 | 3774.44 | 5.92 | 0.02 |
| 13 | ~ I_H | 3779.45 | 3774.58 | 6.06 | 0.01 |
| 14 | ~ type + I_H + I_L | 3779.75 | 3774.87 | 6.35 | 0.01 |
| 15 | ~ I_H + I_L | 3780.58 | 3775.58 | 7.06 | 0.01 |
| 16 | ~ type + I_H * I_L | 3780.77 | 3775.80 | 7.28 | 0.01 |
| 17 | ~ type * I_L + I_H | 3780.82 | 3775.99 | 7.47 | 0.01 |
| 18 | ~ type * I_L + I_H * I_L | 3781.61 | 3776.60 | 8.09 | 0.01 |
| 19 | ~ I_H * I_L | 3781.85 | 3776.78 | 8.26 | 0.00 |

**Table S5:** Posterior means and 95% equal-tailed credible intervals for male types from Fig. 2, Fig. S5 and Fig. 4. Minus (“-“) in the second column stands for a poorly lit mounted female, a plus (“+“) for a brightly lit one.

| Male type | Light environment | Mean | Lower | Upper |
| --- | --- | --- | --- | --- |
| *H. t. linaresi* (left in Fig. 2B) | Across all | 0.447 | 0.383 | 0.513 |
| *H. heurippa* (right in Fig. 2B) | Across all | 0.515 | 0.449 | 0.582 |
| *H. t. linaresi* (left in Fig. 2C) | *H. heurippa* - | 0.464 | 0.391 | 0.540 |
| *H. heurippa* (right in Fig. 2C) | *H. heurippa* - | 0.456 | 0.385 | 0.527 |
| *H. t. linaresi* (left in Fig. 2D) | *H. heurippa* + | 0.437 | 0.362 | 0.514 |
| *H. heurippa* (right in Fig. 2D) | *H. heurippa* + | 0.572 | 0.493 | 0.650 |
| BL without red (left in Fig. S5) | Across all | 0.461 | 0.414 | 0.508 |
| BL with red (right in Fig. S5) | Across all | 0.493 | 0.444 | 0.543 |
| BL without red (left in Fig. 4A) | *H. heurippa* - / *H. t. linaresi* - | 0.460 | 0.406 | 0.515 |
| BL with red (right in Fig. 4A) | *H. heurippa* - / *H. t. linaresi* - | 0.475 | 0.419 | 0.531 |
| BL without red (left in Fig. 4B) | *H. heurippa* - / *H. t. linaresi* + | 0.477 | 0.401 | 0.554 |
| BL with red (right in Fig. 4B) | *H. heurippa* - / *H. t. linaresi* + | 0.443 | 0.359 | 0.528 |
| BL without red (left in Fig. 4C) | *H. heurippa* + / *H. t. linaresi* - | 0.426 | 0.350 | 0.505 |
| BL with red (right in Fig. 4C) | *H. heurippa* + / *H. t. linaresi* - | 0.538 | 0.461 | 0.616 |
| BL without red (left in Fig. 4D) | *H. heurippa* + / *H. t. linaresi* + | 0.438 | 0.384 | 0.491 |
| BL with red (right in Fig. 4D) | *H. heurippa* + / *H. t. linaresi* + | 0.504 | 0.446 | 0.563 |

**Table S6:** Posterior means and 95% equal-tailed credible intervals for estimates of proportions of each mating outcome from tetrad experiments. First row is based on empirically measured data, the other rows are stochastic predictions based on mounted female data from all light environments, from a poorly lit *H. heurippa* mounted female and a brightly lit one, respectively.

| Estimates based on | *H. heurippa* ♀<br>*H. t. linaresi* ♂ | *H. t. linaresi* ♀<br>*H. t. linaresi* ♂ | *H. heurippa* ♀<br>*H. heurippa* ♂ | *H. t. linaresi* ♀<br>*H. heurippa* ♂ |
| --- | --- | --- | --- | --- |
| tetrad experiment | 0.168<br>[0.099; 0.253] | 0.157<br>[0.09; 0.239] | 0.405<br>[0.305; 0.508] | 0.27<br>[0.183; 0.366] |
| mounted females:<br>across all | 0.149<br>[0.084; 0.23] | 0.184<br>[0.111; 0.27] | 0.349<br>[0.255; 0.45] | 0.329<br>[0.236; 0.429] |
| mounted females: poorly<br>lit *H. heurippa* | 0.152<br>[0.086; 0.233] | 0.181<br>[0.109; 0.266] | 0.315<br>[0.224; 0.414] | 0.363<br>[0.267; 0.465] |
| mounted females:<br>brightly lit *H. heurippa* | 0.146<br>[0.082; 0.226] | 0.187<br>[0.114; 0.273] | 0.387<br>[0.289; 0.489] | 0.291<br>[0.203; 0.389] |

**Table S7:** 19 possible fixed-effect structures for the BL males (fixed effect ‘type’ refers here to wing colour phenotype of hybrids). ‘I_H’ and ‘I_L’ stand for log-illuminance measures at the *H. heurippa* and the *H. t. linaresi* mounted females, respectively. A ‘*’ sign in a model formula indicates that all involved variables as well as all possible interactions between them are included. All models include trial, male ID and brood as random effects. Each model structure has a LOOIC and WAIC value, which mostly agree. Based on the WAIC value, differences between best model and other models are calculated (ΔWAIC) as well as a model-weight score.

| # | Fixed Effects Term | LOOIC | WAIC | ΔWAIC | Weight<sub>WAIC</sub> |
| --- | --- | --- | --- | --- | --- |
| 1 | ~ type * I_H * I_L | 6815.06 | 6811.69 | 0.00 | 0.38 |
| 2 | ~ type * I_H + type * I_L + I_H * I_L | 6815.98 | 6812.60 | 0.91 | 0.24 |
| 3 | ~ type * I_H + type * I_L | 6817.15 | 6813.84 | 2.15 | 0.13 |
| 4 | ~ type * I_H + I_H * I_L | 6817.82 | 6814.43 | 2.74 | 0.10 |
| 5 | ~ type * I_H | 6820.15 | 6816.86 | 5.17 | 0.03 |
| 6 | ~ type * I_H + I_L | 6820.78 | 6817.48 | 5.79 | 0.02 |
| 7 | ~ type + I_H * I_L | 6821.33 | 6817.96 | 6.27 | 0.02 |
| 8 | ~ I_H * I_L | 6822.03 | 6818.67 | 6.98 | 0.01 |
| 9 | ~ type | 6821.94 | 6818.68 | 6.99 | 0.01 |
| 10 | ~ type * I_L + I_H * I_L | 6822.28 | 6818.95 | 7.27 | 0.01 |
| 11 | ~ 1 | 6822.46 | 6819.18 | 7.50 | 0.01 |
| 12 | ~ type + I_H | 6822.96 | 6819.70 | 8.01 | 0.01 |
| 13 | ~ type + I_L | 6823.18 | 6819.94 | 8.26 | 0.01 |
| 14 | ~ I_H | 6823.50 | 6820.19 | 8.50 | 0.01 |
| 15 | ~ I_L | 6823.80 | 6820.52 | 8.84 | 0.00 |
| 16 | ~ type + I_H + I_L | 6824.36 | 6821.03 | 9.35 | 0.00 |
| 17 | ~ type * I_L | 6825.21 | 6821.92 | 10.23 | 0.00 |
| 18 | ~ I_H + I_L | 6825.20 | 6821.93 | 10.24 | 0.00 |
| 19 | ~ type * I_L + I_H | 6825.55 | 6822.23 | 10.54 | 0.00 |

