## Supplemental R Markdown for "Light environment influences mating behaviours during the early stages of divergence in tropical butterflies": Hausmann_et_al_2020_Markdown.html

- 1 Initial explantions and setup
- 2 Choice experiment with *heu* and *lin* mounted females
  - 2.1 Data Read-in & Exploration
    - 2.1.1 Read in data
    - 2.1.2 Correlation of individual preferences during a trial with preference of other males
    - 2.1.3 Potential for collinearity between the illuminance measures
    - 2.1.4 Adding new columns
    - 2.1.5 Subsetting and Scaling
  - 2.2 Preference of *H. t. linaresi*, *H. heurippa* and backcross-to-*H. t. linaresi* hybrids and the effect of illuminance at the mounted *H. heurippa* female
    - 2.2.1 Modelling, Posteriors and CrIs
      - 2.2.1.1 Effect of type and its interaction with illuminance measures on visual mate preference in *heu* and *lin* males
      - 2.2.1.2 Effect of presence/absence of red colour in backcross hybrids and its interaction with illuminance measures on visual mate preference
    - 2.2.2 Figure 2
    - 2.2.3 Figure 4
    - 2.2.4 Supplementary figure 5
    - 2.2.5 Supplementary figure 4
    - 2.2.6 Supplementary figure 2
    - 2.2.7 Supplementary figure 3
- 3 Tetrad experiments with alive females
  - 3.1 Modelling
  - 3.2 Posteriors
  - 3.3 Plot, CrIs hypothesis tests
    - 3.3.0.1 CrI
    - 3.3.0.2 Numeric predictions given mounted female data
    - 3.3.0.3 Figure 3
    - 3.3.0.4 Hypothesis testing
- 4 Session Info

Alexander E. Hausmann

last edited: 22 July 2020

### 1 Initial explantions and setup

**Note**: the order of the analyses in this Markdown does not always follow the order of the Results section from the manuscript.

**Note**: *Heliconius heurippa* will be abreviated with ‘*heu*’, *Heliconius timareta linaresi* with ‘*lin*’

**Note**: even though we differentiated between ‘approaches’, ‘courtship initiation’ and ‘sitting on mounted female’ during scoring of the video material, we unite these three behaviours here as ‘interactions’ and analyse them as one variable. All three types of behaviours always include an approach of the male, which means that we are basically only looking at approaches in this analysis. All three behaviours were correlated within types and uniting them increases our power to detect differences.

**Note**: if we speak of preference values in the following, these refer to the proportion of interactions males perform towards the *heu* ‘female’ (a preference score of 0 indicates that all interactions were performed towards the *lin* ‘female’, a value of 1 indicates that all interactions were performed towards the *heu* ‘female’, a value of 0.5 indicates that there was equally many interactions with each type of ‘female’).

**Note**: to a point estimate for posterior distributions, we will use (unconventionally) the **mean** here (not the median, as most people do). The reason for that is that we will perform a lot of “posterior calculus” below. For example, for the Gardner-Altman plots below, we will plot the estimators of two groups, say A and B, as well as the estimator for the difference between these two groups. Now, median(B)-median(A) does most likely not equal median(B-A). But mean(B)-mean(A) equals mean(B-A). Therefore, for all point estimators to add up to values that make sense as a whole, we have to use the mean. To describe the 95% credible interval (CrI), we will use (as quite common), the 95% equal-tailed interval. Alternatively, one could use the high density interval.

Call `plot_proportion_stats.R`, which contains a function that will later be used for producing the stripchart graphs. We outsourced this function to GitHub to make this Markdown more concise.

```
source("https://raw.githubusercontent.com/SpeciationBehaviour/visual_preference_heurippa_linaresi/master/Analyses_and_Data/Skripts_called_from_Markdown/Plot_Proportion_Stats.R")
```

Call `other_plot_functions.R` which contains functions that will be used below for plotting and which are saved in a separate script, again, to make this Mardown more concise.

```
source("https://raw.githubusercontent.com/SpeciationBehaviour/visual_preference_heurippa_linaresi/master/Analyses_and_Data/Skripts_called_from_Markdown/other_plot_functions.R")
```

To be safe, we set a seed here (but all functions requiring a random number generator should have a seed set within the function).

Note that if you use an R version < 3.6 you may not be able to reproduce our exact results on your computer since R implemented a new mechanism for random number generation and set it as the new default. But this new option is not available in older versions.

```
set.seed(42)
```

Load/install libraries

For Bayesian regression models (using ‘Stan’)

```
if (!"brms" %in% rownames(installed.packages())) {
    install.packages("brms")
}
suppressMessages(suppressWarnings(library(brms)))
```

Note that for `brms` to work, you need to have a C++ compiler (which can be the R internal `Rtools`) as well as the R library `rstan` installed. Both may be installed automatically if you install `brms`. We ran analyses on a Windows machine and had to re-install `rstan`, since otherwise some models crashed. Use the following code if you’re on Windows:

`install.packages("rstan", type = "win.binary", dependencies = TRUE)`

For glmer

```
if (!"lme4" %in% rownames(installed.packages())) {
    install.packages("lme4")
}
suppressMessages(suppressWarnings(library(lme4)))
```

For estimated marginal means and contrasts between different factors.

```
if (!"emmeans" %in% rownames(installed.packages())) {
    install.packages("emmeans")
}
suppressMessages(suppressWarnings(library(emmeans)))
```

For weighted histograms

```
if (!"weights" %in% rownames(installed.packages())) {
    install.packages("weights")
}
suppressMessages(suppressWarnings(library(weights)))
```

For converting PDF to PNG

```
if (!"pdftools" %in% rownames(installed.packages())) {
    install.packages("pdftools")
}
suppressMessages(suppressWarnings(library(pdftools)))
```

For reading in PNGs

```
if (!"png" %in% rownames(installed.packages())) {
    install.packages("png")
}
suppressMessages(suppressWarnings(library(png)))
```

For displaying PNGs

```
if (!"grid" %in% rownames(installed.packages())) {
    install.packages("grid")
}
suppressMessages(suppressWarnings(library(grid)))
```

For weighted 2-D Kernel densities we need `ggtern`.

```
if (!"ggtern" %in% rownames(installed.packages())) {
    install.packages("ggtern")
}
suppressMessages(suppressWarnings(library(ggtern)))
```

`MASS` is used for (unweighted) Kernel densitites.

```
if (!"MASS" %in% rownames(installed.packages())) {
    install.packages("MASS")
}
suppressMessages(suppressWarnings(library(MASS)))
```

### 2 Choice experiment with *heu* and *lin* mounted females

#### 2.1 Data Read-in & Exploration

##### 2.1.1 Read in data

Keep all strings as characters and change the relevant columns to factors later.

```
pref_stat <- read.csv("Data/HC_TLC_choice_all_static.csv", header = T, 
    stringsAsFactors = F)
```

One row of the table represents one single interaction between a male and a mounted female.

The columns indicate:

- **court\_lin\_heu\_01** = *0*: Interaction of male with *lin* ‘female’, *1*: Interaction with *heu* ‘female’
- **behaviour** = Type of interaction with ‘female’; *approach*: approach, *court\_init*: approach + courtship initiation, *sitting*: approach + courtship initiation + sitting on ‘female’
- **ID** = Individual identifier
- **type** = *HC*: *heu*, *TLC*: *lin*, *TLCxHC*: F1 with *lin* mother and *heu* father (this type was used to produce backcrosses), *HCxTLC*: F1 with *heu* mother and *lin* father, *(TLCxHC)xTLC*: (female-informative) backcross to *lin* with F1 mother, *TLCx(TLCxHC)*: (male-informative) backcross to *lin* with F1 father, *HCx(TLCxHC)*: (male-informative) backcross to *heu* with F1 father
- **redYN** = Whether male has red on wings (*Y*) or not (*N*)
- **date** = Date in yyyy\_mm\_dd
- **trial** = Trial of the day
- **time\_of\_day** = Time of day in seconds
- **age\_male** = Age of male
- **HC\_model\_ID** = ID of *heu* ‘female’
- **TLC\_model\_ID** = ID of *lin* ‘female’
- **position\_HC** = Recording position of *heu* ‘female’ in cage
- **position\_TLC** = Recording position of *lin* ‘female’ in cage
- **light\_HC** = lux intensity at *heu* ‘female’
- **light\_TLC** = lux intensity at *lin* ‘female’
- **temp** = Temperature
- **hum** = Humidity
- **observer** = Observer ID
- **broods** = Stock individuals (pure species) indicated with *Stock* in beginning. Equal identifiers may indicate that males are brothers (if born around the same date), but may also not. Hybrid cross IDs always start with *C1…*; males with same cross ID are always brothers.
- **HC\_type** = Only of relevance for *heu* males. *Old\_Stock* are males from the first *heu* stock we used, which was created from populations from Lejanias, Buenavista and Santa Maria. *SM-Stock* males come from a stock which exclusively traces back to the population from Santa Maria.
- **exp\_line** = Only of relevance for *lin*-*heu* hybrids. Gives their affiliation with a certain F1 cross which were each used to generate multiple backcrosses. E.g. TLCx(TLCxHC) males may be of different broods, but from the same experimental line. This means they trace back to the same F1 cross and had therefore the same *heu* grandfather. Note that line\_3 males do not result from the same F1 cross, but have the same *heu* grandfather (who mated two *lin* females).
- **males\_inside** = Number of males in cage during day of experiment.
- **resp\_other\_males** = Number of interactions that other males showed with either ‘female’ in the trial during which data point was generated
- **pref\_other\_males** = Preference of other males in the trial during which data point was generated.

Discard immature male data (we assume only males of an age of 5+ days will be sexually mature (and therefore show preferences)).

```
pref_stat <- pref_stat[pref_stat$age_male > 4, ]
```

##### 2.1.2 Correlation of individual preferences during a trial with preference of other males

We first want to get a brief idea of how preference of individual males during a trial correlates with preference of all other males during that trial (mind though correlation vs. causation!).

**Note**: our data sheet does not contain all data from all males that were flying through our cage during experiments (we excluded data from a few males from types that play no role for this publication to keep the data sheet more understandable). The column `pref_other_males` was though calculated including those males into the calculations (and therefore shows the “correct” preference of all other males present).

We fit some easy generalized linear mixed-effects models with `glmer`. To just get a quick idea of the correlations, we predict the probabilities of a male to respond to the *heu* ‘female’ by setting preference of other males either to 0, 0.5 or 1. We first do this for all males appearing in the experiment and then look at the different types separately. Individual ID is set as random effect.

```
# Run model for all males
all_corr <- glmer(court_lin_heu_01 ~ pref_other_males + (1 | 
    ID), family = binomial, data = pref_stat)
# Predict preference of individual while preference of other
# males is at 0, 0.5 or 1.
pred_all_corr <- as.numeric(predict(all_corr, data.frame(pref_other_males = c(0, 
    0.5, 1)), type = "response", re.form = NA))
# Create data frame which will be filled with additional
# information on all the types later
corr_data <- data.frame(type = "ALL", predict_0 = round(pred_all_corr[1], 
    2), predict_0.5 = round(pred_all_corr[2], 2), predict_1 = round(pred_all_corr[3], 
    2), stringsAsFactors = F, row.names = c())
# Get all unique types
unique_types <- sort(unique(pref_stat$type))
# Run through all types
for (typo in 1:length(unique_types)) {
    # Run the model for the current type
    sub_corr <- glmer(court_lin_heu_01 ~ pref_other_males + (1 | 
        ID), family = binomial, data = pref_stat[pref_stat$type == 
        unique_types[typo], ])
    pred_sub_corr <- as.numeric(predict(sub_corr, data.frame(pref_other_males = c(0, 
        0.5, 1)), type = "response", re.form = NA))
    # Append to data frame
    corr_data <- rbind(corr_data, c(unique_types[typo], round(pred_sub_corr, 
        2)))
}
# Show results
corr_data
```

We can see that the higher the preference for the *heu* ‘female’ of other males during a trial, the more likely it is that an individual interacts with the *heu* ‘female’, throughout all types of males. This may be due to:

- Light conditions differing at the two recording stations (which may not only affect male choice, but also whether our program detected movement or not)
- Certain recording stations being more attractive because of other reasons (e.g. because of closer proximity to food)
- One of the ‘females’ being specifically attractive/unattractive during a recording day
- Learning of individuals from other individuals

While the first three would speak for mere correlation between behaviour of the butterflies in our cage, the latter would speak for causation in a sense that individuals learn from each other. We decided not to include preference of other males during a trial in our models later on, since we cannot really correct for number of responses of other males (e.g. a preference score for other males coming from one observation during a trial results in a skewed preference of 0 or 1 and this will get the same weight as a score coming from many observations); this is also why the presented table above has to be interpreted with CAUTION.

We also avoided including recording position in the cage of the two mounted females during a trial as random effect in our models (e.g. `(1|position_HC) + (1|position_TLC) + (1|position_HC:position_TLC)`). Variance explained by cage positions is tighly linked with illuminance differences since certain positions always were more prone to exposure to sun illuminance than others. Since conditions around the recording positions also changed immensly over the course of the experiment, including it as a “static” factor seemed to be questionable.

To avoid overcomplication of the model, we also did not include ‘mounted female model’-IDs into our statistical model (with e.g. `(1|HC_model_ID) + (1|HC_model_ID) + (1|HC_model_ID:TLC_model_ID)`). Both variables contained a large amount of factors and added little improvement to model fit. Also, same as for the recording positions, attractiveness of ‘females’ may have changed over the course of their use, which is why including them as “static” factors also would bear problems.

We did attempt correcting for all the variance caused by all these different effects by including trial (and date) as random effects in the models (see below). This should cover the majority of differences in recording positions and ‘female’ attractiveness.

##### 2.1.3 Potential for collinearity between the illuminance measures

Since our mounted females were always set up in relatively close proximity to each other, we expect the two illuminance measurements to be correlated. Assuming logarithmic perception of illuminance in Heliconius (see more detailed explanation later), we plot the “log-illuminance” at the *heu* ‘female’ (`log10` of *heu* lux measurement) against the “log-illuminance” at the *lin* ‘female’ (`log10` of *lin* lux measurement).

```
par(mar = c(4, 4, 0.3, 0))
plot(log10(pref_stat$light_HC), log10(pref_stat$light_TLC), pch = 20, 
    col = adjustcolor("black", 0.03))
```

We see that those two are quite correlated. If we look at the correlation coefficient and the \({R^2}\) value, we can see that they do show some degree of correlation, which is though not enough to be worried about including both measurements into the same model.

```
cor(log10(pref_stat$light_HC), log10(pref_stat$light_TLC), use = "pairwise.complete.obs")
```

```
## [1] 0.5702961
```

```
summary(lm(log10(pref_stat$light_HC) ~ log10(pref_stat$light_TLC)))$r.squared
```

```
## [1] 0.3252376
```

##### 2.1.4 Adding new columns

In short, lux is a measurement of the amount of photons hitting a surface per time. We assume that perception of illuminance in *Heliconius*, as in humans, is logarithmically scaled, i.e. a change from 100 and 1’000 lux (illuminance) would be perceived as much of a change as going from 1’000 to 10’000 lux (such ‘logarithmic perception of illuminance’ has been shown in in the fly, see Laughlin, Simon B. “Matching coding, circuits, cells, and molecules to signals: general principles of retinal design in the fly’s eye.” Progress in retinal and eye research 13.1 (1994): 165-196.). The exact formula for defining ‘lighting steps’ (i.e. whether e.g. 1000 lux would be perceived twice or thrice as bright as 100 lux) is hard to predict for *Heliconius* butterflies. We later center and scale all log-illuminance variables to make the choice of the base of the logarithm irrelevant.

To calculate the log-illuminance score at each ‘female’ we take the logarithm to the base 10 of each illumance measurement.

```
pref_stat$log_light_HC <- log10(pref_stat$light_HC)
pref_stat$log_light_TLC <- log10(pref_stat$light_TLC)
```

We quickly compare our total recorded log-illuminance values to the log-illuminance values when the males actually responded. For this we will load two big txt files containing illuminance measures at the two ‘females’ at every second of the experiment. These data will only be used for this (not so relevant) following exploratory graph (and most likely we won’t provide these online, since file size is so big).

Read in illuminance data for all seconds of the experiment.

```
lux_heu <- read.table("Data/Light_heurippa.txt", header = T, 
    stringsAsFactors = F)
lux_lin <- read.table("Data/Light_linaresi.txt", header = T, 
    stringsAsFactors = F)
```

Calculate range of log-illuminance values.

```
range_all <- range(c(log10(lux_heu$lux), log10(lux_lin$lux), 
    pref_stat$log_light_HC, pref_stat$log_light_TLC), na.rm = T)
```

Generate histograms for the total recorded log-illuminance values at the *heu* ‘female’ as well as the log-illuminance values at the *heu* ‘female’ when responses occured at this ‘female’. The histograms for the the total recorded log-illuminance are weighted by number of individual males present in the cage at each respective day. Do the same for the *lin* ‘female’.

```
hist_heu_light <- wtd.hist(log10(lux_heu$lux), weight = lux_heu$males_5d, 
    plot = F, breaks = seq(range_all[1], range_all[2], length.out = 30))
hist_lin_light <- wtd.hist(log10(lux_lin$lux), weight = lux_lin$males_5d, 
    plot = F, breaks = seq(range_all[1], range_all[2], length.out = 30))

hist_heu_resp <- hist(log10(pref_stat$light_HC[pref_stat$court_lin_heu_01 == 
    1]), plot = F, breaks = seq(range_all[1], range_all[2], length.out = 30))
hist_lin_resp <- hist(log10(pref_stat$light_TLC[pref_stat$court_lin_heu_01 == 
    0]), plot = F, breaks = seq(range_all[1], range_all[2], length.out = 30))
```

Scale the histograms to have the same area.

```
hist_heu_resp$counts <- hist_heu_resp$counts * (sum(hist_heu_light$counts)/sum(hist_heu_resp$counts))
hist_lin_resp$counts <- hist_lin_resp$counts * (sum(hist_lin_light$counts)/sum(hist_lin_resp$counts))
```

Plot histograms! Black for total recorded, beige for log-illuminance when responses occurred.

```
# Layout setup
layout(matrix(1:2, nrow = 1))
par(oma = c(0, 0, 1, 0))
par(mar = c(4.2, 0.5, 1.5, 0.5))

# Add histograms
plot(hist_heu_light, col = adjustcolor("black", 0.5), ylab = "", 
    xlab = "log10 lux", yaxt = "n", main = substitute(paste("Mounted ", 
        italic("H. heurippa"), " female")))
plot(hist_heu_resp, col = adjustcolor("goldenrod", 0.5), add = T)

plot(hist_lin_light, col = adjustcolor("black", 0.5), ylab = "", 
    xlab = "log10 lux", yaxt = "n", main = substitute(paste("Mounted ", 
        italic("H. t. linaresi"), " female")))
plot(hist_lin_resp, col = adjustcolor("goldenrod", 0.5), add = T)

# Add title
mtext("Total Recorded vs. Responses", 3, outer = T, line = 0)
```

```
# Reset layout
layout(1)
```

We can see that the recording density is roughly the same for both ‘females’ and also the distribution of the responses is similar. At both mounted females, way more responses occurred at bright light conditions (more male activity or higher sensitivity of the motion-detection software).

Finally, we add a new column to the dataset where we concatenate date and trial (to get an ID for each unique trial)

```
pref_stat$date_trial <- paste0(pref_stat$date, "_", pref_stat$trial)
```

##### 2.1.5 Subsetting and Scaling

We create one new subset table for the *lin*/*heu* dataset and the backcross dataset.

```
pref_stat_TLC_HC <- pref_stat[pref_stat$type %in% c("TLC", "HC"), 
    ]
pref_stat_BC <- pref_stat[pref_stat$type %in% c("TLCx(TLCxHC)", 
    "(TLCxHC)xTLC"), ]
```

We then center and scale `log_light_HC` and `log_light_TLC` in the two datasets. The scaling ensures that the chosen base of logarithm will not play a role for model outcome and centering is relevant for avoiding unproportional leverage of some data points.

```
pref_stat_TLC_HC$log_light_HC.z <- scale(pref_stat_TLC_HC$log_light_HC, 
    center = T, scale = T)
pref_stat_TLC_HC$log_light_TLC.z <- scale(pref_stat_TLC_HC$log_light_TLC, 
    center = T, scale = T)
pref_stat_BC$log_light_HC.z <- scale(pref_stat_BC$log_light_HC, 
    center = T, scale = T)
pref_stat_BC$log_light_TLC.z <- scale(pref_stat_BC$log_light_TLC, 
    center = T, scale = T)
```

#### 2.2 Preference of *H. t. linaresi*, *H. heurippa* and backcross-to-*H. t. linaresi* hybrids and the effect of illuminance at the mounted *H. heurippa* female

##### 2.2.1 Modelling, Posteriors and CrIs

Define vector of column names involved in models that should become factors and make those columns factors in the three tables.

```
make_fact <- c("type", "redYN", "ID", "broods", "exp_line", "date", 
    "trial", "date_trial", "observer")
for (change_fact in 1:length(make_fact)) {
    pref_stat_TLC_HC[, make_fact[change_fact]] <- as.factor(pref_stat_TLC_HC[, 
        make_fact[change_fact]])
    pref_stat_BC[, make_fact[change_fact]] <- as.factor(pref_stat_BC[, 
        make_fact[change_fact]])
}
```

The full (initial) model encompasses the following fixed effects:

1. **type** (which (species) type a male belongs to) *OR* **redYN** (=genotype around *optix*, i.e. whether a backcross-to-*lin* hybrid has or has not red on its wings)
2. **log\_light\_HC** (the scaled and centered log-illuminance at the *heu* ‘female’)
3. **log\_light\_TLC** (the scaled and centered log-illuminance at the *lin* ‘female’)
4. The **interaction** between 1) and 2)
5. The **interaction** between 1) and 3)
6. The **interaction** between 2) and 3)
7. The **interaction** between 1), 2) and 3)

Of particular interest are those fixed effects which involve **type** or **redYN** (1, 4, 5, 7).

As random effects, we decide for the following possible options:

1. **ID**: necessary to prevent pseudo-replication/overdispersion.
2. **date** and/or (nested) **date\_trial**: this also allows to control for pseudo-replication during experimental units, within dates/trials (e.g. Chamberlain, Nicola L., et al. “Polymorphic butterfly reveals the missing link in ecological speciation.” Science 326.5954 (2009): 847-850. included trial as random effect). We use the column **date\_trial** as a unique trial identifier (since trial IDs repeat over days).
3. **observer**: control for the person that scored a data point from video material.
4. **exp\_line** (only for models involving hybrids): experimental line a hybrid belongs to, i.e. from which F1 cross it resulted.
5. **broods** (only for models involving hybrids): brood a hybrid belongs to, i.e. from which backcross it resulted.

While random effect 1) is absolutely necessary to include, we decided to check for all other random effects whether it makes sense to include them or not (see below).

We set some relatively weak priors for the `intercepts` and all additional `type`/`redYN`-coefficients in all models in this section. Note that these model coefficients represent the model estimators for different male types when all log-illuminance measurements are at their minimum. We use the normal distribution, with a mean of 0 and a standard deviation of 3. The used priors help us eliminate extreme preference values close to 0 and 1. Due to close relationship between the types, we don’t have any clear expectation for their preferences, except that those preferences will not be close to 0 or 1.

Note that this prior acts on the logit scale (all our models use a logit-link), so we have to inverse-logit transform our prior to make it understandable. We quickly visualize what this prior looks like when inverse-logit transformed:

```
plot(plogis(seq(-5, 5, 0.01)), dnorm(seq(-5, 5, 0.01), 0, 3), 
    type = "l", xlab = "Prior (Inverse Logit Scale)", ylab = "Probability")
```

Recode contrasts (recommended to retrieve good model estimates later from `brm` model). (Another option would be using `contr.bayes`, as suggested by some people.) First, save old contrasts.

```
old_contr <- options("contrasts")
options(contrasts = c("contr.sum", "contr.poly"))
```

###### 2.2.1.1 Effect of type and its interaction with illuminance measures on visual mate preference in *heu* and *lin* males

###### 2.2.1.1.1 Random effects

We first look at *heu* and *lin* males. We start off checking how stable our models will be concerning possible random effects. Possible random effects are:

- ID
- trial (potentially nested in date)
- observer

It is a common “problem” in ‘Stan’ models (as implemented in `brms`) that random effects (group-level effects) with only few levels lead to divergent transitions (i.e. the model basically fails converging). This problem becomes apparent for `observer` as well as if we nest `date_trial` within `data`. Although `date_trial` and `date` have a lot of different levels, if we use the nested design, we have a maximum of 5 levels of `date_trial` per group-level `date`.

We use the full model (type and both log-illuminance measures as well as all interactions as fixed effects) to test what `brms` tells us when including certain random effects.

For starters, we include ID, trial nested in date and observer as random effects.

Note that we are setting a seed within the `brm` function.

```
mod1.0 <- suppressMessages(brm(court_lin_heu_01 ~ type * log_light_HC.z * 
    log_light_TLC.z + (1 | ID) + (1 | date/date_trial) + (1 | 
    observer), prior = c(set_prior("normal(0,3)", class = "b", 
    coef = "type1"), set_prior("normal(0,3)", class = "Intercept")), 
    family = "bernoulli", data = pref_stat_TLC_HC, chains = 5, 
    iter = 6000, warmup = 3000, refresh = 0, silent = TRUE, seed = 1))
```

```
## Warning: Rows containing NAs were excluded from the model.
```

```
## Warning: There were 203 divergent transitions after warmup. Increasing adapt_delta above 0.8 may help. See
## http://mc-stan.org/misc/warnings.html#divergent-transitions-after-warmup
```

```
## Warning: Examine the pairs() plot to diagnose sampling problems
```

```
## Warning: Tail Effective Samples Size (ESS) is too low, indicating posterior variances and tail quantiles may be unreliable.
## Running the chains for more iterations may help. See
## http://mc-stan.org/misc/warnings.html#tail-ess
```

We get a warning from Stan about divergent transaitions. Looking through the `Rhat`, `Bulk_ESS` and `Tail_ESS` estimates, we can see that `date` as random effect seems to cause most trouble. We see a relatively big `Rhat` (`Rhat=1` would indicate convergence) and small ESS values. (If we’d run the same model as a GLMM with the `lme4` package, we could see that the variance explained by `date` alone is very close to 0.)

```
suppressWarnings(summary(mod1.0)$random$date)
```

```
##                Estimate Est.Error   l-95% CI  u-95% CI     Rhat Bulk_ESS
## sd(Intercept) 0.3509888 0.2147209 0.01846039 0.7867553 1.006304      775
##               Tail_ESS
## sd(Intercept)     2221
```

We exclude `date` and only keep `date_trial` as random (no more nested).

```
mod1.1 <- suppressMessages(update(mod1.0, formula. = ~. + (1 | 
    date_trial) - (1 | date/date_trial), refresh = 0, silent = TRUE, 
    seed = 2))
```

```
## Warning: There were 205 divergent transitions after warmup. Increasing adapt_delta above 0.8 may help. See
## http://mc-stan.org/misc/warnings.html#divergent-transitions-after-warmup
```

```
## Warning: Examine the pairs() plot to diagnose sampling problems
```

We still get lots of warnings about divergent transitions. Looking again at diagnostics about the model, we can see that also `observer` has a quite high `Rhat` and comparibly weak EES values (again, we would see the same in a frequentist GLMM).

```
suppressWarnings(summary(mod1.1)$random$observer)
```

```
##                Estimate Est.Error   l-95% CI u-95% CI     Rhat Bulk_ESS
## sd(Intercept) 0.5414252 0.4045074 0.04984416 1.639833 1.002506     2099
##               Tail_ESS
## sd(Intercept)     1918
```

We discard it as well.

```
mod1.2 <- suppressMessages(update(mod1.1, formula. = ~. - (1 | 
    observer), refresh = 0, silent = TRUE, seed = 3))
```

No more warnings. The model seems to converge fine. We build another model reduced by `date_trial` as random effect to see how important it is.

```
mod1.3 <- suppressMessages(update(mod1.2, formula. = ~. - (1 | 
    date_trial), refresh = 0, silent = TRUE, seed = 4))
```

Perform approximate leave-one-out (LOO) cross-validation with `mod1.2` and `mod1.3.`

```
loo_compare(list(loo(mod1.2), loo(mod1.3)))
```

```
##        elpd_diff se_diff
## mod1.2    0.0       0.0 
## mod1.3 -207.7      19.0
```

Comparing ELPD and SE, we can see that `mod1.2` performs tremendously better! We find a massive ELPD-difference between the models, many folds higher than the standard error of ELPD. `date_trial` should be kept as random factor.

We definitely want to keep `ID` as random effect as well to correct for pseudoreplication/overdispersion. Therefore we are settled on the random effects and chose `mod1.2` as blueprint for random effects design.

###### 2.2.1.1.2 Fixed effects

Now we check which fixed effects we want to use. We try out all possible (nested/reduced) variations of the model and then use the *WAIC* (and additionally also *LOOIC*) information criterion to assess model fit.

Because we have some `NA` observations in the log-illuminance columns, we have to create a reduced dataset, where columns with `NA` get removed. This will allow that all models will be fitted to the same dataset (otherwise, due to different involved columns, some `NA` columns will be dropped for one model, but not for the other).

```
pref_stat_TLC_HC_red <- pref_stat_TLC_HC[!is.na(pref_stat_TLC_HC$log_light_HC.z) & 
    !is.na(pref_stat_TLC_HC$log_light_TLC.z), ]
# For safety, we drop levels (may cause some problems if we
# don't)
pref_stat_TLC_HC_red <- droplevels(pref_stat_TLC_HC_red)
```

Create reduced version of `mod1.2` (which should actually be exactly the same as `mod1.2`, since `NA`s are anyway discarded during modelling). We need to do this step so we can fit the model to the new `pref_stat_TLC_HC_red` dataset. We will below update this model (avoids compiling time), so we have to make sure it’s fitted to the new dataset.

```
mod1.2_red <- suppressMessages(brm(court_lin_heu_01 ~ type * 
    log_light_HC.z * log_light_TLC.z + (1 | ID) + (1 | date_trial), 
    prior = c(set_prior("normal(0,3)", class = "b", coef = "type1"), 
        set_prior("normal(0,3)", class = "Intercept")), family = "bernoulli", 
    data = pref_stat_TLC_HC_red, chains = 5, iter = 6000, warmup = 3000, 
    refresh = 0, silent = TRUE, seed = 5))
```

Now we run all the reduced versions of the full model.

```
# The null model (all fixed effects and interactions removed)
mod1.200 <- suppressMessages(update(mod1.2_red, formula. = ~. - 
    type - log_light_HC.z - log_light_TLC.z - type:log_light_HC.z - 
    type:log_light_TLC.z - log_light_HC.z:log_light_TLC.z - type:log_light_HC.z:log_light_TLC.z, 
    refresh = 0, silent = TRUE, seed = 6))
# The model only including type as fixed
mod1.201 <- suppressMessages(update(mod1.2_red, formula. = ~. - 
    log_light_HC.z - log_light_TLC.z - type:log_light_HC.z - 
    type:log_light_TLC.z - log_light_HC.z:log_light_TLC.z - type:log_light_HC.z:log_light_TLC.z, 
    refresh = 0, silent = TRUE, seed = 7))
# The model only including log-illuminance at heu 'female'
mod1.202 <- suppressMessages(update(mod1.2_red, formula. = ~. - 
    type - log_light_TLC.z - type:log_light_HC.z - type:log_light_TLC.z - 
    log_light_HC.z:log_light_TLC.z - type:log_light_HC.z:log_light_TLC.z, 
    refresh = 0, silent = TRUE, seed = 8))
# The model only including log-illuminance at lin 'female'
mod1.203 <- suppressMessages(update(mod1.2_red, formula. = ~. - 
    type - log_light_HC.z - type:log_light_HC.z - type:log_light_TLC.z - 
    log_light_HC.z:log_light_TLC.z - type:log_light_HC.z:log_light_TLC.z, 
    refresh = 0, silent = TRUE, seed = 9))
# The model including type and log-illuminance at heu
# 'female'
mod1.204 <- suppressMessages(update(mod1.2_red, formula. = ~. - 
    log_light_TLC.z - type:log_light_HC.z - type:log_light_TLC.z - 
    log_light_HC.z:log_light_TLC.z - type:log_light_HC.z:log_light_TLC.z, 
    refresh = 0, silent = TRUE, seed = 10))
# The model including type and log-illuminance at lin
# 'female'
mod1.205 <- suppressMessages(update(mod1.2_red, formula. = ~. - 
    log_light_HC.z - type:log_light_HC.z - type:log_light_TLC.z - 
    log_light_HC.z:log_light_TLC.z - type:log_light_HC.z:log_light_TLC.z, 
    refresh = 0, silent = TRUE, seed = 11))
# The model including log-illuminance at heu 'female' and
# log-illuminance at lin 'female'
mod1.206 <- suppressMessages(update(mod1.2_red, formula. = ~. - 
    type - type:log_light_HC.z - type:log_light_TLC.z - log_light_HC.z:log_light_TLC.z - 
    type:log_light_HC.z:log_light_TLC.z, refresh = 0, silent = TRUE, 
    seed = 12))
# The model including type, log-illuminance at heu 'female'
# and log-illuminance at lin 'female'
mod1.207 <- suppressMessages(update(mod1.2_red, formula. = ~. - 
    type:log_light_HC.z - type:log_light_TLC.z - log_light_HC.z:log_light_TLC.z - 
    type:log_light_HC.z:log_light_TLC.z, refresh = 0, silent = TRUE, 
    seed = 13))
# The model including type, log-illuminance at heu 'female'
# and their interaction
mod1.208 <- suppressMessages(update(mod1.2_red, formula. = ~. - 
    log_light_TLC.z - type:log_light_TLC.z - log_light_HC.z:log_light_TLC.z - 
    type:log_light_HC.z:log_light_TLC.z, refresh = 0, silent = TRUE, 
    seed = 14))
# The model including type, log-illuminance at lin 'female'
# and their interaction
mod1.209 <- suppressMessages(update(mod1.2_red, formula. = ~. - 
    log_light_HC.z - type:log_light_HC.z - log_light_HC.z:log_light_TLC.z - 
    type:log_light_HC.z:log_light_TLC.z, refresh = 0, silent = TRUE, 
    seed = 15))
# The model including log-illuminance at heu 'female',
# log-illuminance at lin 'female' and their interaction
mod1.210 <- suppressMessages(update(mod1.2_red, formula. = ~. - 
    type - type:log_light_HC.z - type:log_light_TLC.z - type:log_light_HC.z:log_light_TLC.z, 
    refresh = 0, silent = TRUE, seed = 16))
# The model including type, log-illuminance at heu 'female',
# log-illuminance at lin 'female' and the interaction between
# type and log-illuminance at heu 'female'
mod1.211 <- suppressMessages(update(mod1.2_red, formula. = ~. - 
    type:log_light_TLC.z - log_light_HC.z:log_light_TLC.z - type:log_light_HC.z:log_light_TLC.z, 
    refresh = 0, silent = TRUE, seed = 17))
# The model including type, log-illuminance at heu 'female',
# log-illuminance at lin 'female' and the interaction between
# type and log-illuminance at lin 'female'
mod1.212 <- suppressMessages(update(mod1.2_red, formula. = ~. - 
    type:log_light_HC.z - log_light_HC.z:log_light_TLC.z - type:log_light_HC.z:log_light_TLC.z, 
    refresh = 0, silent = TRUE, seed = 18))
# The model including type, log-illuminance at heu 'female',
# log-illuminance at lin 'female' and the interaction between
# the two log-illuminance measures
mod1.213 <- suppressMessages(update(mod1.2_red, formula. = ~. - 
    type:log_light_HC.z - type:log_light_TLC.z - type:log_light_HC.z:log_light_TLC.z, 
    refresh = 0, silent = TRUE, seed = 19))
# The model including type, log-illuminance at heu 'female',
# log-illuminance at lin 'female', the interaction between
# type and log-illuminance at heu 'female' and the
# interaction between the two log-illuminance measures
mod1.214 <- suppressMessages(update(mod1.2_red, formula. = ~. - 
    type:log_light_TLC.z - type:log_light_HC.z:log_light_TLC.z, 
    refresh = 0, silent = TRUE, seed = 20))
# The model including type, log-illuminance at heu 'female',
# log-illuminance at lin 'female', the interaction between
# type and log-illuminance at lin 'female' and the
# interaction between the two log-illuminance measures
mod1.215 <- suppressMessages(update(mod1.2_red, formula. = ~. - 
    type:log_light_HC.z - type:log_light_HC.z:log_light_TLC.z, 
    refresh = 0, silent = TRUE, seed = 21))
# The model including type, log-illuminance at heu 'female',
# log-illuminance at lin 'female', the interaction between
# type and log-illuminance at heu 'female' and the
# interaction between type and log-illuminance at lin
# 'female'
mod1.216 <- suppressMessages(update(mod1.2_red, formula. = ~. - 
    log_light_HC.z:log_light_TLC.z - type:log_light_HC.z:log_light_TLC.z, 
    refresh = 0, silent = TRUE, seed = 22))
# The model including type, log-illuminance at heu 'female',
# log-illuminance at lin 'female' and all two-way
# interactions
mod1.217 <- suppressMessages(update(mod1.2_red, formula. = ~. - 
    type:log_light_HC.z:log_light_TLC.z, refresh = 0, silent = TRUE, 
    seed = 23))
```

Calculate LOOIC and WAIC for `mod1.2_red` and all nested, reduced models.

```
all_looic <- c(loo(mod1.2_red)$estimates[3, 1], sapply(0:17, 
    function(x) loo(get(paste0("mod1.2", ifelse(x < 10, paste0("0", 
        x), x))))$estimates[3, 1]))
all_waic <- c(waic(mod1.2_red)$estimates[3, 1], sapply(0:17, 
    function(x) waic(get(paste0("mod1.2", ifelse(x < 10, paste0("0", 
        x), x))))$estimates[3, 1]))
```

To create model weights based on WAIC, we use the exact same code as the `model_weights` function of `brms` would do. We calculate it here manually, since the `model_weights` function would otherwise calculate all WAIC values again (which would cost time and we’d also have to be careful with settings seeds, since WAIC calculation relies on the random number generator).

```
ic_diffs <- all_waic - min(all_waic)
out <- exp(-ic_diffs/2)
out <- as.numeric(out)
out <- out/sum(out)
```

We summarize LOOIC, WAIC and WAIC weights in a table, and additionally calculate WAIC differences between best fitting model and the other models. We then sort this table by WAIC value and view it.

```
IC_table <- data.frame(Model = c("mod1.2_red", paste0("mod1.2", 
    ifelse((0:17) < 10, paste0("0", 0:17), 0:17))), LOOIC = all_looic, 
    WAIC = all_waic, WAIC_Diff = ic_diffs, WAIC_weight = out)
IC_table <- IC_table[order(IC_table$WAIC), ]
cbind(Model = IC_table[, 1], round(IC_table[, -1], 2))
```

We can see that the six models, which include the interaction term between `type` and log-illuminance at the *heu* ‘female’ are the six best fitting models. If we add all their model weights up, we can see that these models have a high weight together:

```
round(sum(IC_table$WAIC_weight[1:6]), 2)
```

```
## [1] 0.8
```

We can see that the model with the best fit is `mod1.208`, which includes `type`, log-illuminance at *heu* ‘female’ and their interaction.

We safe this table:

```
write.csv(IC_table, "Tables/WAIC_pure.csv", row.names = F, quote = F)
```

We therefore settle for `mod1.208`. We fit the model again, but now to the full dataset (this should result in almost the same model, since almost all `NA` rows were shared between `log_light_HC.z` and `log_light_TLC.z`). We fit it explicitely here (without the `update()` function), so it’s better visible how the model looks like. We use the same seed as before (which though shouldn’t play any role, since we use a new dataset).

```
mod1.208 <- suppressMessages(brm(court_lin_heu_01 ~ type * log_light_HC.z + 
    (1 | ID) + (1 | date_trial), prior = c(set_prior("normal(0,3)", 
    class = "b", coef = "type1"), set_prior("normal(0,3)", class = "Intercept")), 
    family = "bernoulli", data = pref_stat_TLC_HC, chains = 5, 
    iter = 6000, warmup = 3000, refresh = 0, silent = TRUE, seed = 14))
```

```
## Warning: Rows containing NAs were excluded from the model.
```

For visualization purposes later, we use the same model structure to generate a new “categorical” illuminance model, where we split log-illuminance at the *heu* model into dark and bright, following the median of the dataset. We have to add a new column for that, putting rows into each of the two classes.

```
pref_stat_TLC_HC$quantile_low_up <- as.integer(cut(pref_stat_TLC_HC$log_light_HC.z, 
    quantile(pref_stat_TLC_HC$log_light_HC.z, probs = c(0, 0.5, 
        1), na.rm = T), include.lowest = TRUE))
pref_stat_TLC_HC$quantile_low_up <- paste0("q", pref_stat_TLC_HC$quantile_low_up)
```

We run the categorical model (we again use the same seed as before, but it again doesn’t matter at all really).

```
# Exclude NA measurements
mod1.208_cat <- suppressMessages(brm(court_lin_heu_01 ~ type * 
    quantile_low_up + (1 | ID) + (1 | date_trial), prior = c(set_prior("normal(0,3)", 
    class = "b", coef = "type1"), set_prior("normal(0,3)", class = "Intercept")), 
    family = "bernoulli", data = pref_stat_TLC_HC[pref_stat_TLC_HC$quantile_low_up != 
        "qNA", ], chains = 5, iter = 6000, warmup = 3000, refresh = 0, 
    silent = TRUE, seed = 14))
```

We return back to the main model.

Let’s do some quick model checking. We see “hairy caterpillars” in the trace plots on the right, which is a good sign (the MCMC algorithm covered parameter space well)!

```
plot(mod1.208)
```

Some posterior predictive checks to compare model predictions to actual data. Looks like a good fit!

```
suppressMessages(pp_check(mod1.208))
```

Now let’s have a quick look at the model coefficients. Annotation of `type` is numeric due to `contr.sum()` dummy coding:

```
fixef(mod1.208)[, 1]
```

```
##            Intercept                type1       log_light_HC.z 
##          -0.07704699           0.13759589           0.06539505 
## type1:log_light_HC.z 
##           0.11800715
```

Next, we want to look at model estimators (estimated marginal means) and contrasts. We use the awesome `emmeans` package to extract posteriors for estimators and contrasts.

For this, we first define two functions which will help handle the `emmeans` output.

This function transforms the posteriors stored in an `emmeans` output to a more user-friendly table. This function does essentially the same as `as.mcmc.emmGrid` from `emmeans` or `gather_emmeans_draws` from the `tidybayes` package. This just produces a handier format, as we think (all values below each other with an identifier column, as in `gather_emmeans_draws`, but with ‘non-tidy’ format):

```
extract_emm_posterior <- function(emm_object) {
    
    # Get parameter combinations
    level_comb <- expand.grid(emm_object@levels, stringsAsFactors = F)
    # Get number of elements per posterior
    iterat <- nrow(emm_object@post.beta)
    
    # We need to do matrix multiplication of the posterior slot
    # to get the posteriors relating to the emmeans output (in
    # some cases, this step is not necessary, but it doesn't
    # matter to do it anyways):
    slopes <- emm_object@post.beta %*% t(emm_object@linfct)
    
    # Append each posterior and give parameter combination and
    # iteration ID
    post_out <- data.frame(level_comb[rep(seq_len(nrow(level_comb)), 
        each = iterat), , drop = F], iteration = rep(1:iterat, 
        nrow(level_comb)), value = as.vector(slopes), stringsAsFactors = F)
    
    # Remove row names and return
    rownames(post_out) <- NULL
    return(post_out)
}
```

This function calculates a credible interval and a point estimate for different posteriors from an `emmeans` output. `emmeans` does already return such summaries, but it defaults to pass the median and the 95% high-density interval of a posterior. This function allows to extract instead the mean (decided by `middle` argument) and/or the 95% equal-tailed interval (decided by `type0.95` argument), if wished.

```
extract_emm_cri <- function(emm_object, middle = "mean", type0.95 = "eti") {
    
    # Get parameter combinations
    level_comb <- expand.grid(emm_object@levels, stringsAsFactors = F)
    
    # We need to do matrix multiplication of the posterior slot
    # to get the posteriors relating to the emmeans output (in
    # some cases, this step is not necessary):
    slopes <- emm_object@post.beta %*% t(emm_object@linfct)
    # If equal-tailed interval, calculate this for posteriors of
    # each parameter combination
    if (type0.95 == "eti") {
        summ_estimators <- data.frame(level_comb, t(sapply(1:prod(lengths(emm_object@levels)), 
            function(x) c(eval(parse(text = paste0(middle, "(slopes[,x])"))), 
                quantile(slopes[, x], probs = c(0.025, 0.975))))))
        # If not equal-tailed interval, calculate high-density
        # interval for posteriors of each parameter combination
    } else {
        # For this, we have to load the HDInterval library (this will
        # actually never happen in this Markdown, which is why we
        # didn't do so in the beginning).
        if (!"HDInterval" %in% rownames(installed.packages())) {
            install.packages("HDInterval")
        }
        suppressMessages(suppressWarnings(library(HDInterval)))
        
        summ_estimators <- data.frame(level_comb, t(sapply(1:prod(lengths(emm_object@levels)), 
            function(x) c(eval(parse(text = paste0(middle, "(slopes[,x])"))), 
                hdi(slopes[, x])))))
    }
    
    # Give names to the new table and return
    names(summ_estimators) <- c(names(emm_object@levels), "estimate", 
        "lower", "upper")
    return(summ_estimators)
}
```

Let’s look at the credible intervals for `type` alone. We first create an `emmeans` object.

```
group_est_main <- suppressWarnings(suppressMessages(emmeans(mod1.208, 
    ~type, transform = "response")))
```

We could call the `extract_emm_cri` function we just created; but for now, we actually also need the posteriors, so we do this manually. We extract the posteriors.

```
group_posteriors_main <- extract_emm_posterior(group_est_main)
```

Split posteriors by type

```
TLC_posterior_main <- group_posteriors_main$value[group_posteriors_main$type == 
    "TLC"]
HC_posterior_main <- group_posteriors_main$value[group_posteriors_main$type == 
    "HC"]
```

Create “difference posterior”

```
difference_posterior_main <- HC_posterior_main - TLC_posterior_main
```

Now we take a look at mean and 95% CrI for *lin*, *heu* and the difference between them.

```
est_types <- data.frame(c("TLC", "HC", "HC-TLC"), rbind(c(mean(TLC_posterior_main), 
    quantile(TLC_posterior_main, probs = c(0.025, 0.975))), c(mean(HC_posterior_main), 
    quantile(HC_posterior_main, probs = c(0.025, 0.975))), c(mean(difference_posterior_main), 
    quantile(difference_posterior_main, probs = c(0.025, 0.975)))))
names(est_types) <- c("object", "mean", "lower", "upper")
print(est_types)
```

```
##   object       mean        lower     upper
## 1    TLC 0.44677856  0.383358838 0.5128524
## 2     HC 0.51506008  0.448606834 0.5817226
## 3 HC-TLC 0.06828152 -0.003130615 0.1404361
```

We see an effect size of about 0.07.

Now we test how likely it is that the predictors for *lin* and *heu* differ. We test three hypotheses:

1. *lin* males have a smaller preference score than *heu* males.
2. *heu* males show a preference for their own type.
3. *lin* males show a preference for their own type.

While hypothesis 1) is of high interest, 2) and 3) don’t matter very much. 0.5 is a meaningless “null-hypothesis”, which doesn’tmake much sense when testing males with dead, mounted females. What matters is that their visual attraction diverges!

```
print(data.frame(test = c("HC-TLC > 0", "HC > 0", "TLC < 0"), 
    posterior_prop = c(sum(difference_posterior_main > 0)/length(difference_posterior_main), 
        sum(HC_posterior_main > 0.5)/length(HC_posterior_main), 
        sum(TLC_posterior_main < 0.5)/length(TLC_posterior_main))))
```

```
##         test posterior_prop
## 1 HC-TLC > 0      0.9694000
## 2     HC > 0      0.6662667
## 3    TLC < 0      0.9438667
```

We can see that the posterior overlap of the two predictors is small! Therefore it is very probable that the preference of the two male types differ. The probability for *heu* males to have a preference for their own type (>0.5) is smaller than for *lin* males (<0.5).

Now, let’s look at the **simple** slopes of the interaction terms between `type` and log-illuminance at the *heu* ‘female’.

This gets us the ‘simple slopes’ from the interaction term. We use the `emtrends` function for that.

```
group_est_main_interact <- suppressWarnings(suppressMessages(emtrends(mod1.208, 
    ~type, var = "log_light_HC.z", transform = "response")))
```

Note: this is not the true slope at all points along the gradient. The interaction term is in fact not linear. Retrieving the **simple** slope is the general solution to describe the ‘trend’ of an interaction. Also, this slope is quite hard to interpret (better is the vsualization, which follows later), as it refers to the centered&scaled illuminance measures. What matters to us now is into which direction these two slopes differ from 0 and if they are different from each other.

We print the mean and 95% equal-tailed interval for these slopes.

```
print(extract_emm_cri(group_est_main_interact))
```

```
##   type    estimate        lower      upper
## 1   HC  0.04558845  0.009779743 0.08210984
## 2  TLC -0.01293356 -0.047578276 0.02238481
```

We get the posteriors for the simple slopes

```
group_posteriors_main_interact <- extract_emm_posterior(group_est_main_interact)
```

We use the same hypotheses as before. In the interaction term context, they mean something slightly different though.

1. The slope of the interaction term between *lin* and log-illuminance at *heu* ‘female’ is smaller than that of the interaction term between *heu* and log-illuminance at *heu* ‘female’.
2. Increase in log-illuminance around the *heu* ‘female’ leads to an increase in preference of *heu* males for their own type (positive slope).
3. Increase in log-illuminance around the *heu* ‘female’ leads to an increase in preference of *lin* males for their own type (negative slope).

```
iterations <- nrow(group_posteriors_main_interact)/2
print(data.frame(test = c("HC > TLC", "HC > 0", "TLC < 0"), posterior_prob = c(sum(group_posteriors_main_interact$value[group_posteriors_main_interact$type == 
    "HC"] > group_posteriors_main_interact$value[group_posteriors_main_interact$type == 
    "TLC"])/iterations, sum(group_posteriors_main_interact$value[group_posteriors_main_interact$type == 
    "HC"] > 0)/iterations, sum(group_posteriors_main_interact$value[group_posteriors_main_interact$type == 
    "TLC"] < 0)/iterations)))
```

```
##       test posterior_prob
## 1 HC > TLC      0.9960000
## 2   HC > 0      0.9934000
## 3  TLC < 0      0.7683333
```

We can see that with a high probability, the slopes of the two interaction terms differ. The posteriors barely overlap. The slope for *heu* males is with very high probability positive. The slope of *lin* males is negative with quite high probability, but the effect is less clear.

We repeat a similar procedure with the categorical illuminace model, `mod1.208_cat`.

Extract the posteriors for all levels of type from the categorical model, under varying categorical illumination at *heu* model

```
group_est_cat <- suppressWarnings(suppressMessages(emmeans(mod1.208_cat, 
    ~type | quantile_low_up, transform = "response")))
```

We extract the posteriors.

```
group_posteriors_cat <- extract_emm_posterior(group_est_cat)
```

Split posteriors by type and illuminance

```
TLC_posterior_cat_1 <- group_posteriors_cat$value[group_posteriors_cat$type == 
    "TLC" & group_posteriors_cat$quantile_low_up == "q1"]
HC_posterior_cat_1 <- group_posteriors_cat$value[group_posteriors_cat$type == 
    "HC" & group_posteriors_cat$quantile_low_up == "q1"]
TLC_posterior_cat_2 <- group_posteriors_cat$value[group_posteriors_cat$type == 
    "TLC" & group_posteriors_cat$quantile_low_up == "q2"]
HC_posterior_cat_2 <- group_posteriors_cat$value[group_posteriors_cat$type == 
    "HC" & group_posteriors_cat$quantile_low_up == "q2"]
```

Create “difference posteriors”

```
difference_posterior_cat_1 <- HC_posterior_cat_1 - TLC_posterior_cat_1
difference_posterior_cat_2 <- HC_posterior_cat_2 - TLC_posterior_cat_2
```

Now we take a look at mean and 95% CrI for *lin*, *heu* and the difference between them.

```
est_types <- data.frame(rep(c("TLC", "HC", "HC-TLC"), 2), rep(c("dark", 
    "bright"), each = 3), rbind(c(mean(TLC_posterior_cat_1), 
    quantile(TLC_posterior_cat_1, probs = c(0.025, 0.975))), 
    c(mean(HC_posterior_cat_1), quantile(HC_posterior_cat_1, 
        probs = c(0.025, 0.975))), c(mean(difference_posterior_cat_1), 
        quantile(difference_posterior_cat_1, probs = c(0.025, 
            0.975))), c(mean(TLC_posterior_cat_2), quantile(TLC_posterior_cat_2, 
        probs = c(0.025, 0.975))), c(mean(HC_posterior_cat_2), 
        quantile(HC_posterior_cat_2, probs = c(0.025, 0.975))), 
    c(mean(difference_posterior_cat_2), quantile(difference_posterior_cat_2, 
        probs = c(0.025, 0.975)))))
names(est_types) <- c("object", "light_env_heu", "mean", "lower", 
    "upper")
print(est_types)
```

```
##   object light_env_heu       mean       lower      upper
## 1    TLC          dark 0.45564920  0.38507409 0.52745617
## 2     HC          dark 0.46375815  0.39124146 0.54010027
## 3 HC-TLC          dark 0.00810895 -0.07363075 0.09293149
## 4    TLC        bright 0.43732555  0.36199240 0.51429536
## 5     HC        bright 0.57151034  0.49267676 0.64921957
## 6 HC-TLC        bright 0.13418479  0.05093610 0.21869133
```

Looks pretty similar to the continuous model!

To check if it is really log-illuminance at the *heu* ‘female’ and not log-illuminance in general driving this difference, we now test the effect of the two log-illuminance measures from the full model `mod1.2`. This model includes `type`, and log-illuminance measures at the two mounted females, as well as all possible interactions. We now only look at the two-way interactions; for a visualization of the three-way interaction, see later.

The description of the following is reduced to the minimum, since it’s basically a repetition of what we just did.

First, we repeat what we just did for the main model, and estimate the interaction term with log-illuminance at the *heu* model.

```
group_est_main_interact <- suppressWarnings(suppressMessages(emtrends(mod1.2, 
    ~type, var = "log_light_HC.z", transform = "response")))
group_posteriors_main_interact <- extract_emm_posterior(group_est_main_interact)
iterations <- nrow(group_posteriors_main_interact)/2
print(data.frame(test = c("HC > TLC", "HC > 0", "TLC < 0"), posterior_prob = c(sum(group_posteriors_main_interact$value[group_posteriors_main_interact$type == 
    "HC"] > group_posteriors_main_interact$value[group_posteriors_main_interact$type == 
    "TLC"])/iterations, sum(group_posteriors_main_interact$value[group_posteriors_main_interact$type == 
    "HC"] > 0)/iterations, sum(group_posteriors_main_interact$value[group_posteriors_main_interact$type == 
    "TLC"] < 0)/iterations)))
```

```
##       test posterior_prob
## 1 HC > TLC      0.9967333
## 2   HC > 0      0.9912000
## 3  TLC < 0      0.8304667
```

Looks the same. Now we repeat the same for the interaction term with log-illuminance at the *lin* model. We use the same hypotheses, i.e. that more intense light on the ‘female’ increase preference for own female type in both male types, and that both responses are in opposite directions.

```
group_est_main_interact <- suppressWarnings(suppressMessages(emtrends(mod1.2, 
    ~type, var = "log_light_TLC.z", transform = "response")))
group_posteriors_main_interact <- extract_emm_posterior(group_est_main_interact)
iterations <- nrow(group_posteriors_main_interact)/2
print(data.frame(test = c("HC > TLC", "HC > 0", "TLC < 0"), posterior_prob = c(sum(group_posteriors_main_interact$value[group_posteriors_main_interact$type == 
    "HC"] > group_posteriors_main_interact$value[group_posteriors_main_interact$type == 
    "TLC"])/iterations, sum(group_posteriors_main_interact$value[group_posteriors_main_interact$type == 
    "HC"] > 0)/iterations, sum(group_posteriors_main_interact$value[group_posteriors_main_interact$type == 
    "TLC"] < 0)/iterations)))
```

```
##       test posterior_prob
## 1 HC > TLC      0.1902667
## 2   HC > 0      0.3918667
## 3  TLC < 0      0.1940000
```

We see very weak support for log-illuminance at the *lin* ‘female’ having much of an effect (posterior probabilities are neither close to 1, nor to 0). It’s clear that light at the *lin* ‘female’ seems to not affect preference of either type very much.

###### 2.2.1.2 Effect of presence/absence of red colour in backcross hybrids and its interaction with illuminance measures on visual mate preference

For the backcross-to-*lin* males, we basically follow the same pipeline as before, just that instead of `type` (species), we now use presence/absence of red colour (`redYN` = genotpye at optix region) on the male wings as fixed effect.

We set the same weak priors for genotype as we did before for `type`.

We encountered the exact same problems when including `date` (with `date_trial` nested) and `observer` as random effects. We won’t show this here, but start right away with the same type of model structure as `mod1.2` above.

Since we can track hybrid broods, we decide to include `broods` nested in `exp_line` as random effect. Remember: `broods` explains relationships at the sibling-level, while `exp_line` explains relationship at the grandchildren-level. Since the model gets more complex, we increase the `adapt_delta` parameter, which makes the MCMC algorith taking “smaller steps” and therefore avoiding convergence issues.

```
mod2.2_with_exp_line <- suppressMessages(brm(court_lin_heu_01 ~ 
    redYN * log_light_HC.z * log_light_TLC.z + (1 | ID) + (1 | 
        date_trial) + (1 | exp_line/broods), prior = c(set_prior("normal(0,3)", 
    class = "b", coef = "redYN1"), set_prior("normal(0,3)", class = "Intercept")), 
    family = "bernoulli", data = pref_stat_BC, chains = 5, iter = 6000, 
    warmup = 3000, refresh = 0, silent = TRUE, seed = 24, control = list(adapt_delta = 0.99)))
```

```
## Warning: Rows containing NAs were excluded from the model.
```

```
## Warning: There were 5 divergent transitions after warmup. Increasing adapt_delta above 0.99 may help. See
## http://mc-stan.org/misc/warnings.html#divergent-transitions-after-warmup
```

```
## Warning: Examine the pairs() plot to diagnose sampling problems
```

We can see quickly that `exp_line` is causing trouble. Our model does not converge well because we have too many group-level states with often too few observations per individual level.

Looking at the model summary, we see that we get a similar warning as above concerning divergent transitions. This is not a good sign. We even get one remarkably bad `Rhat` and ESS measures.

```
suppressWarnings(summary(mod2.2_with_exp_line)$random$exp_line)
```

```
##               Estimate Est.Error   l-95% CI u-95% CI     Rhat Bulk_ESS Tail_ESS
## sd(Intercept)  0.52464  0.593141 0.04178682 2.197386 1.000648     4490     6278
```

… we can also see this we extract variance and correlation components (the estimation error is very high).

```
VarCorr(mod2.2_with_exp_line)
```

```
## $date_trial
## $date_trial$sd
##            Estimate  Est.Error      Q2.5    Q97.5
## Intercept 0.9248998 0.06328796 0.8069704 1.055001
## 
## 
## $exp_line
## $exp_line$sd
##           Estimate Est.Error       Q2.5    Q97.5
## Intercept  0.52464  0.593141 0.04178682 2.197386
## 
## 
## $`exp_line:broods`
## $`exp_line:broods`$sd
##             Estimate Est.Error        Q2.5     Q97.5
## Intercept 0.07884123  0.062205 0.002946977 0.2328851
## 
## 
## $ID
## $ID$sd
##            Estimate  Est.Error      Q2.5     Q97.5
## Intercept 0.3487939 0.05355487 0.2464474 0.4579797
```

Therefore, we remove `exp_line` from our model, but instead look separately at the different experimental lines later (see below). We create a new model without `exp_line`.

```
mod2.2 <- suppressMessages(update(mod2.2_with_exp_line, formula. = ~. - 
    (1 | exp_line/broods) + (1 | broods), refresh = 0, silent = TRUE, 
    seed = 25))
```

We again reduce the dataset to non-`NA`

```
pref_stat_BC_red <- pref_stat_BC[!is.na(pref_stat_BC$log_light_HC.z) & 
    !is.na(pref_stat_BC$log_light_TLC.z), ]
# For safety, we drop levels (seems to cause some problems if
# we don't)
pref_stat_BC_red <- droplevels(pref_stat_BC_red)
```

We build the reduced version of the model. For safety, we build it from scratch, so there is no problem with the adaptation of the precompiled model.

```
mod2.2_red <- suppressMessages(brm(court_lin_heu_01 ~ redYN * 
    log_light_HC.z * log_light_TLC.z + (1 | ID) + (1 | date_trial) + 
    (1 | broods), prior = c(set_prior("normal(0,3)", class = "b", 
    coef = "redYN1"), set_prior("normal(0,3)", class = "Intercept")), 
    family = "bernoulli", data = pref_stat_BC_red, chains = 5, 
    iter = 6000, warmup = 3000, refresh = 0, silent = TRUE, seed = 26, 
    control = list(adapt_delta = 0.99)))
```

Now we check which fixed effects we want to use. We again try out all possible combinations of the model.

```
# The null model (all fixed effects and interactions removed)
mod2.200 <- suppressMessages(update(mod2.2_red, formula. = ~. - 
    redYN - log_light_HC.z - log_light_TLC.z - redYN:log_light_HC.z - 
    redYN:log_light_TLC.z - log_light_HC.z:log_light_TLC.z - 
    redYN:log_light_HC.z:log_light_TLC.z, refresh = 0, silent = TRUE, 
    seed = 27))
# The model only including redYN as fixed
mod2.201 <- suppressMessages(update(mod2.2_red, formula. = ~. - 
    log_light_HC.z - log_light_TLC.z - redYN:log_light_HC.z - 
    redYN:log_light_TLC.z - log_light_HC.z:log_light_TLC.z - 
    redYN:log_light_HC.z:log_light_TLC.z, refresh = 0, silent = TRUE, 
    seed = 28))
# The model only including log-illuminance at heu 'female'
mod2.202 <- suppressMessages(update(mod2.2_red, formula. = ~. - 
    redYN - log_light_TLC.z - redYN:log_light_HC.z - redYN:log_light_TLC.z - 
    log_light_HC.z:log_light_TLC.z - redYN:log_light_HC.z:log_light_TLC.z, 
    refresh = 0, silent = TRUE, seed = 29))
# The model only including log-illuminance at lin 'female'
mod2.203 <- suppressMessages(update(mod2.2_red, formula. = ~. - 
    redYN - log_light_HC.z - redYN:log_light_HC.z - redYN:log_light_TLC.z - 
    log_light_HC.z:log_light_TLC.z - redYN:log_light_HC.z:log_light_TLC.z, 
    refresh = 0, silent = TRUE, seed = 30))
# The model including redYN and log-illuminance at heu
# 'female'
mod2.204 <- suppressMessages(update(mod2.2_red, formula. = ~. - 
    log_light_TLC.z - redYN:log_light_HC.z - redYN:log_light_TLC.z - 
    log_light_HC.z:log_light_TLC.z - redYN:log_light_HC.z:log_light_TLC.z, 
    refresh = 0, silent = TRUE, seed = 31))
# The model including redYN and log-illuminance at lin
# 'female'
mod2.205 <- suppressMessages(update(mod2.2_red, formula. = ~. - 
    log_light_HC.z - redYN:log_light_HC.z - redYN:log_light_TLC.z - 
    log_light_HC.z:log_light_TLC.z - redYN:log_light_HC.z:log_light_TLC.z, 
    refresh = 0, silent = TRUE, seed = 32))
# The model including log-illuminance at heu 'female' and
# log-illuminance at lin 'female'
mod2.206 <- suppressMessages(update(mod2.2_red, formula. = ~. - 
    redYN - redYN:log_light_HC.z - redYN:log_light_TLC.z - log_light_HC.z:log_light_TLC.z - 
    redYN:log_light_HC.z:log_light_TLC.z, refresh = 0, silent = TRUE, 
    seed = 33))
# The model including redYN, log-illuminance at heu 'female'
# and log-illuminance at lin 'female'
mod2.207 <- suppressMessages(update(mod2.2_red, formula. = ~. - 
    redYN:log_light_HC.z - redYN:log_light_TLC.z - log_light_HC.z:log_light_TLC.z - 
    redYN:log_light_HC.z:log_light_TLC.z, refresh = 0, silent = TRUE, 
    seed = 34))
# The model including redYN, log-illuminance at heu 'female'
# and their interaction
mod2.208 <- suppressMessages(update(mod2.2_red, formula. = ~. - 
    log_light_TLC.z - redYN:log_light_TLC.z - log_light_HC.z:log_light_TLC.z - 
    redYN:log_light_HC.z:log_light_TLC.z, refresh = 0, silent = TRUE, 
    seed = 35))
# The model including redYN, log-illuminance at lin 'female'
# and their interaction
mod2.209 <- suppressMessages(update(mod2.2_red, formula. = ~. - 
    log_light_HC.z - redYN:log_light_HC.z - log_light_HC.z:log_light_TLC.z - 
    redYN:log_light_HC.z:log_light_TLC.z, refresh = 0, silent = TRUE, 
    seed = 36))
# The model including log-illuminance at heu 'female',
# log-illuminance at lin 'female' and their interaction
mod2.210 <- suppressMessages(update(mod2.2_red, formula. = ~. - 
    redYN - redYN:log_light_HC.z - redYN:log_light_TLC.z - redYN:log_light_HC.z:log_light_TLC.z, 
    refresh = 0, silent = TRUE, seed = 37))
# The model including redYN, log-illuminance at heu 'female',
# log-illuminance at lin 'female' and the interaction between
# redYN and log-illuminance at heu 'female'
mod2.211 <- suppressMessages(update(mod2.2_red, formula. = ~. - 
    redYN:log_light_TLC.z - log_light_HC.z:log_light_TLC.z - 
    redYN:log_light_HC.z:log_light_TLC.z, refresh = 0, silent = TRUE, 
    seed = 38))
# The model including redYN, log-illuminance at heu 'female',
# log-illuminance at lin 'female' and the interaction between
# redYN and log-illuminance at lin 'female'
mod2.212 <- suppressMessages(update(mod2.2_red, formula. = ~. - 
    redYN:log_light_HC.z - log_light_HC.z:log_light_TLC.z - redYN:log_light_HC.z:log_light_TLC.z, 
    refresh = 0, silent = TRUE, seed = 39))
# The model including redYN, log-illuminance at heu 'female',
# log-illuminance at lin 'female' and the interaction between
# the two log-illuminance measures
mod2.213 <- suppressMessages(update(mod2.2_red, formula. = ~. - 
    redYN:log_light_HC.z - redYN:log_light_TLC.z - redYN:log_light_HC.z:log_light_TLC.z, 
    refresh = 0, silent = TRUE, seed = 40))
# The model including redYN, log-illuminance at heu 'female',
# log-illuminance at lin 'female', the interaction between
# redYN and log-illuminance at heu 'female' and the
# interaction between the two log-illuminance measures
mod2.214 <- suppressMessages(update(mod2.2_red, formula. = ~. - 
    redYN:log_light_TLC.z - redYN:log_light_HC.z:log_light_TLC.z, 
    refresh = 0, silent = TRUE, seed = 41))
# The model including redYN, log-illuminance at heu 'female',
# log-illuminance at lin 'female', the interaction between
# redYN and log-illuminance at lin 'female' and the
# interaction between the two log-illuminance measures
mod2.215 <- suppressMessages(update(mod2.2_red, formula. = ~. - 
    redYN:log_light_HC.z - redYN:log_light_HC.z:log_light_TLC.z, 
    refresh = 0, silent = TRUE, seed = 42))
# The model including redYN, log-illuminance at heu 'female',
# log-illuminance at lin 'female', the interaction between
# redYN and log-illuminance at heu 'female' and the
# interaction between redYN and log-illuminance at lin
# 'female'
mod2.216 <- suppressMessages(update(mod2.2_red, formula. = ~. - 
    log_light_HC.z:log_light_TLC.z - redYN:log_light_HC.z:log_light_TLC.z, 
    refresh = 0, silent = TRUE, seed = 43))
# The model including redYN, log-illuminance at heu 'female',
# log-illuminance at lin 'female' and all two-way
# interactions
mod2.217 <- suppressMessages(update(mod2.2_red, formula. = ~. - 
    redYN:log_light_HC.z:log_light_TLC.z, refresh = 0, silent = TRUE, 
    seed = 44))
```

Calculate LOOIC and WAIC for `mod2.2_red` and all nested, reduced models. We do this in a big chunk of code because (unlike for the pure males), these models are too big to allocate enough memory for running them all in one line.

```
all_looic <- rep(NA, 19)
all_looic[1] <- loo(mod2.2_red)$estimates[3, 1]
all_looic[2] <- loo(mod2.200)$estimates[3, 1]
all_looic[3] <- loo(mod2.201)$estimates[3, 1]
all_looic[4] <- loo(mod2.202)$estimates[3, 1]
all_looic[5] <- loo(mod2.203)$estimates[3, 1]
all_looic[6] <- loo(mod2.204)$estimates[3, 1]
all_looic[7] <- loo(mod2.205)$estimates[3, 1]
all_looic[8] <- loo(mod2.206)$estimates[3, 1]
all_looic[9] <- loo(mod2.207)$estimates[3, 1]
all_looic[10] <- loo(mod2.208)$estimates[3, 1]
all_looic[11] <- loo(mod2.209)$estimates[3, 1]
all_looic[12] <- loo(mod2.210)$estimates[3, 1]
all_looic[13] <- loo(mod2.211)$estimates[3, 1]
all_looic[14] <- loo(mod2.212)$estimates[3, 1]
all_looic[15] <- loo(mod2.213)$estimates[3, 1]
all_looic[16] <- loo(mod2.214)$estimates[3, 1]
all_looic[17] <- loo(mod2.215)$estimates[3, 1]
all_looic[18] <- loo(mod2.216)$estimates[3, 1]
all_looic[19] <- loo(mod2.217)$estimates[3, 1]

all_waic <- rep(NA, 19)
all_waic[1] <- waic(mod2.2_red)$estimates[3, 1]
all_waic[2] <- waic(mod2.200)$estimates[3, 1]
all_waic[3] <- waic(mod2.201)$estimates[3, 1]
all_waic[4] <- waic(mod2.202)$estimates[3, 1]
all_waic[5] <- waic(mod2.203)$estimates[3, 1]
all_waic[6] <- waic(mod2.204)$estimates[3, 1]
all_waic[7] <- waic(mod2.205)$estimates[3, 1]
all_waic[8] <- waic(mod2.206)$estimates[3, 1]
all_waic[9] <- waic(mod2.207)$estimates[3, 1]
all_waic[10] <- waic(mod2.208)$estimates[3, 1]
all_waic[11] <- waic(mod2.209)$estimates[3, 1]
all_waic[12] <- waic(mod2.210)$estimates[3, 1]
all_waic[13] <- waic(mod2.211)$estimates[3, 1]
all_waic[14] <- waic(mod2.212)$estimates[3, 1]
all_waic[15] <- waic(mod2.213)$estimates[3, 1]
all_waic[16] <- waic(mod2.214)$estimates[3, 1]
all_waic[17] <- waic(mod2.215)$estimates[3, 1]
all_waic[18] <- waic(mod2.216)$estimates[3, 1]
all_waic[19] <- waic(mod2.217)$estimates[3, 1]
```

To create model weights based on WAIC, we use the same method as above.

```
ic_diffs <- all_waic - min(all_waic)
out <- exp(-ic_diffs/2)
out <- as.numeric(out)
out <- out/sum(out)
```

We summarize LOOIC, WAIC and WAIC weights in a table, and additionally calculate WAIC differences between best fitting model and the other models. We then sort this table by WAIC value and view it.

```
IC_table <- data.frame(Model = c("mod2.2_red", paste0("mod2.2", 
    ifelse((0:17) < 10, paste0("0", 0:17), 0:17))), LOOIC = all_looic, 
    WAIC = all_waic, WAIC_Diff = ic_diffs, WAIC_weight = out)
IC_table <- IC_table[order(IC_table$WAIC), ]
cbind(Model = IC_table[, 1], round(IC_table[, -1], 2))
```

We can again see that the six models, which include the interaction term between `type` and log-illuminance at the *heu* ‘female’ are the six best fitting models. If we add all their model weights up, we see that these models have a high weight together:

```
round(sum(IC_table$WAIC_weight[1:6]), 2)
```

```
## [1] 0.91
```

We can see that the model with the best fit is `mod2.2_red`, which includes `type`, log-illuminance at *heu* ‘female’, log-illuminance at *lin* ‘female’ and possible interaction terms. We already generated this model structure fit to the full data set above, as `mod2.2`.

We safe this table:

```
write.csv(IC_table, "Tables/WAIC_backcross.csv", row.names = F, 
    quote = F)
```

For visualization purposes later, we use the same model structure to generate a new “categorical” illuminance model, where we split log-illuminance at the *heu* model and at the *lin* into dark and bright, following the median measures of the dataset. We have to add two new columns for that, putting rows into each of the two times two classes.

```
pref_stat_BC$quantile_low_up_1 <- as.integer(cut(pref_stat_BC$log_light_HC.z, 
    quantile(pref_stat_BC$log_light_HC.z, probs = c(0, 0.5, 1), 
        na.rm = T), include.lowest = TRUE))
pref_stat_BC$quantile_low_up_1 <- paste0("q", pref_stat_BC$quantile_low_up_1)
pref_stat_BC$quantile_low_up_2 <- as.integer(cut(pref_stat_BC$log_light_TLC.z, 
    quantile(pref_stat_BC$log_light_TLC.z, probs = c(0, 0.5, 
        1), na.rm = T), include.lowest = TRUE))
pref_stat_BC$quantile_low_up_2 <- paste0("q", pref_stat_BC$quantile_low_up_2)
```

We run the categorical model (we again use the same seed as before, but it again doesn’t matter at all really).

```
# Exclude NA measurements
mod2.2_cat <- suppressMessages(brm(court_lin_heu_01 ~ redYN * 
    quantile_low_up_1 * quantile_low_up_2 + (1 | ID) + (1 | date_trial) + 
    (1 | broods), prior = c(set_prior("normal(0,3)", class = "b", 
    coef = "redYN1"), set_prior("normal(0,3)", class = "Intercept")), 
    family = "bernoulli", data = pref_stat_BC[pref_stat_BC$quantile_low_up_1 != 
        "qNA" & pref_stat_BC$quantile_low_up_2 != "qNA", ], chains = 5, 
    iter = 6000, warmup = 3000, refresh = 0, silent = TRUE, seed = 26, 
    control = list(adapt_delta = 0.99)))
```

We return back to the main model.

From here on, we repeat again almost the exact same steps as for the pure taxa, which is why they will be commented on less extensively.

Model checking. We see “hairy caterpillars” on the right!

```
plot(mod2.2)
```

Posterior predictive checks:

```
suppressMessages(pp_check(mod2.2))
```

Model coefficients:

```
fixef(mod2.2)[, 1]
```

```
##                             Intercept                                redYN1 
##                           -0.09184538                           -0.06623930 
##                        log_light_HC.z                       log_light_TLC.z 
##                           -0.06555397                            0.07592584 
##                 redYN1:log_light_HC.z                redYN1:log_light_TLC.z 
##                           -0.13387040                            0.11339109 
##        log_light_HC.z:log_light_TLC.z redYN1:log_light_HC.z:log_light_TLC.z 
##                           -0.05097461                           -0.03651975
```

Next, we want to look at model estimators (estimated marginal means) and contrasts.

Let’s look at the credible intervals for `redYN` alone.

```
group_est_main_BC <- suppressWarnings(suppressMessages(emmeans(mod2.2, 
    ~redYN, transform = "response")))
```

We extract the posteriors.

```
group_posteriors_main_BC <- extract_emm_posterior(group_est_main_BC)
```

Split posteriors by redYN

```
non_red_posterior_main <- group_posteriors_main_BC$value[group_posteriors_main_BC$redYN == 
    "N"]
red_posterior_main <- group_posteriors_main_BC$value[group_posteriors_main_BC$redYN == 
    "Y"]
```

Create “difference posterior”

```
difference_posterior_main_BC <- red_posterior_main - non_red_posterior_main
```

Now we take a look at mean and 95% CrI for non-red backcross, red backcross and the difference between them.

```
est_redYN <- data.frame(c("N", "Y", "Y-N"), rbind(c(mean(non_red_posterior_main), 
    quantile(non_red_posterior_main, probs = c(0.025, 0.975))), 
    c(mean(red_posterior_main), quantile(red_posterior_main, 
        probs = c(0.025, 0.975))), c(mean(difference_posterior_main_BC), 
        quantile(difference_posterior_main_BC, probs = c(0.025, 
            0.975)))))
names(est_redYN) <- c("object", "mean", "lower", "upper")
print(est_redYN)
```

```
##   object       mean       lower      upper
## 1      N 0.46113367  0.41407451 0.50801677
## 2      Y 0.49345021  0.44358742 0.54251053
## 3    Y-N 0.03231653 -0.01759918 0.08253211
```

We see an effect size of about 0.03.

Now we test how likely it is that the predictors for non-red and red backcross differ. We test three hypotheses:

1. non-red males have a smaller preference score than red males.
2. red males show a preference for their own type.
3. non-red males show a preference for their own type.

Again, 2) and 3) don’t matter very much.

```
print(data.frame(test = c("Y-N > 0", "Y > 0", "N < 0"), posterior_prop = c(sum(difference_posterior_main_BC > 
    0)/length(difference_posterior_main_BC), sum(red_posterior_main > 
    0.5)/length(red_posterior_main), sum(non_red_posterior_main < 
    0.5)/length(non_red_posterior_main))))
```

```
##      test posterior_prop
## 1 Y-N > 0      0.8968667
## 2   Y > 0      0.3980667
## 3   N < 0      0.9452667
```

We can see that the posterior overlap of the two predictors is quite small! Therefore it is quite probable that the preference of the males with and without red differ. The probability for red males to have a preference is close to 50%, meaning that there is no evidence for them having a preference for their own type. We are more certain that males without red have a preference for their own type.

Now, let’s look at the **simple** slopes of the interaction terms between `redYN` and log-illuminance at the *heu* ‘female’.

We use the `emtrends` function again.

```
group_est_main_BC_interact <- suppressWarnings(suppressMessages(emtrends(mod2.2, 
    ~redYN, var = "log_light_HC.z", transform = "response")))
```

Mean and 95% equal-tailed interval.

```
print(extract_emm_cri(group_est_main_BC_interact))
```

```
##   redYN    estimate       lower       upper
## 1     N -0.04943635 -0.08210625 -0.01705253
## 2     Y  0.01703313 -0.01420864  0.04848909
```

We get the posteriors for the simple slopes

```
group_posteriors_main_BC_interact <- extract_emm_posterior(group_est_main_BC_interact)
```

We use the same hypotheses as before. In the interaction term context, they mean again something slightly different.

1. The slope of the interaction term between *lin* and log-illuminance at *heu* ‘female’ is smaller than the one of the interaction term between *heu* and log-illuminance at *heu* ‘female’.
2. Increase in log-illuminance around the *heu* ‘female’ leads to an increase in preference of *heu* males for their own type (positive slope).
3. Increase in log-illuminance around the *heu* ‘female’ leads to an increase in preference of *lin* males for their own type (negative slope).

```
iterations <- nrow(group_posteriors_main_BC_interact)/2
print(data.frame(test = c("Y > N", "Y > 0", "N < 0"), posterior_prob = c(sum(group_posteriors_main_BC_interact$value[group_posteriors_main_BC_interact$redYN == 
    "Y"] > group_posteriors_main_BC_interact$value[group_posteriors_main_BC_interact$redYN == 
    "N"])/iterations, sum(group_posteriors_main_BC_interact$value[group_posteriors_main_BC_interact$redYN == 
    "Y"] > 0)/iterations, sum(group_posteriors_main_BC_interact$value[group_posteriors_main_BC_interact$redYN == 
    "N"] < 0)/iterations)))
```

```
##    test posterior_prob
## 1 Y > N      0.9991333
## 2 Y > 0      0.8592000
## 3 N < 0      0.9986667
```

We can see that with a high probability, the slopes of the two interaction terms differ. The posteriors almost don’t overlap at all. The slope for males without red colour is with very high probability negative. The slope of males with red on the wings is positive, but the support is less strong.

We repeat the same analyses with same null hypotheses on the interaction term between male colouration and the log-illuminance of the *lin* ‘female’.

`emtrends`:

```
group_est_main_BC_interact <- suppressWarnings(suppressMessages(emtrends(mod2.2, 
    ~redYN, var = "log_light_TLC.z", transform = "response")))
```

Mean and 95% equal-tailed interval.

```
print(extract_emm_cri(group_est_main_BC_interact))
```

```
##   redYN    estimate       lower      upper
## 1     N  0.04714985  0.01643107 0.07767933
## 2     Y -0.00930522 -0.03981547 0.02188869
```

We get the posteriors for the simple slopes

```
group_posteriors_main_BC_interact <- extract_emm_posterior(group_est_main_BC_interact)
```

Hypothesis testing:

```
iterations <- nrow(group_posteriors_main_BC_interact)/2
print(data.frame(test = c("Y > N", "Y > 0", "N < 0"), posterior_prob = c(sum(group_posteriors_main_BC_interact$value[group_posteriors_main_BC_interact$redYN == 
    "Y"] > group_posteriors_main_BC_interact$value[group_posteriors_main_BC_interact$redYN == 
    "N"])/iterations, sum(group_posteriors_main_BC_interact$value[group_posteriors_main_BC_interact$redYN == 
    "Y"] > 0)/iterations, sum(group_posteriors_main_BC_interact$value[group_posteriors_main_BC_interact$redYN == 
    "N"] < 0)/iterations)))
```

```
##    test posterior_prob
## 1 Y > N      0.0030000
## 2 Y > 0      0.2775333
## 3 N < 0      0.0014000
```

We can see that there is not much evidence for intensity at the *lin* ‘female’ affecting preference of males with red wing colour (termed `Y`). Males without red on the wings though seem to get more attracted to the *heu* ‘female’ if the *lin* ‘female’ is bright. The hypothesis of a negative slope is not met and both, posterior probability and Bayes factor show how unlikely a negative slope is. This positive slope of males without red causes also that there is little posterior probability for the slope of red males being bigger (in a sense of “more positive”) than the slope of non-red males (in fact, it seems to be the other way around).

We repeat a similar procedure with the categorical illuminace model, `mod2.2_cat`.

Extract the posteriors for all levels of redYN from the categorical model, under varying categorical illumination at *heu* and at the *lin* model.

```
group_est_cat_BC <- suppressWarnings(suppressMessages(emmeans(mod2.2_cat, 
    ~redYN | quantile_low_up_1 | quantile_low_up_2, transform = "response")))
```

We extract the posteriors.

```
group_posteriors_cat <- extract_emm_posterior(group_est_cat_BC)
```

Split posteriors by redYN and illuminance categories at each models

```
non_red_posterior_cat_1_1 <- group_posteriors_cat$value[group_posteriors_cat$redYN == 
    "N" & group_posteriors_cat$quantile_low_up_1 == "q1" & group_posteriors_cat$quantile_low_up_2 == 
    "q1"]
red_posterior_cat_1_1 <- group_posteriors_cat$value[group_posteriors_cat$redYN == 
    "Y" & group_posteriors_cat$quantile_low_up_1 == "q1" & group_posteriors_cat$quantile_low_up_2 == 
    "q1"]
non_red_posterior_cat_1_2 <- group_posteriors_cat$value[group_posteriors_cat$redYN == 
    "N" & group_posteriors_cat$quantile_low_up_1 == "q1" & group_posteriors_cat$quantile_low_up_2 == 
    "q2"]
red_posterior_cat_1_2 <- group_posteriors_cat$value[group_posteriors_cat$redYN == 
    "Y" & group_posteriors_cat$quantile_low_up_1 == "q1" & group_posteriors_cat$quantile_low_up_2 == 
    "q2"]
non_red_posterior_cat_2_1 <- group_posteriors_cat$value[group_posteriors_cat$redYN == 
    "N" & group_posteriors_cat$quantile_low_up_1 == "q2" & group_posteriors_cat$quantile_low_up_2 == 
    "q1"]
red_posterior_cat_2_1 <- group_posteriors_cat$value[group_posteriors_cat$redYN == 
    "Y" & group_posteriors_cat$quantile_low_up_1 == "q2" & group_posteriors_cat$quantile_low_up_2 == 
    "q1"]
non_red_posterior_cat_2_2 <- group_posteriors_cat$value[group_posteriors_cat$redYN == 
    "N" & group_posteriors_cat$quantile_low_up_1 == "q2" & group_posteriors_cat$quantile_low_up_2 == 
    "q2"]
red_posterior_cat_2_2 <- group_posteriors_cat$value[group_posteriors_cat$redYN == 
    "Y" & group_posteriors_cat$quantile_low_up_1 == "q2" & group_posteriors_cat$quantile_low_up_2 == 
    "q2"]
```

Create “difference posteriors”

```
difference_posterior_cat_1_1 <- red_posterior_cat_1_1 - non_red_posterior_cat_1_1
difference_posterior_cat_1_2 <- red_posterior_cat_1_2 - non_red_posterior_cat_1_2
difference_posterior_cat_2_1 <- red_posterior_cat_2_1 - non_red_posterior_cat_2_1
difference_posterior_cat_2_2 <- red_posterior_cat_2_2 - non_red_posterior_cat_2_2
```

Now we take a look at mean and 95% CrI for non-red backcrosses, red backcrosses and the difference between them. To not have too many lines, we skip pats of the previously used code by using `extract_emm_cri` instead.

```
est_types <- extract_emm_cri(group_est_cat_BC)
names(est_types) <- c("object", "light_env_heu", "light_env_lin", 
    "mean", "lower", "upper")
contrast_table <- unname(data.frame(rep("Y-N", 4), est_types[c(1, 
    3, 5, 7), 2:3], rbind(c(mean(difference_posterior_cat_1_1), 
    quantile(difference_posterior_cat_1_1, probs = c(0.025, 0.975))), 
    c(mean(difference_posterior_cat_2_1), quantile(difference_posterior_cat_2_1, 
        probs = c(0.025, 0.975))), c(mean(difference_posterior_cat_1_2), 
        quantile(difference_posterior_cat_1_2, probs = c(0.025, 
            0.975))), c(mean(difference_posterior_cat_2_2), quantile(difference_posterior_cat_2_2, 
        probs = c(0.025, 0.975)))), stringsAsFactors = F))
names(contrast_table) <- c("object", "light_env_heu", "light_env_lin", 
    "mean", "lower", "upper")
est_types <- rbind(est_types[1:2, ], contrast_table[1, ], est_types[3:4, 
    ], contrast_table[2, ], est_types[5:6, ], contrast_table[3, 
    ], est_types[7:8, ], contrast_table[4, ])
rownames(est_types) <- NULL
print(est_types)
```

```
##    object light_env_heu light_env_lin        mean        lower      upper
## 1       N            q1            q1  0.46025930  0.406151422 0.51529526
## 2       Y            q1            q1  0.47497389  0.418786223 0.53088793
## 3     Y-N            q1            q1  0.01471459 -0.048053362 0.07721418
## 4       N            q2            q1  0.42570502  0.349654163 0.50482659
## 5       Y            q2            q1  0.53845261  0.461047909 0.61563216
## 6     Y-N            q2            q1  0.11274760  0.015179441 0.20718870
## 7       N            q1            q2  0.47748212  0.401311325 0.55407570
## 8       Y            q1            q2  0.44344391  0.359137639 0.52805033
## 9     Y-N            q1            q2 -0.03403821 -0.134976273 0.06642484
## 10      N            q2            q2  0.43810841  0.383992263 0.49133469
## 11      Y            q2            q2  0.50425459  0.444568596 0.56280278
## 12    Y-N            q2            q2  0.06614618  0.002615173 0.12769096
```

Looks pretty similar to the continuous model! Only the effect of illuminance at the *lin* model seems to be not as strong.

Some of these calculations will be repeated by the below code to produce the plots.

##### 2.2.2 Figure 2

Get `emmeans` estimates for the interaction between log-illuminance around *heu* ‘female’ and type (species) from `mod1.208`. We sample at 100 values distributes across the range of illuminance values.

```
# We find the range of illuminance intensities at heu mounted
# female
data_range <- range(mod1.208$data$log_light_HC.z)
# We produce 100 values from the range between the two
# extremes
val_to_check <- do.call(seq, as.list(c(data_range, dist(data_range)/99)))
# Run emmeans
pure_interaction <- extract_emm_cri(emmeans(mod1.208, ~type | 
    log_light_HC.z, transform = "response", at = list(log_light_HC.z = val_to_check)))
```

We backtransform every value of `log_light_HC.z`, using the scaling parameters safed in the original table.

```
pure_interaction$log_light_HC <- pure_interaction$log_light_HC.z * 
    attr(pref_stat_TLC_HC$log_light_HC.z, "scaled:scale") + attr(pref_stat_TLC_HC$log_light_HC.z, 
    "scaled:center")
```

Create histogram of the difference posteriors in the Gardner-Altman plots. We take a BIN size of 0.005.

```
post_hist_main <- hist(difference_posterior_main, breaks = seq(floor(200 * 
    min(difference_posterior_main))/200, ceiling(200 * max(difference_posterior_main))/200, 
    0.005), plot = F)
post_hist_cat_1 <- hist(difference_posterior_cat_1, breaks = seq(floor(200 * 
    min(difference_posterior_cat_1))/200, ceiling(200 * max(difference_posterior_cat_1))/200, 
    0.005), plot = F)
post_hist_cat_2 <- hist(difference_posterior_cat_2, breaks = seq(floor(200 * 
    min(difference_posterior_cat_2))/200, ceiling(200 * max(difference_posterior_cat_2))/200, 
    0.005), plot = F)
```

The workflow:

- Call `other_plot_functions.R` to plot panel A
- Calling `plot_proportion_stats` to produce three nicely jittered stripcharts (panel B-D)

```
pdf("Graphs/figure2.pdf",width=9,height=6)
layout(matrix(1:4,ncol=2,byrow=T))

par(oma=c(0,0.8,0,4.2))

#Produce panel A by just calling the following function (if you want to check aesthetic details, check the function in other_plot_functions.R). We present illuminance values as on a log scale.
plot_interaction(covariate="log_light_HC",bay_cov1=pure_interaction,
                 text_inner_fill="",added_title=substitute(paste("Illuminance at mounted ",italic('H. heurippa')," female [",italic('lux'),"]")),
                 xvalues=c(2,5,15,50,150),par_mar1 = c(3,2.9,2,0.3))

#Add line showing cutoff for dark vs bright
abline(v=median(pref_stat_TLC_HC$log_light_HC,na.rm = T),lty="dashed")

#Calling plot_proportion_stats to produce the panel with the full data
testout_bayes<-plot_proportion_stats(input_data=pref_stat_TLC_HC,
                                     # The data set
                                     sum_by_col_plot=c("ID"),
                                     # Sum by ID for plot
                                     response_col="court_lin_heu_01",
                                     # Response variable
                                     sub1_col="type",
                                     # Fixed effect in model on heu and lin
sub1_states=c("TLC","HC"),
                                     # Panels from left right
                                     stats_for_1sub=F,
                                     #Aesthetics:
                                     par_mar=c(2.4,2.9,2,5),
                                     # Plot margins
                                     sub1_labels=c("H. t. linaresi","H. heurippa"),
                                     # x axis labels
                                     yaxs_label="",
                                     # y axis title
                                     y_tck=-0.015,
                                     # Tick size on y-axis
                                     horiz_0.5=F,
                                     # No horizontal line at 0.5
                                     panel_dividers=F,
                                     # No vertical panel dividers
                                     dot_size=0.26,
                                     # Factor to multiply dot sizes with (just aesthetics)
                                     dot_colours=list("blue","red"),
                                     # Dot colours
                                     sub_box_yn=F,
                                     # No Boxplot added
                                     numb_iterations=1000,
                                     # Number iterations for assembling the points
                                     legend_dots=rev(c(10,20,50,100,200)),
                                     # Legend dots
                                     legend_omi=T,
                                     # Use outer margin for legend
                                     squeeze_legend=1.3,
                                     # Tighten space vertically in legend
                                     legend_info=F,
                                     # Add no additional legend info
                                     border_space=0.49,
                                     # space to left and right side of center that dots can be jittered to
                                                                          seedi=123,
                                     # Seed
                                     verbose=F
                                     # Do not print comments
)

#We add a letter to the figure for this panel
text(par("usr")[1]+(0.2/par("pin")[1])*diff(par("usr")[1:2]),
     par("usr")[4]-(0.15/par("pin")[2])*diff(par("usr")[3:4]),
     "B",font=2,cex=1.5,
     pos=2,offset=0)

#Scale all histograms so that length of the maximum bar of all posteriors is maximally 7/9 of the distance the axis has to the plot

max_bar<-max(c(post_hist_main$counts,post_hist_cat_1$counts,post_hist_cat_2$counts))
post_hist_main$counts<-(post_hist_main$counts/max_bar)*(par("cex")*1.4*par("cin")[2]/par("pin")[1])*diff(par("usr")[1:2])
post_hist_cat_1$counts<-(post_hist_cat_1$counts/max_bar)*(par("cex")*1.4*par("cin")[2]/par("pin")[1])*diff(par("usr")[1:2])
post_hist_cat_2$counts<-(post_hist_cat_2$counts/max_bar)*(par("cex")*1.4*par("cin")[2]/par("pin")[1])*diff(par("usr")[1:2])

#We add the Gardner-Altman part
add_gardner_altman(TLC_posterior_main,HC_posterior_main,difference_posterior_main,post_hist_main)

#We add the same type of swarm plot, but this time just including data from dark conditions
testout_bayes<-plot_proportion_stats(input_data=pref_stat_TLC_HC[pref_stat_TLC_HC$quantile_low_up=="q1",],
                                     # The data set
                                     sum_by_col_plot=c("ID"),
                                     # Sum by ID for plot
                                     response_col="court_lin_heu_01",
                                     # Response variable
                                     sub1_col="type",
                                     # Fixed effect in model on heu and lin
sub1_states=c("TLC","HC"),
                                     # Panels from left right
                                     stats_for_1sub=F,
                                     #Aesthetics:
                                     par_mar=c(2.4,2.9,2,5),
                                     # Plot margins
                                     sub1_labels=c("H. t. linaresi","H. heurippa"),
                                     # x axis labels
                                     yaxs_label="",
                                     # y axis title
                                     y_tck=-0.015,
                                     # Tick size on y-axis
                                     horiz_0.5=F,
                                     # No horizontal line at 0.5
                                     panel_dividers=F,
                                     # No vertical panel dividers
                                     dot_size=0.26,
                                     # Factor to multiply dot sizes with (just aesthetics)
                                     dot_colours=list("blue","red"),
                                     # Dot colours
                                     sub_box_yn=F,
                                     # No Boxplot added
                                     numb_iterations=1000,
                                     # Number iterations for assembling the points
                                     legend_dots=c(),
                                     # Add no legend
                                     legend_omi=T,
                                     # Use outer margin for legend
                                     squeeze_legend=1.3,
                                     # Tighten space vertically in legend
                                     legend_info=F,
                                     # Add no additional legend info
                                     border_space=0.49,
                                     # space to left and right side of center that dots can be jittered to
                                                                          seedi=123,
                                     # Seed
                                     verbose=F
                                     # Do not print comments
)

#We add a letter to the figure for this panel
text(par("usr")[1]+(0.2/par("pin")[1])*diff(par("usr")[1:2]),
     par("usr")[4]-(0.15/par("pin")[2])*diff(par("usr")[3:4]),
     "C",font=2,cex=1.5,
     pos=2,offset=0)

#We add the Gardner-Altman part for that panel
add_gardner_altman(TLC_posterior_cat_1,HC_posterior_cat_1,difference_posterior_cat_1,post_hist_cat_1)


#We add the same type of swarm plot, but this time just including data from bright conditions
testout_bayes<-plot_proportion_stats(input_data=pref_stat_TLC_HC[pref_stat_TLC_HC$quantile_low_up=="q2",],
                                     # The data set
                                     sum_by_col_plot=c("ID"),
                                     # Sum by ID for plot
                                     response_col="court_lin_heu_01",
                                     # Response variable
                                     sub1_col="type",
                                     # Fixed effect in model on heu and lin
sub1_states=c("TLC","HC"),
                                     # Panels from left right
                                     stats_for_1sub=F,
                                     #Aesthetics:
                                     par_mar=c(2.4,2.9,2,5),
                                     # Plot margins
                                     sub1_labels=c("H. t. linaresi","H. heurippa"),
                                     # x axis labels
                                     yaxs_label="",
                                     # y axis title
                                     y_tck=-0.015,
                                     # Tick size on y-axis
                                     horiz_0.5=F,
                                     # No horizontal line at 0.5
                                     panel_dividers=F,
                                     # No vertical panel dividers
                                     dot_size=0.26,
                                     # Factor to multiply dot sizes with (just aesthetics)
                                     dot_colours=list("blue","red"),
                                     # Dot colours
                                     sub_box_yn=F,
                                     # No Boxplot added
                                     numb_iterations=1000,
                                     # Number iterations for assembling the points
                                     legend_dots=rev(c(5,10,20,50,100)),
                                     # Legend dots
                                     legend_omi=T,
                                     # Use outer margin for legend
                                     squeeze_legend=1.3,
                                     # Tighten space vertically in legend
                                     legend_info=F,
                                     # Add no additional legend info
                                     border_space=0.49,
                                     # space to left and right side of center that dots can be jittered to
                                                                          seedi=123,
                                     # Seed
                                     verbose=F
                                     # Do not print comments
)

#We add a letter to the figure for this panel
text(par("usr")[1]+(0.2/par("pin")[1])*diff(par("usr")[1:2]),
     par("usr")[4]-(0.15/par("pin")[2])*diff(par("usr")[3:4]),
     "D",font=2,cex=1.5,
     pos=2,offset=0)

#We add the Gardner-Altman part for that panel
add_gardner_altman(TLC_posterior_cat_2,HC_posterior_cat_2,difference_posterior_cat_2,post_hist_cat_2)

mtext(substitute(paste("Proportion interactions with mounted  ",italic('H. heurippa')," female")),
      2,outer=T,line=-0.4,cex=0.8)

#Close plotting device.
invisible(dev.off())
```

We convert the pdf to png, so we have both file types at hand later. Unfortunately, the Greek Delta sign gets lost in this process… It is there though in the pdf.

```
pdf_convert("Graphs/figure2.pdf", filenames = "Graphs/figure2.png", 
    dpi = 300)
```

```
## Converting page 1 to Graphs/figure2.png... done!
```

```
## [1] "Graphs/figure2.png"
```

Show png we just made

```
fig2 <- readPNG("Graphs/figure2.png")
grid.raster(fig2)
```

##### 2.2.3 Figure 4

Now we look at how the light environment influences visual preferences in differently coloured backcross to *lin* males.

Create histogram of the difference posteriors in the following Gardner-Altman plots. We take a BIN size of 0.005.

```
post_hist_BC_cat_1_1 <- hist(difference_posterior_cat_1_1, breaks = seq(floor(200 * 
    min(difference_posterior_cat_1_1))/200, ceiling(200 * max(difference_posterior_cat_1_1))/200, 
    0.005), plot = F)
post_hist_BC_cat_1_2 <- hist(difference_posterior_cat_1_2, breaks = seq(floor(200 * 
    min(difference_posterior_cat_1_2))/200, ceiling(200 * max(difference_posterior_cat_1_2))/200, 
    0.005), plot = F)
post_hist_BC_cat_2_1 <- hist(difference_posterior_cat_2_1, breaks = seq(floor(200 * 
    min(difference_posterior_cat_2_1))/200, ceiling(200 * max(difference_posterior_cat_2_1))/200, 
    0.005), plot = F)
post_hist_BC_cat_2_2 <- hist(difference_posterior_cat_2_2, breaks = seq(floor(200 * 
    min(difference_posterior_cat_2_2))/200, ceiling(200 * max(difference_posterior_cat_2_2))/200, 
    0.005), plot = F)
```

We produce the figure.

```
pdf("Graphs/figure4.pdf",height=7.5,width=5)

layout(matrix(c(rep(1,6),rep(2,5),rep(3,6),rep(4,5)),nrow=2,byrow=T))

#We reduce the cex parameter so that elements don't get too big
par(cex=0.66)

par(oma=c(0,1.6,0,0))

#We add the same type of swarm plot, but this time just including data from dark conditions
testout_bayes<-plot_proportion_stats(input_data=pref_stat_BC[pref_stat_BC$quantile_low_up_1=="q1"&pref_stat_BC$quantile_low_up_2=="q1",],
                                     # The data set
                                     sum_by_col_plot=c("ID"),
                                     # Sum by ID for plot
                                     response_col="court_lin_heu_01",
                                     # Response variable
                                     sub1_col="redYN",
                                     # First fixed effect in model on heu and lin
                                     sub1_states=c("N","Y"),
                                     # Panels from left right
                                     stats_for_1sub=F,
                                     #Aesthetics:
                                     par_mar=c(2.6,2.4,2,8.298),
                                     # Plot margins
                                     sub1_labels=c("Non-Red","Red"),
                                     # x axis labels
                                     yaxs_label="",
                                     # y axis title
                                     y_tck=-0.015,
                                     # Tick size on y-axis
                                     horiz_0.5=F,
                                     # No horizontal line at 0.5
                                     panel_dividers=F,
                                     # No vertical panel dividers
                                     dot_size=0.26,
                                     # Factor to multiply dot sizes with (just aesthetics)
                                     dot_colours=list("blue","red"),
                                     # Dot colours
                                     sub_box_yn=F,
                                     # No Boxplot added
                                     numb_iterations=1000,
                                     # Number iterations for assembling the points
                                     legend_dots=rev(c(5,10,20,50,100)),
                                     # Legend dots
                                     squeeze_legend=0.75,
                                     # Tighten space vertically in legend
                                     legend_info=F,
                                     # Add no additional legend info
                                     legend_omi=T,
                                     # Use outer plot space
                                     border_space=0.49,
                                     # space to left and right side of center that dots can be jittered to
                                     seedi=123,
                                     # Seed
                                     verbose=F
                                     # Do not print comments
)

#We add a letter to the figure for this panel
text(par("usr")[1]+(0.2/par("pin")[1])*diff(par("usr")[1:2]),
     par("usr")[4]-(0.25/par("pin")[2])*diff(par("usr")[3:4]),
     "A",font=2,cex=1.5,
     pos=2,offset=0)

#Scale all histograms so that length of the maximum bar of all posteriors is maximally 7/9 of the distance the axis has to the plot
max_bar<-max(c(post_hist_BC_cat_1_1$counts,post_hist_BC_cat_1_2$counts,post_hist_BC_cat_2_1$counts,post_hist_BC_cat_2_2$counts))
post_hist_BC_cat_1_1$counts<-(post_hist_BC_cat_1_1$counts/max_bar)*(par("cex")*1.4*par("cin")[2]/par("pin")[1])*diff(par("usr")[1:2])
post_hist_BC_cat_1_2$counts<-(post_hist_BC_cat_1_2$counts/max_bar)*(par("cex")*1.4*par("cin")[2]/par("pin")[1])*diff(par("usr")[1:2])
post_hist_BC_cat_2_1$counts<-(post_hist_BC_cat_2_1$counts/max_bar)*(par("cex")*1.4*par("cin")[2]/par("pin")[1])*diff(par("usr")[1:2])
post_hist_BC_cat_2_2$counts<-(post_hist_BC_cat_2_2$counts/max_bar)*(par("cex")*1.4*par("cin")[2]/par("pin")[1])*diff(par("usr")[1:2])

#We add the Gardner-Altman part for that panel
add_gardner_altman(non_red_posterior_cat_1_1,red_posterior_cat_1_1,difference_posterior_cat_1_1,post_hist_BC_cat_1_1,add_title=F)


#We add the same type of swarm plot, but this time just including data from dark heu and bright lin conditions
testout_bayes<-plot_proportion_stats(input_data=pref_stat_BC[pref_stat_BC$quantile_low_up_1=="q1"&pref_stat_BC$quantile_low_up_2=="q2",],
                                     # The data set
                                     sum_by_col_plot=c("ID"),
                                     # Sum by ID for plot
                                     response_col="court_lin_heu_01",
                                     # Response variable
                                     sub1_col="redYN",
                                     # First fixed effect in model on heu and lin
                                     sub1_states=c("N","Y"),
                                     # Panels from left right
                                     stats_for_1sub=F,
                                     #Aesthetics:
                                     par_mar=c(2.6,2.4,2,5),
                                     # Plot margins
                                     sub1_labels=c("Non-Red","Red"),
                                     # x axis labels
                                     yaxs_label="",
                                     # y axis title
                                     y_tck=-0.015,
                                     # Tick size on y-axis
                                     horiz_0.5=F,
                                     # No horizontal line at 0.5
                                     panel_dividers=F,
                                     # No vertical panel dividers
                                     dot_size=0.26,
                                     # Factor to multiply dot sizes with (just aesthetics)
                                     dot_colours=list("blue","red"),
                                     # Dot colours
                                     sub_box_yn=F,
                                     # No Boxplot added
                                     numb_iterations=1000,
                                     # Number iterations for assembling the points
                                     legend_dots=c(),
                                     # Add no additional legend info
                                     border_space=0.49,
                                     # space to left and right side of center that dots can be jittered to
                                     seedi=123,
                                     # Seed
                                     verbose=F
                                     # Do not print comments
)

#We add a letter to the figure for this panel
text(par("usr")[1]+(0.2/par("pin")[1])*diff(par("usr")[1:2]),
     par("usr")[4]-(0.25/par("pin")[2])*diff(par("usr")[3:4]),
     "B",font=2,cex=1.5,
     pos=2,offset=0)

#We add the Gardner-Altman part for that panel
add_gardner_altman(non_red_posterior_cat_1_2,red_posterior_cat_1_2,difference_posterior_cat_1_2,post_hist_BC_cat_1_2)


#We add the same type of swarm plot, but this time just including data lin and bright heu conditions
testout_bayes<-plot_proportion_stats(input_data=pref_stat_BC[pref_stat_BC$quantile_low_up_1=="q2"&pref_stat_BC$quantile_low_up_2=="q1",],
                                     # The data set
                                     sum_by_col_plot=c("ID"),
                                     # Sum by ID for plot
                                     response_col="court_lin_heu_01",
                                     # Response variable
                                     sub1_col="redYN",
                                     # First fixed effect in model on heu and lin
                                     sub1_states=c("N","Y"),
                                     # Panels from left right
                                     stats_for_1sub=F,
                                     #Aesthetics:
                                     par_mar=c(2.6,2.4,2,8.298),
                                     # Plot margins
                                     sub1_labels=c("Non-Red","Red"),
                                     # x axis labels
                                     yaxs_label="",
                                     # y axis title
                                     y_tck=-0.015,
                                     # Tick size on y-axis
                                     horiz_0.5=F,
                                     # No horizontal line at 0.5
                                     panel_dividers=F,
                                     # No vertical panel dividers
                                     dot_size=0.26,
                                     # Factor to multiply dot sizes with (just aesthetics)
                                     dot_colours=list("blue","red"),
                                     # Dot colours
                                     sub_box_yn=F,
                                     # No Boxplot added
                                     numb_iterations=1000,
                                     # Number iterations for assembling the points
                                     legend_dots=rev(c(5,10,20,50,100)),
                                     # Legend dots
                                     squeeze_legend=0.75,
                                     # Tighten space vertically in legend
                                     legend_info=F,
                                     # Add no additional legend info
                                     legend_omi=T,
                                     # Use outer plot space
                                     border_space=0.49,
                                     # space to left and right side of center that dots can be jittered to
                                     seedi=123,
                                     # Seed
                                     verbose=F
                                     # Do not print comments
)

#We add a letter to the figure for this panel
text(par("usr")[1]+(0.2/par("pin")[1])*diff(par("usr")[1:2]),
     par("usr")[4]-(0.25/par("pin")[2])*diff(par("usr")[3:4]),
     "C",font=2,cex=1.5,
     pos=2,offset=0)

#We add the Gardner-Altman part for that panel
add_gardner_altman(non_red_posterior_cat_2_1,red_posterior_cat_2_1,difference_posterior_cat_2_1,post_hist_BC_cat_2_1,add_title=F)


#We add the same type of swarm plot, but this time just including data from dark heu and bright lin conditions
testout_bayes<-plot_proportion_stats(input_data=pref_stat_BC[pref_stat_BC$quantile_low_up_1=="q2"&pref_stat_BC$quantile_low_up_2=="q2",],
                                     # The data set
                                     sum_by_col_plot=c("ID"),
                                     # Sum by ID for plot
                                     response_col="court_lin_heu_01",
                                     # Response variable
                                     sub1_col="redYN",
                                     # First fixed effect in model on heu and lin
                                     sub1_states=c("N","Y"),
                                     # Panels from left right
                                     stats_for_1sub=F,
                                     #Aesthetics:
                                     par_mar=c(2.6,2.4,2,5),
                                     # Plot margins
                                     sub1_labels=c("Non-Red","Red"),
                                     # x axis labels
                                     yaxs_label="",
                                     # y axis title
                                     y_tck=-0.015,
                                     # Tick size on y-axis
                                     horiz_0.5=F,
                                     # No horizontal line at 0.5
                                     panel_dividers=F,
                                     # No vertical panel dividers
                                     dot_size=0.26,
                                     # Factor to multiply dot sizes with (just aesthetics)
                                     dot_colours=list("blue","red"),
                                     # Dot colours
                                     sub_box_yn=F,
                                     # No Boxplot added
                                     numb_iterations=1000,
                                     # Number iterations for assembling the points
                                     legend_dots=c(),
                                     # Add no additional legend info
                                     border_space=0.49,
                                     # space to left and right side of center that dots can be jittered to
                                     seedi=123,
                                     # Seed
                                     verbose=F
                                     # Do not print comments
)

#We add a letter to the figure for this panel
text(par("usr")[1]+(0.2/par("pin")[1])*diff(par("usr")[1:2]),
     par("usr")[4]-(0.25/par("pin")[2])*diff(par("usr")[3:4]),
     "D",font=2,cex=1.5,
     pos=2,offset=0)

#We add the Gardner-Altman part for that panel
add_gardner_altman(non_red_posterior_cat_2_2,red_posterior_cat_2_2,difference_posterior_cat_2_2,post_hist_BC_cat_2_2)


mtext(substitute(paste("Proportion interactions with mounted  ",italic('H. heurippa')," female")),
      2,outer=T,line=0.3,cex=0.8)

#Close plotting device.
invisible(dev.off())
```

We convert the pdf to png, so we have both file types at hand later. Unfortunately, the Greek Delta sign gets lost in this process… It is there though in the pdf.

```
pdf_convert("Graphs/figure4.pdf", filenames = "Graphs/figure4.png", 
    dpi = 300)
```

```
## Converting page 1 to Graphs/figure4.png... done!
```

```
## [1] "Graphs/figure4.png"
```

Show png we just made

```
fig4 <- readPNG("Graphs/figure4.png")
grid.raster(fig4)
```

##### 2.2.4 Supplementary figure 5

Now we look at backcross to *lin* hybrids without subsetting into different light categories. We look at the model estimates from `mod2.2`.

Create histogram of the difference posterior (same as for the figure above).

```
post_hist_main_BC <- hist(difference_posterior_main_BC, breaks = seq(floor(200 * 
    min(difference_posterior_main_BC))/200, ceiling(200 * max(difference_posterior_main_BC))/200, 
    0.005), plot = F)
```

Produce the figure.

```
png("Graphs/supplementary_figure5.png",width=2000,height=1600,res=300)
#Calling plot_proportion_stats to produce the panel with the full data
testout_bayes<-plot_proportion_stats(input_data=pref_stat_BC,
                                     # The data set
                                     sum_by_col_plot=c("ID"),
                                     # Sum by ID for plot
                                     response_col="court_lin_heu_01",
                                     # Response variable
                                     sub1_col="redYN",
                                     # First fixed effect in model on heu and lin
                                     sub1_states=c("N","Y"),
                                     # Panels from left right
                                     stats_for_1sub=F,
                                     #Aesthetics:
                                     par_mar=c(2.6,3,0.3,9),
                                     # Plot margins
                                     sub1_labels=c("Non-Red","Red"),
                                     # x axis labels
                                     yaxs_label=substitute(paste("Proportion interactions with mounted  ",italic('H. heurippa')," female")),
                                     # y axis title
                                     y_tck=-0.015,
                                     # Tick size on y-axis
                                     horiz_0.5=F,
                                     # No horizontal line at 0.5
                                     panel_dividers=F,
                                     # No vertical panel dividers
                                     dot_size=0.26,
                                     # Factor to multiply dot sizes with (just aesthetics)
                                     dot_colours=list("blue","red"),
                                     # Dot colours
                                     sub_box_yn=F,
                                     # No Boxplot added
                                     numb_iterations=1000,
                                     # Number iterations for assembling the points
                                     legend_dots=rev(c(10,20,50,100,200)),
                                     # Legend dots
                                     legend_omi=T,
                                     # Use outer margin for legend
                                     squeeze_legend=0.75,
                                     # Tighten space vertically in legend
                                     legend_info=F,
                                     # Add no additional legend info
                                     border_space=0.49,
                                     # space to left and right side of center that dots can be jittered to
                                     seedi=123,
                                     # Seed
                                     verbose=F
                                     # Do not print comments
)

#Scale histogram
post_hist_main_BC$counts<-(post_hist_main_BC$counts/max(post_hist_main_BC$counts))*(par("cex")*1.4*par("cin")[2]/par("pin")[1])*diff(par("usr")[1:2])

#We add the Gardner-Altman part
add_gardner_altman(non_red_posterior_main,red_posterior_main,difference_posterior_main_BC,post_hist_main_BC)

#We add the estimators from figure 2B onto the right side
par(xpd=NA)
segments(2+(par("cex")*1.3*par("cin")[2]/par("pin")[1])*diff(par("usr")[1:2]),mean(TLC_posterior_main),2+(par("cex")*1.7*par("cin")[2]/par("pin")[1])*diff(par("usr")[1:2]),mean(TLC_posterior_main),lwd=3,col="blue",lend=3)
segments(2+(par("cex")*1.3*par("cin")[2]/par("pin")[1])*diff(par("usr")[1:2]),mean(HC_posterior_main),2+(par("cex")*1.7*par("cin")[2]/par("pin")[1])*diff(par("usr")[1:2]),mean(HC_posterior_main),lwd=3,col="red",lend=3)
par(xpd=F)

#Close plotting device.
invisible(dev.off())
```

Show png we just made

```
suppl_fig5 <- readPNG("Graphs/supplementary_figure5.png")
grid.raster(suppl_fig5)
```

##### 2.2.5 Supplementary figure 4

We now want to produce an overview over different hybrid crosses, and how they are affected by illuminance at the *heu* model. Also, we want to show how preferences of individuals change, therefore we’ll produce so-called “reaction norm” plots.

In order to do this, we have to restructure the posteriors we created for *lin*, *heu* and the backcross to *lin* hybrids. Additionally, we have to let a new categorical model run for F1 and backcross to *heu* hybrids.

First, create a new dataset for F1 and backcross to *heu* hybrids.

```
pref_stat_other_hyb <- pref_stat[pref_stat$type %in% c("HCxTLC", 
    "TLCxHC", "HCx(TLCxHC)"), ]
# We unite both types of F1 crosses
pref_stat_other_hyb$type[pref_stat_other_hyb$type %in% c("HCxTLC", 
    "TLCxHC")] <- "F1"
# center+scale log-illuminance
pref_stat_other_hyb$log_light_HC.z <- scale(pref_stat_other_hyb$log_light_HC, 
    center = T, scale = T)
# Make relevant columns factors
make_fact <- c("type", "ID", "date_trial", "broods")
for (change_fact in 1:length(make_fact)) {
    pref_stat_other_hyb[, make_fact[change_fact]] <- as.factor(pref_stat_other_hyb[, 
        make_fact[change_fact]])
}
```

We create two categories for light conditions.

```
pref_stat_other_hyb$quantile_low_up <- as.integer(cut(pref_stat_other_hyb$log_light_HC.z, 
    quantile(pref_stat_other_hyb$log_light_HC.z, probs = c(0, 
        0.5, 1), na.rm = T), include.lowest = TRUE))
pref_stat_other_hyb$quantile_low_up <- paste0("q", pref_stat_other_hyb$quantile_low_up)
```

We run a new model for these two types.

```
mod_other_hyb_cat <- suppressMessages(brm(court_lin_heu_01 ~ 
    type * quantile_low_up + (1 | ID) + (1 | date_trial) + (1 | 
        broods), prior = c(set_prior("normal(0,3)", class = "b", 
    coef = "type1"), set_prior("normal(0,3)", class = "Intercept")), 
    family = "bernoulli", data = pref_stat_other_hyb[pref_stat_other_hyb$quantile_low_up != 
        "qNA", ], chains = 5, iter = 6000, warmup = 3000, refresh = 0, 
    silent = TRUE, seed = 45, control = list(adapt_delta = 0.999)))
```

Extract the posteriors for all levels of type from the categorical model, under varying categorical illumination at *heu* model

```
other_hyb_est_cat <- suppressWarnings(suppressMessages(emmeans(mod_other_hyb_cat, 
    ~type | quantile_low_up, transform = "response")))
```

We extract the posteriors.

```
other_hyb_posteriors_cat <- extract_emm_posterior(other_hyb_est_cat)
```

Split posteriors by type and illuminance

```
F1_posterior_cat_1 <- other_hyb_posteriors_cat$value[other_hyb_posteriors_cat$type == 
    "F1" & other_hyb_posteriors_cat$quantile_low_up == "q1"]
BC_H_posterior_cat_1 <- other_hyb_posteriors_cat$value[other_hyb_posteriors_cat$type == 
    "HCx(TLCxHC)" & other_hyb_posteriors_cat$quantile_low_up == 
    "q1"]
F1_posterior_cat_2 <- other_hyb_posteriors_cat$value[other_hyb_posteriors_cat$type == 
    "F1" & other_hyb_posteriors_cat$quantile_low_up == "q2"]
BC_H_posterior_cat_2 <- other_hyb_posteriors_cat$value[other_hyb_posteriors_cat$type == 
    "HCx(TLCxHC)" & other_hyb_posteriors_cat$quantile_low_up == 
    "q2"]
```

Create “difference posteriors” (this time, we compare **within** type, **not between** types)

```
difference_posterior_F1 <- F1_posterior_cat_2 - F1_posterior_cat_1
difference_posterior_BC_H <- BC_H_posterior_cat_2 - BC_H_posterior_cat_1
```

Get the histograms of the differences

```
post_hist_F1 <- hist(difference_posterior_F1, breaks = seq(floor(200 * 
    min(difference_posterior_F1))/200, ceiling(200 * max(difference_posterior_F1))/200, 
    0.005), plot = F)
post_hist_BC_H <- hist(difference_posterior_BC_H, breaks = seq(floor(200 * 
    min(difference_posterior_BC_H))/200, ceiling(200 * max(difference_posterior_BC_H))/200, 
    0.005), plot = F)
```

Now we update the backcrosses to *lin*:

Extract the posteriors for all levels of type from the categorical model, under varying categorical illumination at *heu* model

```
group_est_cat_BC_only_heu <- suppressWarnings(suppressMessages(emmeans(mod2.2_cat, 
    ~redYN | quantile_low_up_1, transform = "response")))
```

We extract the posteriors.

```
group_posteriors_cat_only_heu <- extract_emm_posterior(group_est_cat_BC_only_heu)
```

Split posteriors by colour pattern and illuminance

```
non_red_posterior_cat_1_only_heu <- group_posteriors_cat_only_heu$value[group_posteriors_cat_only_heu$redYN == 
    "N" & group_posteriors_cat_only_heu$quantile_low_up_1 == 
    "q1"]
non_red_posterior_cat_2_only_heu <- group_posteriors_cat_only_heu$value[group_posteriors_cat_only_heu$redYN == 
    "N" & group_posteriors_cat_only_heu$quantile_low_up_1 == 
    "q2"]
red_posterior_cat_1_only_heu <- group_posteriors_cat_only_heu$value[group_posteriors_cat_only_heu$redYN == 
    "Y" & group_posteriors_cat_only_heu$quantile_low_up_1 == 
    "q1"]
red_posterior_cat_2_only_heu <- group_posteriors_cat_only_heu$value[group_posteriors_cat_only_heu$redYN == 
    "Y" & group_posteriors_cat_only_heu$quantile_low_up_1 == 
    "q2"]
```

Create “difference posteriors” (this time, we compare **within** type, **not between** types)

```
non_red_difference_posterior_only_heu <- non_red_posterior_cat_2_only_heu - 
    non_red_posterior_cat_1_only_heu
red_difference_posterior_only_heu <- red_posterior_cat_2_only_heu - 
    red_posterior_cat_1_only_heu
```

Get the histograms of the differences

```
post_hist_non_red_only_heu <- hist(non_red_difference_posterior_only_heu, 
    breaks = seq(floor(200 * min(non_red_difference_posterior_only_heu))/200, 
        ceiling(200 * max(non_red_difference_posterior_only_heu))/200, 
        0.005), plot = F)
post_hist_red_only_heu <- hist(red_difference_posterior_only_heu, 
    breaks = seq(floor(200 * min(red_difference_posterior_only_heu))/200, 
        ceiling(200 * max(red_difference_posterior_only_heu))/200, 
        0.005), plot = F)
```

Now we have to update histograms for *lin* and *heu*, such that they are comparing within type

```
difference_posterior_within_TLC <- TLC_posterior_cat_2 - TLC_posterior_cat_1
difference_posterior_within_HC <- HC_posterior_cat_2 - HC_posterior_cat_1
```

New posteriors for differences

```
post_hist_within_TLC <- hist(difference_posterior_within_TLC, 
    breaks = seq(floor(200 * min(difference_posterior_within_TLC))/200, 
        ceiling(200 * max(difference_posterior_within_TLC))/200, 
        0.005), plot = F)
post_hist_within_HC <- hist(difference_posterior_within_HC, breaks = seq(floor(200 * 
    min(difference_posterior_within_HC))/200, ceiling(200 * max(difference_posterior_within_HC))/200, 
    0.005), plot = F)
```

Produce the figure.

```
png("Graphs/supplementary_figure4.png",width=6500,height=1600,res=300)

layout(matrix(1:6,ncol=6))

par(oma=c(0,1,0,5))

#lin

#Calling plot_proportion_stats to produce the panel with the full data
testout_bayes<-plot_proportion_stats(input_data=pref_stat_TLC_HC[pref_stat_TLC_HC$quantile_low_up!="qNA"&pref_stat_TLC_HC$type=="TLC",],
                                     # The data set
                                     sum_by_col_plot=c("ID","quantile_low_up"),
                                     # Sum by ID for plot
                                     response_col="court_lin_heu_01",
                                     # Response variable
                                     sub2_col="quantile_low_up",
                                     # First fixed effect in model on heu and lin
                                     sub2_states=list(c("q1","q2")),
                                     # Panels from left right
                                     stats_for_1sub=F,
                                     stats_for_2sub=F,
                                     #Aesthetics:
                                     par_mar=c(2.6,3,0.3,5),
                                     # Plot margins
                                     sub1_labels="H. t. linaresi",
                                     sub2_labels=list(c("poorly lit","brigthly lit")),
                                     # x axis labels
                                     label_count_col = "ID",
                                     # column by which to determine sample size
                                     yaxs_label="",
                                     # y axis title
                                     y_tck=-0.015,
                                     # Tick size on y-axis
                                     horiz_0.5=F,
                                     # No horizontal line at 0.5
                                     panel_dividers=F,
                                     # No vertical panel dividers
                                     dot_size=0.28,
                                     # Factor to multiply dot sizes with (just aesthetics)
                                     dot_colours=list(c("blue","blue")),
                                     # Dot colours
                                     sub_box_yn=F,
                                     # No Boxplot added
                                     numb_iterations=1000,
                                     # Number iterations for assembling the points
                                     legend_dots=c(),
                                     # No legend dots
                                     border_space=0.49,
                                     # space to left and right side of center that dots can be jittered to
                                     center_space_cat=0.15,
                                     #Empty space in center for reaction norm
                                     reaction_norm_col = "ID",
                                     #Column for reaction norm
                                     reaction_norm_weights=2,
                                     #Reaction norm lines unweighted
                                     reaction_norm_max_trans=0.5,
                                     #Transparency of reaction norm lines
                                     seedi=123,
                                     # Seed
                                     verbose=F
                                     # Do not print comments
)

text(par("usr")[1]+(0.3/par("pin")[1])*diff(par("usr")[1:2]),
     par("usr")[4]-(0.25/par("pin")[2])*diff(par("usr")[3:4]),
     "A",font=2,cex=2.5,
     pos=2,offset=0)

#Scale histograms
max_counts<-max(c(post_hist_within_TLC$counts,
                  post_hist_non_red_only_heu$counts,
                  post_hist_red_only_heu$counts,
                  post_hist_F1$counts,
                  post_hist_BC_H$counts,
                  post_hist_within_HC$counts))
post_hist_within_TLC$counts<-(post_hist_within_TLC$counts/max_counts)*(par("cex")*1.4*par("cin")[2]/par("pin")[1])*diff(par("usr")[1:2])
post_hist_non_red_only_heu$counts<-(post_hist_non_red_only_heu$counts/max_counts)*(par("cex")*1.4*par("cin")[2]/par("pin")[1])*diff(par("usr")[1:2])
post_hist_red_only_heu$counts<-(post_hist_red_only_heu$counts/max_counts)*(par("cex")*1.4*par("cin")[2]/par("pin")[1])*diff(par("usr")[1:2])
post_hist_F1$counts<-(post_hist_F1$counts/max_counts)*(par("cex")*1.4*par("cin")[2]/par("pin")[1])*diff(par("usr")[1:2])
post_hist_BC_H$counts<-(post_hist_BC_H$counts/max_counts)*(par("cex")*1.4*par("cin")[2]/par("pin")[1])*diff(par("usr")[1:2])
post_hist_within_HC$counts<-(post_hist_within_HC$counts/max_counts)*(par("cex")*1.4*par("cin")[2]/par("pin")[1])*diff(par("usr")[1:2])

#We add the Gardner-Altman part
add_gardner_altman(TLC_posterior_cat_1,TLC_posterior_cat_2,difference_posterior_within_TLC,post_hist_within_TLC,at_pos=c(0.25,0.75),add_title=F)

#We add the estimators from figure 2B onto the right side
par(xpd=NA)
segments(1+(par("cex")*1.3*par("cin")[2]/par("pin")[1])*diff(par("usr")[1:2]),mean(TLC_posterior_main),1+(par("cex")*1.7*par("cin")[2]/par("pin")[1])*diff(par("usr")[1:2]),mean(TLC_posterior_main),lwd=3,col="blue",lend=3)
segments(1+(par("cex")*1.3*par("cin")[2]/par("pin")[1])*diff(par("usr")[1:2]),mean(HC_posterior_main),1+(par("cex")*1.7*par("cin")[2]/par("pin")[1])*diff(par("usr")[1:2]),mean(HC_posterior_main),lwd=3,col="red",lend=3)
par(xpd=F)


#backcross to lin, no red

#Calling plot_proportion_stats to produce the panel with the full data
testout_bayes<-plot_proportion_stats(input_data=pref_stat_BC[pref_stat_BC$quantile_low_up_1!="qNA"&pref_stat_BC$redYN=="N",],
                                     # The data set
                                     sum_by_col_plot=c("ID","quantile_low_up_1"),
                                     # Sum by ID for plot
                                     response_col="court_lin_heu_01",
                                     # Response variable
                                     sub2_col="quantile_low_up_1",
                                     # First fixed effect in model on heu and lin
                                     sub2_states=list(c("q1","q2")),
                                     # Panels from left right
                                     stats_for_1sub=F,
                                     stats_for_2sub=F,
                                     #Aesthetics:
                                     par_mar=c(2.6,3,0.3,5),
                                     # Plot margins
                                     sub1_labels="backcross to H. t. linaresi (no red)",
                                     sub2_labels=list(c("poorly lit","brigthly lit")),
                                     # x axis labels
                                     label_count_col = "ID",
                                     # column by which to determine sample size
                                     yaxs_label="",
                                     # y axis title
                                     y_tck=-0.015,
                                     # Tick size on y-axis
                                     horiz_0.5=F,
                                     # No horizontal line at 0.5
                                     panel_dividers=F,
                                     # No vertical panel dividers
                                     dot_size=0.28,
                                     # Factor to multiply dot sizes with (just aesthetics)
                                     dot_colours=list(c("blue","blue")),
                                     # Dot colours
                                     sub_box_yn=F,
                                     # No Boxplot added
                                     numb_iterations=1000,
                                     # Number iterations for assembling the points
                                     legend_dots=c(),
                                     # No legend dots
                                     border_space=0.49,
                                     # space to left and right side of center that dots can be jittered to
                                     center_space_cat=0.15,
                                     #Empty space in center for reaction norm
                                     reaction_norm_col = "ID",
                                     #Column for reaction norm
                                     reaction_norm_weights=2,
                                     #Reaction norm lines unweighted
                                     reaction_norm_max_trans=0.5,
                                     #Transparency of reaction norm lines
                                     seedi=123,
                                     # Seed
                                     verbose=F
                                     # Do not print comments
)

text(par("usr")[1]+(0.3/par("pin")[1])*diff(par("usr")[1:2]),
     par("usr")[4]-(0.25/par("pin")[2])*diff(par("usr")[3:4]),
     "B",font=2,cex=2.5,
     pos=2,offset=0)

#We add the Gardner-Altman part
add_gardner_altman(non_red_posterior_cat_1_only_heu,non_red_posterior_cat_2_only_heu,non_red_difference_posterior_only_heu,post_hist_non_red_only_heu,at_pos=c(0.25,0.75),add_title=F)

#We add the estimators from figure 2B onto the right side
par(xpd=NA)
segments(1+(par("cex")*1.3*par("cin")[2]/par("pin")[1])*diff(par("usr")[1:2]),mean(TLC_posterior_main),1+(par("cex")*1.7*par("cin")[2]/par("pin")[1])*diff(par("usr")[1:2]),mean(TLC_posterior_main),lwd=3,col="blue",lend=3)
segments(1+(par("cex")*1.3*par("cin")[2]/par("pin")[1])*diff(par("usr")[1:2]),mean(HC_posterior_main),1+(par("cex")*1.7*par("cin")[2]/par("pin")[1])*diff(par("usr")[1:2]),mean(HC_posterior_main),lwd=3,col="red",lend=3)
par(xpd=F)


#backcross to lin, red

#Calling plot_proportion_stats to produce the panel with the full data
testout_bayes<-plot_proportion_stats(input_data=pref_stat_BC[pref_stat_BC$quantile_low_up_1!="qNA"&pref_stat_BC$redYN=="Y",],
                                     # The data set
                                     sum_by_col_plot=c("ID","quantile_low_up_1"),
                                     # Sum by ID for plot
                                     response_col="court_lin_heu_01",
                                     # Response variable
                                     sub2_col="quantile_low_up_1",
                                     # First fixed effect in model on heu and lin
                                     sub2_states=list(c("q1","q2")),
                                     # Panels from left right
                                     stats_for_1sub=F,
                                     stats_for_2sub=F,
                                     #Aesthetics:
                                     par_mar=c(2.6,3,0.3,5),
                                     # Plot margins
                                     sub1_labels="backcross to H. t. linaresi (red)",
                                     sub2_labels=list(c("poorly lit","brigthly lit")),
                                     # x axis labels
                                     label_count_col = "ID",
                                     # column by which to determine sample size
                                     yaxs_label="",
                                     # y axis title
                                     y_tck=-0.015,
                                     # Tick size on y-axis
                                     horiz_0.5=F,
                                     # No horizontal line at 0.5
                                     panel_dividers=F,
                                     # No vertical panel dividers
                                     dot_size=0.28,
                                     # Factor to multiply dot sizes with (just aesthetics)
                                     dot_colours=list(c("red","red")),
                                     # Dot colours
                                     sub_box_yn=F,
                                     # No Boxplot added
                                     numb_iterations=1000,
                                     # Number iterations for assembling the points
                                     legend_dots=c(),
                                     # No legend dots
                                     border_space=0.49,
                                     # space to left and right side of center that dots can be jittered to
                                     center_space_cat=0.15,
                                     #Empty space in center for reaction norm
                                     reaction_norm_col = "ID",
                                     #Column for reaction norm
                                     reaction_norm_weights=2,
                                     #Reaction norm lines unweighted
                                     reaction_norm_max_trans=0.5,
                                     #Transparency of reaction norm lines
                                     seedi=123,
                                     # Seed
                                     verbose=F
                                     # Do not print comments
)

text(par("usr")[1]+(0.3/par("pin")[1])*diff(par("usr")[1:2]),
     par("usr")[4]-(0.25/par("pin")[2])*diff(par("usr")[3:4]),
     "C",font=2,cex=2.5,
     pos=2,offset=0)

#We add the Gardner-Altman part
add_gardner_altman(red_posterior_cat_1_only_heu,red_posterior_cat_2_only_heu,red_difference_posterior_only_heu,post_hist_red_only_heu,at_pos=c(0.25,0.75),add_title=F)

#We add the estimators from figure 2B onto the right side
par(xpd=NA)
segments(1+(par("cex")*1.3*par("cin")[2]/par("pin")[1])*diff(par("usr")[1:2]),mean(TLC_posterior_main),1+(par("cex")*1.7*par("cin")[2]/par("pin")[1])*diff(par("usr")[1:2]),mean(TLC_posterior_main),lwd=3,col="blue",lend=3)
segments(1+(par("cex")*1.3*par("cin")[2]/par("pin")[1])*diff(par("usr")[1:2]),mean(HC_posterior_main),1+(par("cex")*1.7*par("cin")[2]/par("pin")[1])*diff(par("usr")[1:2]),mean(HC_posterior_main),lwd=3,col="red",lend=3)
par(xpd=F)


#F1s

#Calling plot_proportion_stats to produce the panel with the full data
testout_bayes<-plot_proportion_stats(input_data=pref_stat_other_hyb[pref_stat_other_hyb$quantile_low_up!="qNA"&pref_stat_other_hyb$type=="F1",],
                                     # The data set
                                     sum_by_col_plot=c("ID","quantile_low_up"),
                                     # Sum by ID for plot
                                     response_col="court_lin_heu_01",
                                     # Response variable
                                     sub2_col="quantile_low_up",
                                     # First fixed effect in model on heu and lin
                                     sub2_states=list(c("q1","q2")),
                                     # Panels from left right
                                     stats_for_1sub=F,
                                     stats_for_2sub=F,
                                     #Aesthetics:
                                     par_mar=c(2.6,3,0.3,5),
                                     # Plot margins
                                     sub1_labels="F1 males",
                                     sub2_labels=list(c("poorly lit","brigthly lit")),
                                     # x axis labels
                                     label_count_col = "ID",
                                     # column by which to determine sample size
                                     yaxs_label="",
                                     # y axis title
                                     y_tck=-0.015,
                                     # Tick size on y-axis
                                     horiz_0.5=F,
                                     # No horizontal line at 0.5
                                     panel_dividers=F,
                                     # No vertical panel dividers
                                     dot_size=0.28,
                                     # Factor to multiply dot sizes with (just aesthetics)
                                     dot_colours=list(c("red","red")),
                                     # Dot colours
                                     sub_box_yn=F,
                                     # No Boxplot added
                                     numb_iterations=1000,
                                     # Number iterations for assembling the points
                                     legend_dots=c(),
                                     # No legend dots
                                     border_space=0.49,
                                     # space to left and right side of center that dots can be jittered to
                                     center_space_cat=0.15,
                                     #Empty space in center for reaction norm
                                     reaction_norm_col = "ID",
                                     #Column for reaction norm
                                     reaction_norm_weights=2,
                                     #Reaction norm lines unweighted
                                     reaction_norm_max_trans=0.5,
                                     #Transparency of reaction norm lines
                                     seedi=123,
                                     # Seed
                                     verbose=F
                                     # Do not print comments
)

text(par("usr")[1]+(0.3/par("pin")[1])*diff(par("usr")[1:2]),
     par("usr")[4]-(0.25/par("pin")[2])*diff(par("usr")[3:4]),
     "D",font=2,cex=2.5,
     pos=2,offset=0)

#We add the Gardner-Altman part
add_gardner_altman(F1_posterior_cat_1,F1_posterior_cat_2,difference_posterior_F1,post_hist_F1,at_pos=c(0.25,0.75),add_title=F)

#We add the estimators from figure 2B onto the right side
par(xpd=NA)
segments(1+(par("cex")*1.3*par("cin")[2]/par("pin")[1])*diff(par("usr")[1:2]),mean(TLC_posterior_main),1+(par("cex")*1.7*par("cin")[2]/par("pin")[1])*diff(par("usr")[1:2]),mean(TLC_posterior_main),lwd=3,col="blue",lend=3)
segments(1+(par("cex")*1.3*par("cin")[2]/par("pin")[1])*diff(par("usr")[1:2]),mean(HC_posterior_main),1+(par("cex")*1.7*par("cin")[2]/par("pin")[1])*diff(par("usr")[1:2]),mean(HC_posterior_main),lwd=3,col="red",lend=3)
par(xpd=F)


#Now backcross to heu

#Calling plot_proportion_stats to produce the panel with the full data
testout_bayes<-plot_proportion_stats(input_data=pref_stat_other_hyb[pref_stat_other_hyb$quantile_low_up!="qNA"&pref_stat_other_hyb$type=="HCx(TLCxHC)",],
                                     # The data set
                                     sum_by_col_plot=c("ID","quantile_low_up"),
                                     # Sum by ID for plot
                                     response_col="court_lin_heu_01",
                                     # Response variable
                                     sub2_col="quantile_low_up",
                                     # First fixed effect in model on heu and lin
                                     sub2_states=list(c("q1","q2")),
                                     # Panels from left right
                                     stats_for_1sub=F,
                                     stats_for_2sub=F,
                                     #Aesthetics:
                                     par_mar=c(2.6,3,0.3,5),
                                     # Plot margins
                                     sub1_labels="backcross to H. heurippa",
                                     sub2_labels=list(c("poorly lit","brigthly lit")),
                                     # x axis labels
                                     label_count_col = "ID",
                                     # column by which to determine sample size
                                     yaxs_label="",
                                     # y axis title
                                     y_tck=-0.015,
                                     # Tick size on y-axis
                                     horiz_0.5=F,
                                     # No horizontal line at 0.5
                                     panel_dividers=F,
                                     # No vertical panel dividers
                                     dot_size=0.28,
                                     # Factor to multiply dot sizes with (just aesthetics)
                                     dot_colours=list(c("red","red")),
                                     # Dot colours
                                     sub_box_yn=F,
                                     # No Boxplot added
                                     numb_iterations=1000,
                                     # Number iterations for assembling the points
                                     legend_dots=c(),
                                     # No legend dots
                                     border_space=0.49,
                                     # space to left and right side of center that dots can be jittered to
                                     center_space_cat=0.15,
                                     #Empty space in center for reaction norm
                                     reaction_norm_col = "ID",
                                     #Column for reaction norm
                                     reaction_norm_weights=2,
                                     #Reaction norm lines unweighted
                                     reaction_norm_max_trans=0.5,
                                     #Transparency of reaction norm lines
                                     seedi=123,
                                     # Seed
                                     verbose=F
                                     # Do not print comments
)

text(par("usr")[1]+(0.3/par("pin")[1])*diff(par("usr")[1:2]),
     par("usr")[4]-(0.25/par("pin")[2])*diff(par("usr")[3:4]),
     "E",font=2,cex=2.5,
     pos=2,offset=0)

#We add the Gardner-Altman part
add_gardner_altman(BC_H_posterior_cat_1,BC_H_posterior_cat_2,difference_posterior_BC_H,post_hist_BC_H,at_pos=c(0.25,0.75),add_title=F)

#We add the estimators from figure 2B onto the right side
par(xpd=NA)
segments(1+(par("cex")*1.3*par("cin")[2]/par("pin")[1])*diff(par("usr")[1:2]),mean(TLC_posterior_main),1+(par("cex")*1.7*par("cin")[2]/par("pin")[1])*diff(par("usr")[1:2]),mean(TLC_posterior_main),lwd=3,col="blue",lend=3)
segments(1+(par("cex")*1.3*par("cin")[2]/par("pin")[1])*diff(par("usr")[1:2]),mean(HC_posterior_main),1+(par("cex")*1.7*par("cin")[2]/par("pin")[1])*diff(par("usr")[1:2]),mean(HC_posterior_main),lwd=3,col="red",lend=3)
par(xpd=F)


#heu

#Calling plot_proportion_stats to produce the panel with the full data
testout_bayes<-plot_proportion_stats(input_data=pref_stat_TLC_HC[pref_stat_TLC_HC$quantile_low_up!="qNA"&pref_stat_TLC_HC$type=="HC",],
                                     # The data set
                                     sum_by_col_plot=c("ID","quantile_low_up"),
                                     # Sum by ID for plot
                                     response_col="court_lin_heu_01",
                                     # Response variable
                                     sub2_col="quantile_low_up",
                                     # First fixed effect in model on heu and lin
                                     sub2_states=list(c("q1","q2")),
                                     # Panels from left right
                                     stats_for_1sub=F,
                                     stats_for_2sub=F,
                                     #Aesthetics:
                                     par_mar=c(2.6,3,0.3,5),
                                     # Plot margins
                                     sub1_labels="H. heurippa",
                                     sub2_labels=list(c("poorly lit","brigthly lit")),
                                     # x axis labels
                                     label_count_col = "ID",
                                     # column by which to determine sample size
                                     yaxs_label="",
                                     # y axis title
                                     y_tck=-0.015,
                                     # Tick size on y-axis
                                     horiz_0.5=F,
                                     # No horizontal line at 0.5
                                     panel_dividers=F,
                                     # No vertical panel dividers
                                     dot_size=0.28,
                                     # Factor to multiply dot sizes with (just aesthetics)
                                     dot_colours=list(c("red","red")),
                                     # Dot colours
                                     sub_box_yn=F,
                                     # No Boxplot added
                                     numb_iterations=1000,
                                     # Number iterations for assembling the points
                                     legend_dots=rev(c(10,20,50,100,200)),
                                     # Legend dots
                                     legend_omi=T,
                                     # Use outer margin for legend
                                     squeeze_legend=0.5,
                                     # Tighten space vertically in legend
                                     legend_info=F,
                                     # Add no additional legend info
                                     border_space=0.49,
                                     # space to left and right side of center that dots can be jittered to
                                     center_space_cat=0.15,
                                     #Empty space in center for reaction norm
                                     reaction_norm_col = "ID",
                                     #Column for reaction norm
                                     reaction_norm_weights=2,
                                     #Reaction norm lines unweighted
                                     reaction_norm_max_trans=0.5,
                                     #Transparency of reaction norm lines
                                     seedi=123,
                                     # Seed
                                     verbose=F
                                     # Do not print comments
)

text(par("usr")[1]+(0.3/par("pin")[1])*diff(par("usr")[1:2]),
     par("usr")[4]-(0.25/par("pin")[2])*diff(par("usr")[3:4]),
     "F",font=2,cex=2.5,
     pos=2,offset=0)

#We add the Gardner-Altman part
add_gardner_altman(HC_posterior_cat_1,HC_posterior_cat_2,difference_posterior_within_HC,post_hist_within_HC,at_pos=c(0.25,0.75))

#We add the estimators from figure 2B onto the right side
par(xpd=NA)
segments(1+(par("cex")*1.3*par("cin")[2]/par("pin")[1])*diff(par("usr")[1:2]),mean(TLC_posterior_main),1+(par("cex")*1.7*par("cin")[2]/par("pin")[1])*diff(par("usr")[1:2]),mean(TLC_posterior_main),lwd=3,col="blue",lend=3)
segments(1+(par("cex")*1.3*par("cin")[2]/par("pin")[1])*diff(par("usr")[1:2]),mean(HC_posterior_main),1+(par("cex")*1.7*par("cin")[2]/par("pin")[1])*diff(par("usr")[1:2]),mean(HC_posterior_main),lwd=3,col="red",lend=3)
par(xpd=F)


mtext(substitute(paste("Proportion interactions with mounted  ",italic('H. heurippa')," female")),
      2,outer=T,line=-0.5,cex=1)

#Close plotting device.
invisible(dev.off())
```

Show png we just made. The resolution in this HTML is unfortunately a bit too low to see the x-axis well enough (but it's well visible in the original PNG). From left to right: *lin*, backcross to *lin* without red, backcross to *lin* with red, F1, backcross to *heu*, *heu*. Left in each panel are data from poorly lit *heu* mounted female, right are data from brightly lit *heu* mounted female.

```
suppl_fig4 <- readPNG("Graphs/supplementary_figure4.png")
grid.raster(suppl_fig4)
```

##### 2.2.6 Supplementary figure 2

We now look at all the interaction terms involving type or presence/absence of red colour in backcross hybrids, respectively. For this, we look at the fully saturated models, `mod1.2` and `mod2.2`. Especially displaying a three-way interaction with two continuous variables is a difficult task! Details on the code can be found in `other_plot_functions.R`.

First, get the two-way interaction terms between log-illuminance around ‘females’ and type (species) from `mod1.2` and presence/absence of red colour on backcross males from `mod2.2`.

```
# As before, we also first find the range of illuminance
# intensities at the respective female
data_range <- range(mod1.2$data$log_light_HC.z)
# We produce 100 values from the range between the two
# extremes
val_to_check <- do.call(seq, as.list(c(data_range, dist(data_range)/99)))
# Run emmeans to get slopes for the effect of illuminance
# around heu 'female' from mod1.2
pure_interaction_heu <- extract_emm_cri(suppressWarnings(suppressMessages(emmeans(mod1.2, 
    ~type | log_light_HC.z, transform = "response", at = list(log_light_HC.z = val_to_check)))))

# Repeat for effect of illuminance around lin 'female' from
# mod1.2
data_range <- range(mod1.2$data$log_light_TLC.z)
val_to_check <- do.call(seq, as.list(c(data_range, dist(data_range)/99)))
pure_interaction_lin <- extract_emm_cri(suppressWarnings(suppressMessages(emmeans(mod1.2, 
    ~type | log_light_TLC.z, transform = "response", at = list(log_light_TLC.z = val_to_check)))))

# Repeat for effect of illuminance around heu 'female' from
# mod2.2
data_range <- range(mod2.2$data$log_light_HC.z)
val_to_check <- do.call(seq, as.list(c(data_range, dist(data_range)/99)))
backcross_interaction_heu <- extract_emm_cri(suppressWarnings(suppressMessages(emmeans(mod2.2, 
    ~redYN | log_light_HC.z, transform = "response", at = list(log_light_HC.z = val_to_check)))))

# Repeat for effect of illuminance around lin 'female' from
# mod2.2
data_range <- range(mod2.2$data$log_light_TLC.z)
val_to_check <- do.call(seq, as.list(c(data_range, dist(data_range)/99)))
backcross_interaction_lin <- extract_emm_cri(suppressWarnings(suppressMessages(emmeans(mod2.2, 
    ~redYN | log_light_TLC.z, transform = "response", at = list(log_light_TLC.z = val_to_check)))))
```

We backtransform every value of log-illuminance, using the scaling parameters safed in the original tables.

```
pure_interaction_heu$log_light_HC <- pure_interaction_heu$log_light_HC.z * 
    attr(pref_stat_TLC_HC$log_light_HC.z, "scaled:scale") + attr(pref_stat_TLC_HC$log_light_HC.z, 
    "scaled:center")
pure_interaction_lin$log_light_TLC <- pure_interaction_lin$log_light_TLC.z * 
    attr(pref_stat_TLC_HC$log_light_TLC.z, "scaled:scale") + 
    attr(pref_stat_TLC_HC$log_light_TLC.z, "scaled:center")
backcross_interaction_heu$log_light_HC <- backcross_interaction_heu$log_light_HC.z * 
    attr(pref_stat_BC$log_light_HC.z, "scaled:scale") + attr(pref_stat_BC$log_light_HC.z, 
    "scaled:center")
backcross_interaction_lin$log_light_TLC <- backcross_interaction_lin$log_light_TLC.z * 
    attr(pref_stat_BC$log_light_TLC.z, "scaled:scale") + attr(pref_stat_BC$log_light_TLC.z, 
    "scaled:center")
```

Now the plot.

```
# Opening plotting device and setting up layout.
png("Graphs/supplementary_figure2.png", width = 1280, height = 2400, 
    res = 300)
layout(matrix(c(rep(c(1, 4), 4), rep(c(2, 5), 4), 3, 6, rep(c(7, 
    8), 4), rep(c(9, 10), 4), 11, 11), byrow = T, ncol = 2))

par(oma = c(0, 3, 0, 4))

# x-dimension of the two-way interaction plots
custom_xlim <- range(c(pure_interaction_heu$log_light_HC, pure_interaction_lin$log_light_TLC, 
    backcross_interaction_heu$log_light_HC, backcross_interaction_lin$log_light_TLC))

# Produce the interaction terms with log-illuminance at heu
# 'female'.
plot_interaction(covariate = "log_light_HC", bay_cov1 = pure_interaction_heu, 
    yaxs_side = 2, par_mar1 = c(0, 0.4, 1.6, 0.4), ylabel = "", 
    added_title_cex = 0.65, add_empty = F, add_xaxs = F, text_inner = F, 
    ylabel_outer = F, ylabel_line = 2, numbers = T, add_letter = "A", 
    letter_central = T, custom_xlim = custom_xlim)

plot_interaction(covariate = "log_light_HC", bay_cov1 = backcross_interaction_heu, 
    added_title = substitute(paste("Illuminance ", italic("H. heurippa"), 
        " [", italic("lux"), "]")), yaxs_side = 2, par_mar1 = c(0, 
        0.4, 1.6, 0.4), ylabel = substitute(paste("                                                            Proportion interactions with mounted  ", 
        italic("H. heurippa"), " female")), added_title_cex = 0.65, 
    add_empty = T, add_xaxs = T, text_inner = F, ylabel_outer = F, 
    ylabel_line = 2, numbers = T, colname_subset = "redYN", sub_names = c("N", 
        "Y"), add_letter = "B", letter_central = T, text_inner_fill = "backcross with vs without red", 
    custom_xlim = custom_xlim)

# Add some dashed lines to separate the panels
par(xpd = NA)
segments(par("usr")[2] + diff(par("usr")[1:2]) * (par("mai")[4]/par("pin")[1]), 
    par("usr")[3] - diff(par("usr")[3:4]) * (par("mai")[1]/par("pin")[2]), 
    -100, par("usr")[3] - diff(par("usr")[3:4]) * (par("mai")[1]/par("pin")[2]), 
    lwd = 1.5, lty = "dashed")
segments(par("usr")[2] + diff(par("usr")[1:2]) * (par("mai")[4]/par("pin")[1]), 
    par("usr")[3] - diff(par("usr")[3:4]) * (par("mai")[1]/par("pin")[2]), 
    par("usr")[2] + diff(par("usr")[1:2]) * (par("mai")[4]/par("pin")[1]), 
    100, lwd = 1.5, lty = "dashed")
par(xpd = F)

# Produce the interaction terms with log-illuminance at lin
# 'female'.
plot_interaction(covariate = "log_light_TLC", bay_cov1 = pure_interaction_lin, 
    yaxs_side = 4, par_mar1 = c(0, 0.4, 1.6, 0.4), ylabel = "", 
    added_title_cex = 0.65, add_empty = F, add_xaxs = F, text_inner = F, 
    ylabel_outer = F, ylabel_line = 2, numbers = T, add_letter = "C", 
    letter_central = T, custom_xlim = custom_xlim)

plot_interaction(covariate = "log_light_TLC", bay_cov1 = backcross_interaction_lin, 
    added_title = substitute(paste("Illuminance ", italic("H. t. linaresi"), 
        " [", italic("lux"), "]")), yaxs_side = 4, par_mar1 = c(0, 
        0.4, 1.6, 0.4), ylabel = "", added_title_cex = 0.65, 
    add_empty = T, add_xaxs = T, text_inner = F, ylabel_outer = F, 
    ylabel_line = 2, numbers = T, colname_subset = "redYN", sub_names = c("N", 
        "Y"), add_letter = "D", letter_central = T, text_inner_fill = "backcross with vs without red", 
    custom_xlim = custom_xlim)

# Add some dashed lines to separate the panels
par(xpd = NA)
segments(par("usr")[1] - diff(par("usr")[1:2]) * (par("mai")[2]/par("pin")[1]), 
    par("usr")[3] - diff(par("usr")[3:4]) * (par("mai")[1]/par("pin")[2]), 
    100, par("usr")[3] - diff(par("usr")[3:4]) * (par("mai")[1]/par("pin")[2]), 
    lwd = 1.5, lty = "dashed")
par(xpd = F)


# Add three-way interactions. Check other_plot_functions.R
# for details (this is quite a complex calculation)
interact3(col_range = c(0.2, 0.8))

# Close plotting device.
invisible(dev.off())
```

Show png we just made

```
suppl_fig2 <- readPNG("Graphs/supplementary_figure2.png")
grid.raster(suppl_fig2)
```

The dotted contours are weighted by the preference for *own*heu\* female type (i.e. the more red the more weight). We can see that there is a slight tendency for a three-way interaction, meaning that the effect of a bright *heu* increasing preference for the own type is stronger, if the *lin* ‘female’ is not bright.

##### 2.2.7 Supplementary figure 3

In this figure, we take a closer look at the raw log-illuminance data and “local preferences” in the “light space”. This figure should visualize were in the “illuminance space” our data is distributed in and in which regions of this space certain preferences materialize.

We now plot the raw data and local preferences. As before, details can be found in `other_plot_functions.R`.

```
# Opening plotting device and setting up layout.
png("Graphs/supplementary_figure3.png", res = 300, height = 2000, 
    width = 1270)
layout(matrix(1:8, byrow = T, ncol = 2))
par(oma = c(1.5, 4, 0.05, 4.3))
par(mar = c(1.5, 0.5, 0.05, 0.7))

# Plot raw lin data with Kernel densities
light_plot(whole_dataset = pref_stat_TLC_HC, subset_TF = pref_stat_TLC_HC$type == 
    "TLC", ncat = 6, letter = "A")
mtext(substitute(italic("H. t. linaresi")), 2, line = 2, cex = 0.65)

# Plot local preferences of lin males
light_plot_pref(whole_dataset = pref_stat_TLC_HC, subset_TF = pref_stat_TLC_HC$type == 
    "TLC", number_bins = 15, yat = 4, transparency = F, letter = "E")

# Plot raw heu data with Kernel densities
light_plot(whole_dataset = pref_stat_TLC_HC, subset_TF = pref_stat_TLC_HC$type == 
    "HC", ncat = 6, letter = "B")
mtext(substitute(italic("H. heurippa")), 2, line = 2, cex = 0.65)

# Plot local preferences of heu males
light_plot_pref(whole_dataset = pref_stat_TLC_HC, subset_TF = pref_stat_TLC_HC$type == 
    "HC", number_bins = 15, yat = 4, transparency = F, letter = "F")

# Plot raw non-red backcross data with Kernel densities
light_plot(whole_dataset = pref_stat_BC, subset_TF = pref_stat_BC$redYN == 
    "N", ncat = 6, letter = "C")
mtext(substitute(paste("Non-red BC to ", italic("H. t. linaresi"))), 
    2, line = 2, cex = 0.65)

# Plot local preferences of non-red backcross males
light_plot_pref(whole_dataset = pref_stat_BC, subset_TF = pref_stat_BC$redYN == 
    "N", number_bins = 15, yat = 4, transparency = F, letter = "G")

# Add colour scale
add_scale(xprop_start = 0.235, xprop_end = 0.355, col_range = c(0, 
    1))

# Plot raw red backcross data with Kernel densities
light_plot(whole_dataset = pref_stat_BC, subset_TF = pref_stat_BC$redYN == 
    "Y", ncat = 6, letter = "D")
mtext(substitute(paste("Red BC to ", italic("H. t. linaresi"))), 
    2, line = 2, cex = 0.65)

# Plot local preferences of red backcross males
light_plot_pref(whole_dataset = pref_stat_BC, subset_TF = pref_stat_BC$redYN == 
    "Y", number_bins = 15, yat = 4, transparency = F, letter = "H")

# Add axes titles
mtext(substitute(paste("Illuminance at mounted ", italic("H. heurippa"), 
    " female")), 1, outer = T, line = 0.5, cex = 0.65)
mtext(substitute(paste("Illuminance at mounted ", italic(" H. t. linaresi"), 
    " female")), 2, outer = T, line = 3, cex = 0.65)

# Close plotting device.
invisible(dev.off())
```

Show png we just made

```
suppl_fig3 <- readPNG("Graphs/supplementary_figure3.png")
grid.raster(suppl_fig3)
```

We can see the same trends that the model found. The local preferences show how the model got “tilted” into certain directions. We have to mind though that some of these “local preferences” result from just few data points, so we have to check the raw data graphs on the left to see how “important” each local preference on the right is.

### 3 Tetrad experiments with alive females

Recode contrasts again to old setting (which was `contr.treatment`).

```
options(contrasts = c("contr.treatment", "contr.poly"))
```

Read in tetrad data.

```
comb_tetrad <- read.table("Data/tetrad_data.txt", header = T)
```

The columns indicate:

- **counts**: number of experiments where the combination of conditions in the other columns was matched.
- **region\_heu**: wild population from which the *heu* males and females used in the experiment originated from. *bva* = Buenavista; *lej* = Lejanias; *sma* = Santa Maria; *mix* = mixed stock combined from genepools of all previous regions
- **female**: female species involved in mating. *lin* or *heu*
- **male**: male species involved in mating. *lin* or *heu*
- **combo**: mating pair with female\_male, e.g. *lin\_heu* = *lin* female mated by *heu* male
- **same\_type**: *same* = conspecific mating; *diff* = heterospecific mating

#### 3.1 Modelling

In our model, we look at how mating count depends on mating pair. The model is of family Poisson, but with the divisions done later actually represents something similar to a multinomial model (sensu Royle, J. Andrew, and Robert M. Dorazio. Hierarchical modeling and inference in ecology: the analysis of data from populations, metapopulations and communities. Elsevier, 2008.).

We use the `brms` package as a R interface for Stan.

```
# Model
tetr_mod <- suppressMessages(brm(counts ~ combo, data = comb_tetrad, 
    family = "poisson", chains = 5, iter = 3e+05, warmup = 30000, 
    refresh = 0, silent = TRUE, seed = 46))
```

#### 3.2 Posteriors

We extract the posteriors:

```
posteriors <- posterior_samples(tetr_mod)
```

We calculate the posteriors for the counts for each mating pair. Note that here in all the following calculations, model coefficients always have to be backtransformed (exponentially) to get understandable count data.

```
post_heu_heu <- exp(posteriors[, c("b_Intercept")])
post_heu_lin <- exp(rowSums(posteriors[, c("b_Intercept", "b_comboheu_lin")]))
post_lin_heu <- exp(rowSums(posteriors[, c("b_Intercept", "b_combolin_heu")]))
post_lin_lin <- exp(rowSums(posteriors[, c("b_Intercept", "b_combolin_lin")]))
```

#### 3.3 Plot, CrIs hypothesis tests

We calculate the posterior for the proportion of each mating pair.

```
counts_all <- post_heu_heu + post_heu_lin + post_lin_heu + post_lin_lin
prop_heu_heu <- post_heu_heu/counts_all
prop_heu_lin <- post_heu_lin/counts_all
prop_lin_heu <- post_lin_heu/counts_all
prop_lin_lin <- post_lin_lin/counts_all
```

We calculate the mean, 2.5% and 97.5% quantile of those posteriors. Also here we use the mean (and not the median), since all means will add up to 1, but all medians won’t.

```
quant_heu_heu <- c(mean = mean(prop_heu_heu), quantile(prop_heu_heu, 
    probs = c(0.025, 0.975)))[c(2, 1, 3)]
quant_heu_lin <- c(mean = mean(prop_heu_lin), quantile(prop_heu_lin, 
    probs = c(0.025, 0.975)))[c(2, 1, 3)]
quant_lin_heu <- c(mean = mean(prop_lin_heu), quantile(prop_lin_heu, 
    probs = c(0.025, 0.975)))[c(2, 1, 3)]
quant_lin_lin <- c(mean = mean(prop_lin_lin), quantile(prop_lin_lin, 
    probs = c(0.025, 0.975)))[c(2, 1, 3)]
```

For each type of males (*heu* and *lin*), we also calculate a preference score comparable to previous analyses.

```
pref_heu_males <- post_heu_heu/(post_heu_heu + post_lin_heu)
pref_lin_males <- post_heu_lin/(post_heu_lin + post_lin_lin)
```

We calculate the mean, 2.5% and 97.5% quantile of those posteriors.

```
quant_pref_heu <- c(mean(pref_heu_males), quantile(pref_heu_males, 
    probs = c(0.025, 0.975)))[c(2, 1, 3)]
quant_pref_lin <- c(mean(pref_lin_males), quantile(pref_lin_males, 
    probs = c(0.025, 0.975)))[c(2, 1, 3)]
```

###### 3.3.0.1 CrI

Let’s look at the CrIs in a table.

First the preferences of males of the two types.

```
quant_tab_pref <- data.frame(c("H. t. linaresi males", "H. heurippa males"), 
    round(c(quant_pref_lin[1], quant_pref_heu[1]), 3), round(c(quant_pref_lin[2], 
        quant_pref_heu[2]), 3), round(c(quant_pref_lin[3], quant_pref_heu[3]), 
        3))
names(quant_tab_pref) <- c("Pair", "2.5%", "mean", "97.5%")
quant_tab_pref
```

Now the proportion of each mating pair.

```
quant_tab_pair <- data.frame(c("lin male + heu female", "lin male + lin female", 
    "heu male + heu female", "heu male + lin female"), round(c(quant_heu_lin[1], 
    quant_lin_lin[1], quant_heu_heu[1], quant_lin_heu[1]), 3), 
    round(c(quant_heu_lin[2], quant_lin_lin[2], quant_heu_heu[2], 
        quant_lin_heu[2]), 3), round(c(quant_heu_lin[3], quant_lin_lin[3], 
        quant_heu_heu[3], quant_lin_heu[3]), 3))
names(quant_tab_pair) <- c("Type", "2.5%", "mean", "97.5%")
quant_tab_pair
```

###### 3.3.0.2 Numeric predictions given mounted female data

Given our results with the mounted females, we can make some numeric predictions for the tetrad experiments. Given that we know of *heu* being more vigorous, we can use the previously measured preferences under different light environments and predict what we expect as results from the tetrad experiments.

This is the observed proportion of matings involving a *heu* male:

```
(prop_mating_heu_male <- quant_heu_heu[2] + quant_lin_heu[2])
```

```
##      mean 
## 0.6743626
```

Here is what we predict for the overall visual preferences we measured:

```
pred_heu_lin_all <- (1 - prop_mating_heu_male) * mean(TLC_posterior_main)
pred_lin_lin_all <- (1 - prop_mating_heu_male) * (1 - mean(TLC_posterior_main))
pred_heu_heu_all <- prop_mating_heu_male * mean(HC_posterior_main)
pred_lin_heu_all <- prop_mating_heu_male * (1 - mean(HC_posterior_main))
```

For poorly lit *heu* female (from the categorical model)

```
pred_heu_lin_dark <- (1 - prop_mating_heu_male) * mean(TLC_posterior_cat_1)
pred_lin_lin_dark <- (1 - prop_mating_heu_male) * (1 - mean(TLC_posterior_cat_1))
pred_heu_heu_dark <- prop_mating_heu_male * mean(HC_posterior_cat_1)
pred_lin_heu_dark <- prop_mating_heu_male * (1 - mean(HC_posterior_cat_1))
```

For brightly lit *heu* female (from the categorical model)

```
pred_heu_lin_bright <- (1 - prop_mating_heu_male) * mean(TLC_posterior_cat_2)
pred_lin_lin_bright <- (1 - prop_mating_heu_male) * (1 - mean(TLC_posterior_cat_2))
pred_heu_heu_bright <- prop_mating_heu_male * mean(HC_posterior_cat_2)
pred_lin_heu_bright <- prop_mating_heu_male * (1 - mean(HC_posterior_cat_2))
```

We make a nice table with CI/CrIs of real data and of estimates. We take the estimated means from the `binom.bayes` function, which are always almost exactly the same as the given probability. When given the default prior, the posterior mean is always defined as `(x + 0.5)/(n + 1)`, where `x` is successes and `n` is trials. The result from this returns a value **very** similar to the “true” estimated proportion of successes.

```
tetr_est <- rbind(paste0(quant_tab_pair$mean, " [", paste0(quant_tab_pair$`2.5%`, 
    "; ", quant_tab_pair$`97.5%`), "]"), sapply(c(pred_heu_lin_all, 
    pred_lin_lin_all, pred_heu_heu_all, pred_lin_heu_all), function(x) c(paste0(round(binom.bayes(x * 
    89, 89, type = "central")[6], 3), " [", paste(round(binom.bayes(x * 
    89, 89, type = "central")[7:8], 3), collapse = "; "), "]"))), 
    sapply(c(pred_heu_lin_dark, pred_lin_lin_dark, pred_heu_heu_dark, 
        pred_lin_heu_dark), function(x) c(paste0(round(binom.bayes(x * 
        89, 89, type = "central")[6], 3), " [", paste(round(binom.bayes(x * 
        89, 89, type = "central")[7:8], 3), collapse = "; "), 
        "]"))), sapply(c(pred_heu_lin_bright, pred_lin_lin_bright, 
        pred_heu_heu_bright, pred_lin_heu_bright), function(x) c(paste0(round(binom.bayes(x * 
        89, 89, type = "central")[6], 3), " [", paste(round(binom.bayes(x * 
        89, 89, type = "central")[7:8], 3), collapse = "; "), 
        "]"))))
rownames(tetr_est) <- c("true experiment", "estimate all data", 
    "estimate dark", "estimate bright")
colnames(tetr_est) <- c("Heu_female+Lin_male", "Lin_female+Lin_male", 
    "Heu_female+Heu_male", "Lin_female+Heu_male")
tetr_est <- as.data.frame(tetr_est)
tetr_est
```

###### 3.3.0.3 Figure 3

We visualize these proportions in a plot (see again `other_plot_functions.R` for details).

```
pdf("Graphs/figure3.pdf", width = 4, height = 5)
tetrad_fig(prop_heu_lin = prop_heu_lin, prop_lin_lin = prop_lin_lin, 
    prop_heu_heu = prop_heu_heu, prop_lin_heu = prop_lin_heu, 
    quant_heu_lin = quant_heu_lin, quant_lin_lin = quant_lin_lin, 
    quant_heu_heu = quant_heu_heu, quant_lin_heu = quant_lin_heu, 
    datasheet = comb_tetrad, estimations = matrix(as.numeric(sapply(c(pred_heu_lin_bright, 
        pred_lin_lin_bright, pred_heu_heu_bright, pred_lin_heu_bright), 
        function(x) binom.bayes(x * 89, 89, type = "central")[6:8])), 
        ncol = 4))
invisible(dev.off())
```

We convert the pdf to png, so we have both file types. Unfortunately, the purple dot showing the estimator in the center of each CrI for the prediction derived from the mounted female data is not showing up after the conversion. It is there in the pdf though!

```
pdf_convert("Graphs/figure3.pdf", filenames = "Graphs/figure3.png", 
    dpi = 300)
```

```
## Converting page 1 to Graphs/figure3.png... done!
```

```
## [1] "Graphs/figure3.png"
```

Show png we just made

```
fig3 <- readPNG("Graphs/figure3.png")
grid.raster(fig3)
```

###### 3.3.0.4 Hypothesis testing

We can now test how probable it is that males either of the two types of butterflies mate assortatively, as well as whether these two types of butterflies mate assortatively overall. We could do this simply by comparing posteriors “manually”, but we can also use the handy `hypothesis()` function from `brms`, which gives us some nice additional info.

We construct calculus strings for that. We have to add all model coefficients up with the same logic as we added all posteriors up before.

```
lin_lin <- "exp(Intercept+combolin_lin)"

heu_heu <- "exp(Intercept)"

heu_lin <- "exp(Intercept+comboheu_lin)"

lin_heu <- "exp(Intercept+combolin_heu)"
```

Now we test the hypothesis that *lin* males mate assortatively (this test is explicitely looking if the “preference score” of *lin* males is below 0.5):

```
print(hypothesis(tetr_mod, paste0(lin_lin, ">", heu_lin)))
```

```
## Hypothesis Tests for class b:
##                 Hypothesis Estimate Est.Error CI.Lower CI.Upper Evid.Ratio
## 1 (exp(Intercept+co... > 0    -0.25      1.34    -2.46     1.95       0.74
##   Post.Prob Star
## 1      0.43     
## ---
## 'CI': 90%-CI for one-sided and 95%-CI for two-sided hypotheses.
## '*': For one-sided hypotheses, the posterior probability exceeds 95%;
## for two-sided hypotheses, the value tested against lies outside the 95%-CI.
## Posterior probabilities of point hypotheses assume equal prior probabilities.
```

Here just two examples how we can easily calculate the posterior probability (`Post.Prob`) ourselves “manually” as well, with:

```
# either
print(sum(prop_lin_lin > prop_heu_lin)/length(prop_heu_lin))
```

```
## [1] 0.4251985
```

```
# ...or
print(sum(pref_lin_males < 0.5)/length(pref_lin_males))
```

```
## [1] 0.4251985
```

We see that there is weak support for assortative mating in *lin* males. In fact, they mated slightly more often with the *heu* female. Mind though that the sample size is relatively small, because *lin* males got to mate less often than *heu* males. ~43% of the posterior distribution lies left of the 0.5 line.

We test whether *heu* males mate assortatively (this test is explicitely looking if the “preference score” of *heu* males is above 0.5):

```
print(hypothesis(tetr_mod, paste0(heu_heu, ">", lin_heu)))
```

```
## Hypothesis Tests for class b:
##                 Hypothesis Estimate Est.Error CI.Lower CI.Upper Evid.Ratio
## 1 (exp(Intercept))-... > 0        3      1.94    -0.16     6.21      15.99
##   Post.Prob Star
## 1      0.94     
## ---
## 'CI': 90%-CI for one-sided and 95%-CI for two-sided hypotheses.
## '*': For one-sided hypotheses, the posterior probability exceeds 95%;
## for two-sided hypotheses, the value tested against lies outside the 95%-CI.
## Posterior probabilities of point hypotheses assume equal prior probabilities.
```

We find high support for *heu* males mating more frequently with females of their own type. ~94% of the posterior distribution of “preferences” lies right of the 0.5 line.

Now we test whether there is overall assortative mating between the two types (basically combining the two previous hypothesis tests):

```
print(hypothesis(tetr_mod, paste0(heu_heu, "+", lin_lin, ">", 
    lin_heu, "+", heu_lin)))
```

```
## Hypothesis Tests for class b:
##                 Hypothesis Estimate Est.Error CI.Lower CI.Upper Evid.Ratio
## 1 (exp(Intercept)+e... > 0     2.75      2.36    -1.11     6.65       7.31
##   Post.Prob Star
## 1      0.88     
## ---
## 'CI': 90%-CI for one-sided and 95%-CI for two-sided hypotheses.
## '*': For one-sided hypotheses, the posterior probability exceeds 95%;
## for two-sided hypotheses, the value tested against lies outside the 95%-CI.
## Posterior probabilities of point hypotheses assume equal prior probabilities.
```

There is quite high support that conspecific pairs occur more frequently than heterospecific pairs. We can visualize this hypothesis test by plotting the proportional difference between conspecific and heterospecific mating calculated as \((counts\_c-counts\_h)/(counts\_c+counts\_h)\), where *c* stands for conspecific and *h* for heterospecific. The proportion of datapoints left of the dashed line equals the posterior probability returned by the `hypothesis` function.

```
layout(1)
par(xpd = F)
par(mar = c(4, 0.4, 0.2, 0.4))
hist(((post_heu_heu + post_lin_lin) - (post_heu_lin + post_lin_heu))/((post_heu_heu + 
    post_lin_lin) + (post_heu_lin + post_lin_heu)), main = "", 
    ylab = "", yaxt = "n", xlab = "", breaks = 100, col = "lightgrey")
mtext("Proportional Difference between Con- and Heterospec. Mating", 
    1, line = 2.7, cex = 1)
abline(v = 0, lwd = 3, lty = "dashed")
```

———END———

### 4 Session Info

To get R Markdown to produce a HTML without problems, install Pandoc as described under this link: https://github.com/rstudio/rstudio/issues/4462

Also, I installed the libraries `xcolor` and `formatR` for nicer Markdown output (not sure if they were actually necessary in the end).

For the overall format, I installed the package `rmdformats` and used the theme `readthedown`.

Session info:

```
sessionInfo()
```

```
## R version 3.6.1 (2019-07-05)
## Platform: x86_64-w64-mingw32/x64 (64-bit)
## Running under: Windows 10 x64 (build 18363)
## 
## Matrix products: default
## 
## locale:
## [1] LC_COLLATE=English_Germany.1252  LC_CTYPE=English_Germany.1252   
## [3] LC_MONETARY=English_Germany.1252 LC_NUMERIC=C                    
## [5] LC_TIME=English_Germany.1252    
## 
## attached base packages:
## [1] grid      stats     graphics  grDevices utils     datasets  methods  
## [8] base     
## 
## other attached packages:
##  [1] truncnorm_1.0-8    arm_1.10-1         binom_1.1-1        ENmisc_1.2-7      
##  [5] RColorBrewer_1.1-2 vcd_1.4-7          MASS_7.3-51.5      ggtern_3.3.0      
##  [9] png_0.1-7          pdftools_2.3.1     weights_1.0.1      mice_3.8.0        
## [13] gdata_2.18.0       Hmisc_4.4-0        ggplot2_3.3.0      Formula_1.2-3     
## [17] survival_3.1-12    lattice_0.20-38    emmeans_1.4.6      lme4_1.1-23       
## [21] Matrix_1.2-18      brms_2.13.0        Rcpp_1.0.2         rmdformats_0.3.7  
## [25] knitr_1.28        
## 
## loaded via a namespace (and not attached):
##   [1] backports_1.1.4      plyr_1.8.6           igraph_1.2.5        
##   [4] splines_3.6.1        crosstalk_1.1.0.1    TH.data_1.0-10      
##   [7] rstantools_2.0.0     inline_0.3.15        digest_0.6.25       
##  [10] htmltools_0.4.0      rsconnect_0.8.16     fansi_0.4.1         
##  [13] magrittr_1.5         checkmate_2.0.0      cluster_2.1.0       
##  [16] bayesm_3.1-4         matrixStats_0.56.0   xts_0.12-0          
##  [19] sandwich_2.5-1       askpass_1.1          prettyunits_1.1.1   
##  [22] jpeg_0.1-8.1         colorspace_1.4-1     xfun_0.13           
##  [25] dplyr_0.8.3          jsonlite_1.6.1       callr_3.4.3         
##  [28] crayon_1.3.4         zoo_1.8-7            glue_1.4.0          
##  [31] gtable_0.3.0         compositions_1.40-5  pkgbuild_1.0.6      
##  [34] rstan_2.19.3         DEoptimR_1.0-8       abind_1.4-5         
##  [37] scales_1.0.0         mvtnorm_1.1-0        qpdf_1.1            
##  [40] miniUI_0.1.1.1       xtable_1.8-4         htmlTable_1.13.3    
##  [43] foreign_0.8-76       latex2exp_0.4.0      stats4_3.6.1        
##  [46] StanHeaders_2.21.0-1 DT_0.13              htmlwidgets_1.5.1   
##  [49] threejs_0.3.3        acepack_1.4.1        ellipsis_0.3.0      
##  [52] pkgconfig_2.0.3      loo_2.2.0            nnet_7.3-13         
##  [55] labeling_0.3         tidyselect_0.2.5     rlang_0.4.5         
##  [58] reshape2_1.4.4       later_1.0.0          munsell_0.5.0       
##  [61] tools_3.6.1          cli_2.0.2            generics_0.0.2      
##  [64] broom_0.5.6          ggridges_0.5.2       evaluate_0.14       
##  [67] stringr_1.4.0        fastmap_1.0.1        yaml_2.2.1          
##  [70] processx_3.4.2       robustbase_0.93-6    purrr_0.3.4         
##  [73] nlme_3.1-140         mime_0.9             formatR_1.7         
##  [76] compiler_3.6.1       bayesplot_1.7.2      shinythemes_1.1.2   
##  [79] rstudioapi_0.11      tibble_3.0.1         statmod_1.4.34      
##  [82] stringi_1.4.3        ps_1.3.2             Brobdingnag_1.2-6   
##  [85] nloptr_1.2.2.1       markdown_1.1         shinyjs_1.1         
##  [88] tensorA_0.36.1       vctrs_0.2.4          pillar_1.4.3        
##  [91] lifecycle_0.2.0      lmtest_0.9-37        bridgesampling_1.0-0
##  [94] estimability_1.3     data.table_1.12.8    httpuv_1.5.2        
##  [97] R6_2.4.1             latticeExtra_0.6-29  bookdown_0.18       
## [100] promises_1.1.0       gridExtra_2.3        codetools_0.2-16    
## [103] boot_1.3-24          colourpicker_1.0     gtools_3.8.2        
## [106] assertthat_0.2.1     proto_1.0.0          withr_2.2.0         
## [109] shinystan_2.5.0      multcomp_1.4-13      parallel_3.6.1      
## [112] rpart_4.1-15         tidyr_1.0.0          coda_0.19-3         
## [115] minqa_1.2.4          rmarkdown_2.1        shiny_1.4.0.2       
## [118] base64enc_0.1-3      dygraphs_1.1.1.6
```
